# Suppression of non-canonical autophagy induces endothelial and cardiac dysfunction

**DOI:** 10.64898/2025.12.17.695030

**Authors:** Joëlle Magné, Suresh Poudel, Rona J. Strawbridge, Luigi Mari, Maria Sabater-Lleal, Clifford S. Guy, Thomas Confer, Melissa Johnson, Mia Panlilio, Jaison John, Nicolas Denans, Pashupati Mishra, Joey Ward, Aaron Pitre, Aaron Taylor, Terho Lehtimäki, Olli T. Raitakari, Yadav Sapkota, Abubakar Wani, Halime Kalkavan, Slim Azouzi, Bérengère Koehl, Brant E Isakson, Khaled Khairy, Douglas R Green

## Abstract

**Background:** While roles for canonical autophagy in the pathophysiology of cardiovascular disease have been established, we have limited understanding of the non-canonical functions of autophagy proteins in this context. LC3-asssociated endocytosis (LANDO) is a novel non-canonical function of autophagy proteins, in which LC3 (microtubule-associated protein light chain 3) is conjugated to early endosome membranes using a portion of the canonical autophagy machinery, and functions in the endocytic recycling of several plasma membrane proteins. Here we ask whether perturbation of LANDO can promote cardiovascular pathogenesis.

**Methods:** Cardiac and endothelial functions were assessed by echocardiography and flow-mediated dilatation in mice lacking Rubicon (*Rubcn^-/-^*) or the WD domain of ATG16L1 (*Atg16l1*^ΔWDki^), two known effectors of LANDO. Mice with conditional depletion of Rubicon in the endothelial, myeloid and cardiomyocyte compartments were used as well. Three-dimensional murine cardiac vasculature leakiness was investigated by light sheet fluorescence microcopy. Endothelial activation induced by shear stress was characterized in vitro in primary endothelial cells isolated from murine lungs and human aortic endothelial cells. Associations between genetically predicted expression of candidate genes involved in LANDO and human cardiovascular parameters were studied in the Young Finns Study and the UK Biobank.

**Results:** Compared to littermate controls, young *Rubcn^-/-^* and *Atg16l1*^ΔWDki^ mice showed a decrease in cardiac and endothelial functions, as did mice with endothelium-specific deficiency. VEGFR2 recycling to the plasma membrane and nitric oxide pathway during shear stress were disrupted in LANDO-deficient primary murine and human endothelial cells. Proteomic analysis in primary human aortic endothelial cells revealed an upregulation of intracellular hemoglobin subunit alpha (Hb-⍺) upon shear stress, which was blunted when *RUBCN* was ablated. Genetic expression studies uncovered several candidate genes related to LANDO that correlated with cardiovascular parameters. These included the retromer complex subunit *VPS29*, disruption of which decreased Hb-⍺ expression levels in human endothelial cells.

**Conclusions:** Our data support a pivotal role of non-canonical functions of autophagy proteins in recycling VEGFR2 upon shear stress activation in endothelial cells together with Hb-⍺ expression that may contribute to the etiology of cardiovascular diseases.

## Introduction

Macroautophagy, (herein, autophagy), is a highly conserved degradative and recycling process essential for the preservation of cellular homeostasis during metabolic adaptation, organismal development, immune or stress responses^1^. Cytoplasmic materials such as misfolded proteins, aggregates or damaged organelles are engulfed in a double membrane vesicle called the autophagosome, which ultimately fuses with lysosomes for cargo degradation^2^. Pharmacological interventions along with the generation of gene-edited organisms impairing autophagy revealed the role of autophagy-related genes (ATG) in the pathogenesis of cardiovascular (CV) disorders including cardiac hypertrophy, ischemic heart diseases, chemotherapy-induced cardiotoxicity, endothelial dysfunction, atherosclerosis and heart failure^3–6^. While autophagy is downregulated during aging and promotion of autophagic activity may represent an attractive therapeutic approach for age-associated pathologies, whether autophagy deficiency is causal, primarily contributive or secondarily beneficial to CV disease (CVD) progression remains unclear^7^.

Among others, our lab provided evidence supporting the involvement of several autophagy-related proteins in non-autophagic processes such as phagocytosis, endocytosis, entosis and micropinocytosis^8^. Microtubule-associated protein light chain 3 (LC3), the lipidation of which is widely used as an hallmark of autophagosome formation, is also recruited to single membrane vesicles (i.e. phagosomes and endosomes), in pathways called LC3-associated phagocytosis (LAP) and LC3-associated endocytosis (LANDO)^9,10^. In the myeloid compartment, LAP represents an efficient response to infection and efferocytosis of apoptotic cells and promotes tumor immune tolerance^11^. Initially observed in microglial cells, LANDO is involved in the recycling of receptors of β-amyloid uptake and prevents Alzheimer’s disease and neurodegeneration^12^. Herein, we refer to LAP and LANDO as “non-canonical autophagy” recognizing that these are not autophagy pathways, per se, but rather non-canonical functions of proteins involved in canonical autophagy.

Over the past 15 years, studies suggested that Rubicon (Run domain Beclin-1-Interacting and Cystein-rich domain Containing protein), one of the few endogenous negative regulators of canonical autophagy, is a crucial molecule for LAP and LANDO^10,13^. Contradicting data exists concerning Rubicon’s role in CVD. Whole-body and specific genetic deletion of Rubicon in mice extended their life span and was associated with protection against lipopolysaccharide-induced stroke, cardiac pressure overload, doxorubicin-induced cardiotoxicity and cardiac ischemic reperfusion^14^. Recent human studies reported that while plasma levels of Rubicon were inversely associated with acute coronary syndrome, increased circulating levels of Rubicon were also independently associated with an increased risk for myocardial infarction^15,16^. Therefore, there is a need to uncouple the implications of other molecules involved in LAP and LANDO in CV pathogenesis and to delineate the relevance of non-canonical functions of autophagy proteins in a cell type and patho-physiological context.

The WD40 domain of ATG16L1, a core protein of the ATG12-ATG5-ATG16L1 E3 ligase complex that lipidates LC3 is essential for LAP and LANDO but dispensable for canonical autophagy^17,18^. Although deficiency in the WD40 domain of ATG16L1 induced neuroinflammation in an Alzheimer’s mouse model, and an increased sensitivity to influenza A virus and an aggravated cerulein-induced acute pancreatitis^19–21^, its role in the initiation and development of CVD remains completely unknown.

In the present study, we sought to assess early markers of CV functions in mice deficient for Rubicon or the WD40 domain of ATG16L1, the only two non-canonical autophagy effectors known so far.

## Methods

Detailed methods are provided in the Supplementary Material.

### Experimental Animals

All animal experiments were approved by St. Jude Institutional Animal Care and Use Committee in accordance with the Guide for the Care and Use of Animals and ARRIVE (Animal Research: Reporting of InVivo Experiments) guidelines. Deletion of the WD domain of ATG16L1 in Atg16l1*^ΔWD^ ^ki^* mice was produced and maintained on a C57BL/6N background as previously described^19,20^. *Rubcn^-/-^* and *Rubcn^fl/fl^* mice were previously generated in C57BL/6N background^11^. C57BL/6N wild-type, *Cdh5-Cre^Tg+^*, *Myh6-Cre^Tg+^*and *LysM-Cre^Tg+^* mice were purchased from Jackson Laboratory, bred and maintained in our facilities. All experiments were performed on 2-month-old and/or 6-month-old mice on a male sex cohort (unless otherwise noted).

### Non-invasive measurement of endothelial function In Vivo

On mice maintained on 1.0–1.5% isoflurane, vascular function in vivo was measured as flow-mediated dilation in the femoral artery with a Vevo 3100 with a 29 to 71 MHz linear array microscan transducer (VisualSonics) as described previously^22,23^

### Echocardiography

Mice were anesthetized with 1.0–1.5% isoflurane for evaluation of diastolic and systolic function. The echocardiography technician was blinded to animal treatment groups and genotypes.

### Light sheet microscopy of cardiac vasculature leakiness

For light sheet experiments, mice received intravenous (i.v) tail vein injection 10min before sacrifice with anti-mouse CD31 (clone MEC13.3) conjugated with AlexaFluor (AF)-647 and with 70kDa-Dextran conjugated with Tetramethylrhodamine. After sacrifice, heart clearing and immunostaining were performed according to the iDisco+ protocol^24^. Clearing involved dehydration through a methanol series, delipidation in dichloromethane, and rehydration. Immunostaining was performed for four days with primary antibodies against CD31 directly labeled with AlexaFluor647. Refractive index matching used dehydration in methanol followed by ethyl cinnamate. Imaging used a Zeiss Light Sheet 7 microscope with 5x/0.1 NA light sheet forming objectives and a 5x/0.16 NA detection objective along with dye-specific lasers and filters. Volumetric tiles were stitched using Imaris Stitcher and visualized using Imaris (Bitplane).

### Data and Material Availability

All data associated with this study are presented in the article or Supplemental Material. RNA sequencing data have been uploaded to the National Center for Biotechnology Information Gene Expression Omnibus repository.

### Statistical Analysis

For animal and in vitro studies, statistical analyses were performed using unpaired Student *t* test; or 2-way ANOVA where a Tukey post hoc test was applied (unless otherwise noted). Data were plotted using GraphPad Prism software Version 10.4.1. All bar graphs and line graphs are represented by mean ± SEM, and difference were considered statistically significant when *P*<0.05). The samples size is described in all figure legends.

Genetic association analyses were conducted on the UK Biobank (UKB) using Plink 1.9 by logistic regression for dichotomic traits, and linear regression for quantitative traits, assuming an additive genetic model and adjusting for age, sex, population structure and genotyping chip. For the Young Finns Study, the genetic association analyses were using PLINK 2 using additive models adjusted for age, sex and first five principal components of the genetic data.

## Results

### Deficiency in Rubicon or the WD domain of ATG16L1 leads to a decrease in vascular reactivity and cardiac ejection fraction

Flow-mediated dilatation (FMD) in response to increased shear stress during occlusion-induced hyperhemia of the brachial artery is the standard and most used method in humans to non-invasively assess vascular endothelial dysfunction, an early predictor for CV events^25^. To explore whether deficiency in Rubicon or the WD domain of ATG16L1 in mice could affect FMD, we first adapted and optimized the measurement of endothelial-dependent vasodilatation in mice, using ultrasound imaging of the femoral artery^22^. An occlusion-cuff (O-cuff) of a plethysmograph CODA-monitor was used to reliably stabilize and standardize the pressure inducing the femoral artery ischemia over 5 min (Figure S1A-S1B). This adapted method of quantifying vascular endothelial function was validated to discriminate age-dependent endothelial dysfunction in C57Bl6 mice. We observed a significant decrease in FMD in 10-week-old compared to 20-week-old mice (Figure S1C). These data are in line with previous observations of age-related reduction of FMD in mice and humans in healthy conditions or affected by CV risk factors^22,26–28^. No changes were observed at the baseline measurement of femoral artery diameter or the pulse wave velocity between age-groups or the genotypes (Figure S1D and S1E). Importantly, deficiency in Rubicon or the WD domain of ATG16L1 in 2-month-old mice led to a significant decrease in FMD, suggesting that Rubicon and the WD of ATG16L induce an impairment of endothelial vascular reactivity in young mice, an early hallmark of CVD (Figure 1A and 1B).

**Figure 1.**
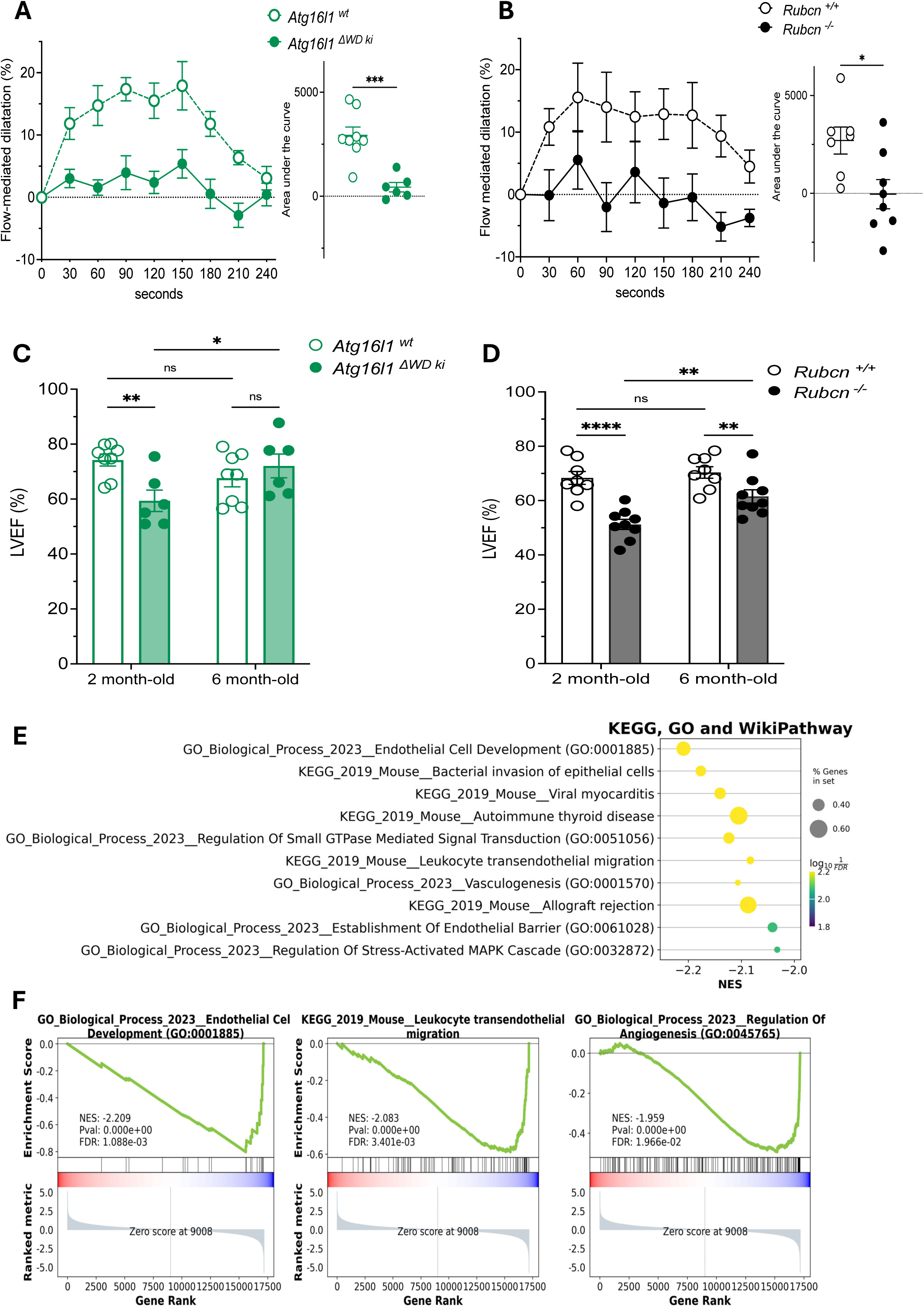
Full body deficiency in Rubicon and the WD40 domain of ATG16L1 in mice led to a decrease in vascular reactivity and a mild LV cardiac systolic dysfunction associated with a down-regulation of endothelial cell development in cardiac LV at a young age. **A** and **B** Flow-mediated dilation (FMD) measured in mice femoral artery during reactive hyperemia over 4 min in response to 5 min ischemia in (**A)** 2-month-old *Atg16l1^wt^* mice n=8 and *Atg16l1^DWD^ ^ki^* mice n=6; Values are expressed as mean±SEM. Significance was calculated for the FMD with a 2-way ANOVA followed by Tukey’s multiple comparison test (Time effect: *\*\*\*\*P<0.0001*; Genotype effect: *\*\*\*P<0.001;* Time x Genotype effect: *\*\*P<0.01*). Significance was calculated for the area under the curve with an unpaired Student *t* test *\*\*\*P<0.001*; (**B**) 2-month-old *Rubcn^+/+^* mice n=7 and *Rubcn^-/-^* mice n=8. Values are expressed as mean±SEM. Significance was calculated for the FMD with a 2-way ANOVA followed by Tukey’s multiple comparison test (Time effect: *\*P<0.05*; Genotype effect:*\*P<0.05*). Significance was calculated for the area under the curve with an unpaired Student *t* test *\*P<0.05*. **C** and **D** Percentage of Left Ventricle Ejection fraction (LVEF) in (**C**) 2-month-old and 6-month-old *Atg16l1^wt^* mice n=8 and *Atg16l1^DWD^ ^ki^* mice n=6; (**D**) 2-month-old and 6-month-old *Rubcn^+/+^*mice n=8 and *Rubcn^-/-^* mice n=9. Values are expressed as mean±SEM. Significance was calculated with a 2-way ANOVA where a Tukey post hoc test was applied. *\*\*P<0.01*, *\*P<0.05*. Color scheme represents deficiency in WD40 domain of ATG16L1 in green and deficiency in Rubicon in black. **E** and **F** Differentially expressed genes in cardiac left ventricle of 2-month-old *Rubcn^-/-^*mice n=6 versus *Rubcn^+/+^* mice n=6. A Negative normalized enrichment score (NES) value indicates the downregulation in the cardiac left ventricle of *Rubcn^-/-^* compared to *Rubcn^+/+^* mice (**E)** Bubble plots exhibiting pathway enrichment analysis of the genes differentially expressed in the LV between *Rubcn^-/-^*vs *Rubcn^+/+^* mice, showing the top 10 Kyoto Encyclopedia of Genes and Genomes (KEGG), Gene Ontology (GO) and WikiPathway. (**F**) Gene set enrichment analysis charts showing the enrichment of genes related endothelial development.

Non-invasive echocardiography recordings showed a significant decrease in left ventricle ejection fraction (LVEF) in 2-month-old *Atg16l1^DWD^ ^ki^* and *Rubcn^-/-^* mice versus their littermate controls (Figure 1C and 1D). At 6-months, this LVEF reduction was significant for the *Rubcn^-/-^*mice versus their littermate controls but not for the *Atg16l1^DWD^ ^ki^* mice (Figure 1C and 1D). Since the LVEF values observed for both genotypes were all above a threshold of 50% (considered as normal or subnormal), this significant decrease in LVEF could be defined according to the America Heart Association guidelines in humans as cardiac dysfunction or heart failure (HF) with preserved or mild reduced LVEF^29^. HF with preserved LVEF is a complex syndrome associated in humans with a high mortality rate comparable to heart failure with reduced LVEF (<50%). Interestingly, in both *Atg16l1^DWD^ ^ki^* and *Rubcn^-/-^* mice versus their littermate controls, significant increases in LV end-systolic volume or diameter were observed (Table S1 and table S2), which are considered as reliable predictors of early onset with HF preserved LVEF. Of note, a LVEF reduction and increase LV end-systolic volume were observed only in male but not in female *Rubcn^-/-^* mice (Table S2 and table S3). No changes in electrocardiography waveforms (ECG), systolic, diastolic or mean blood pressure values were observed in *Atg16l1^DWD^ ^ki^*or *Rubcn^-/-^*mice versus their littermate controls, indicating that these genotypes were not associated with arrythmia or hypertension (Figure S1B, S1C and S1D) during the period measured. While a weight gain was observed over time between 2-month-old and 6-month-old mice, no significant changes were observed in body weights between the genotypes at either period measured.

Collectively, our data showed that deficiency in Rubicon and the WD domain of ATG16L1 induced a significant decrease in endothelial vascular reactivity and a mild reduction in cardiac functions, which was not associated with changes with traditional CV comorbidities such as hypertension, weight gain, hypercholesterolemia or hyperglycemia (Table S4).

No signs of fibrosis or infiltration of immune cells were observed in 2-month-old mice; therefore, to gain insight into the underlying pathways involved in the cardiac dysfunctions observed in young *Rubcn^-/-^*mice, we performed a bulk mRNA seq analysis in their apex and left ventricles of cardiac tissues. Gene set enrichment analysis revealed that GO term pathways such as endothelial cell development, vasculogenesis, establishment of endothelial barrier, and regulation of angiogenesis, and KEGG_2019 term leukocyte trans-endothelial migration, were among the 10 top pathways significantly downregulated in the cardiac tissue of *Rubcn^-/-^*mice compared to their littermate controls (Figure 1E). These observations suggest that vascular endothelial function seems to be a key pathway that was dampened in the hearts of our mice and could be responsible for LVEF reduction.

### Cardiac and endothelial dysfunction observed in full body Rubicon deficiency are both endothelial dependent in 2-month-old mice

To support our transcriptomic findings highlighting the importance of endothelial cell compartments in cardiac dysfunction induced by non-canonical autophagy deficiency, we generated an endothelial-specific *Rubcn*-depletion mouse model, *Rubcn^fl/fl^, Cdh5-Cre^Tg+^.* An endothelial-specific Cre recombinase (*Cdh5-Cre*) enabled cell-type-specific depletion of a functional *Rubcn* gene, confirmed by genotyping and western-blot in pulmonary endothelial cells (Figure S3). As observed in the full-body deficiency, non-invasive echocardiography recordings showed a significant decrease in left ventricle ejection fraction (LVEF) in 2-month-old in *Rubcn^fl/fl^, Cdh5-Cre^Tg+^* mice versus their littermate controls *Rubcn^fl/fl^, Cdh5-Cre^Tg-^* (Figure 2A). No changes in LV end-diastolic volume or diameter were observed in those mice (Table S6).

**Figure 2.**
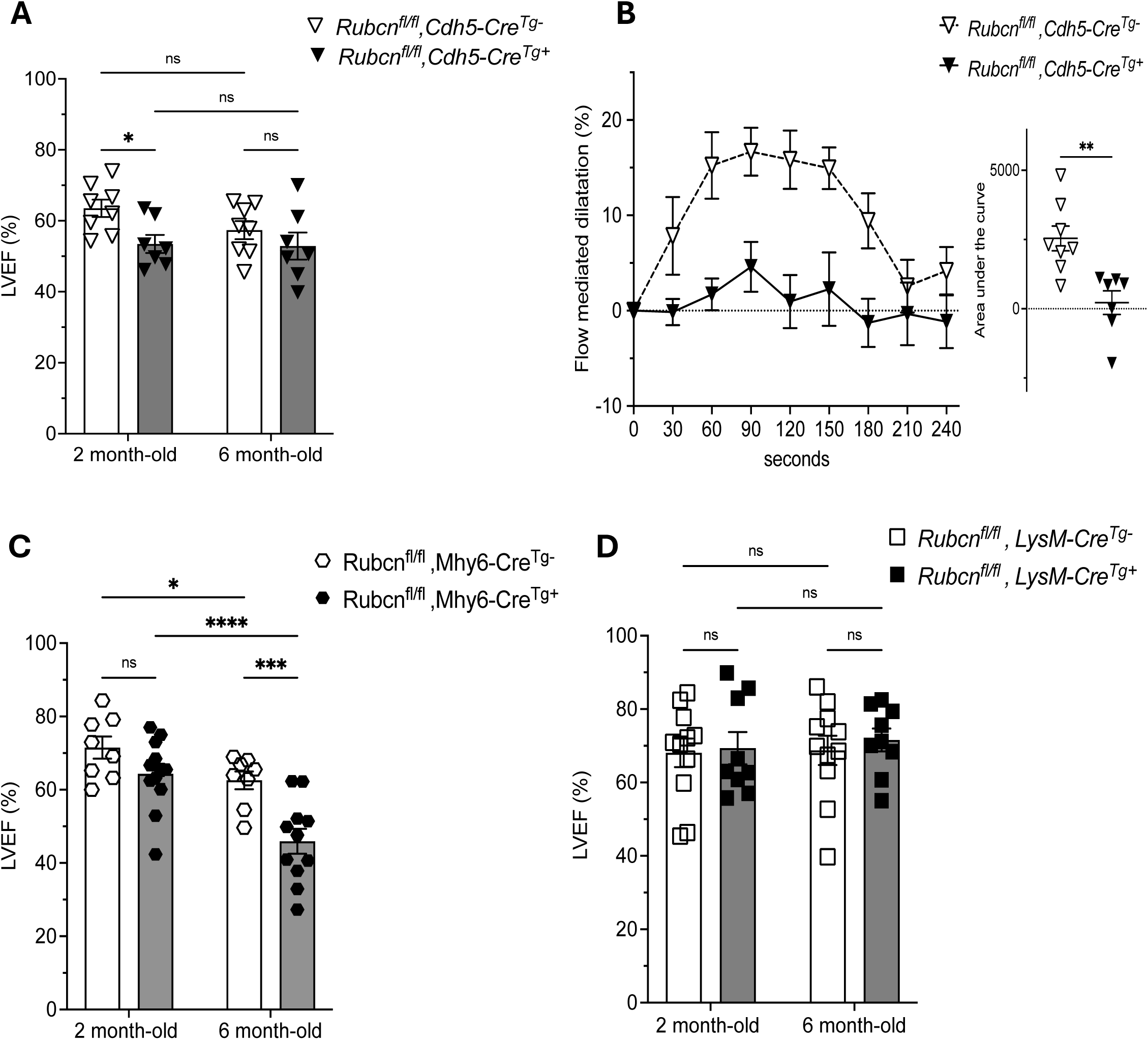
The mild LV cardiac systolic dysfunction and the decrease in vascular reactivity observed in full body deficiency in Rubicon deficient mice are both endothelial-dependent at a young age. **A.** Percentage of Left Ventricle Ejection fraction (LVEF) in 2-month-old and 6-month-old *Rubcn^fl/fl^, Cdh5-Cre^Tg+^* n=7 and *Rubcn^fl/fl^, Cdh5-Cre^Tg-^* mice n=8; Values are expressed as mean±SEM. Significance was calculated with a 2-way ANOVA where a Tukey post hoc test was applied. *\*P<0.05*. **B.** Flow-mediated dilation (FMD) measured in mice femoral artery during reactive hyperemia over 4 min in response to 5 min ischemia in 2-month-old mice *Rubcn^fl/fl^,Cdh5-Cre^Tg+^* n=7 and *Rubcn ^fl/fl^,Cdh5-Cre^Tg-^*mice n=8; Values are expressed as mean±SEM. Significance was calculated for the FMD with a 2-way ANOVA followed by Tukey’s multiple comparison test (Time effect: *\*\*\*P<0.001*; Genotype effect: *\*\*P<0.01;* Time x Genotype effect: *\*P<0.05*). Significance was calculated for the area under the curve with an unpaired Student *t* test *\*\*P<0.01*; **C** and **D** Percentage of LVEF in (**C**) 2-month-old and 6-month-old *Rubcn^fl/fl^, Myh6-Cre^Tg+^* n=8 and *Rubcn^fl/fl^, Myh6-Cre^Tg-^* mice n=11; (**D**) 2-month-old and 6-month-old *Rubcn^fl/fl^, LysM-Cre^Tg+^*n=9 and *Rubcn^fl/fl^, LysM-Cre^Tg-^* mice n=11 . Values are expressed as mean±SEM. Significance was calculated with a 2-way ANOVA where a Tukey post hoc test was applied. *\*\*\*\*P<0.0001*, *\*\*\*P<0.001*, *\*P<0.05*.

As expected, and supporting our earlier observations in full body *Rubcn* deficient mice, *Rubcn^fl/fl^, Cdh5-Cre^Tg+^* also showed a significant decrease in FMD (Figure 2B), demonstrating that the impairment of vascular reactivity induced by non-canonical autophagy deficiency was effectively endothelial-dependent. To further delineate whether myeloid cells or cardiomyocytes could participate in the cardiac dysfunctions at young age induced by non-canonical autophagy, we generated a myeloid-specific *Rubcn*-depletion mouse model, *Rubcn^fl/fl^, LysM-Cre^Tg+^* using a myeloid-specific Cre recombinase (*LysM-Cre*) and a cardiomyocyte-specific *Rubcn*-depletion mouse model, *Rubcn^fl/fl^,Myh6-Cre^Tg+^*using a cardiomyocyte-specific Cre recombinase (*Myh6-Cre*). While a significant decrease in LVEF was observed in 6-month-old *Rubcn^fl/fl^, Myh6-Cre^Tg+^*compared to their littermate controls, no changes in cardiac functions were observed in 2-month-old *Rubcn^fl/fl^,LysM-Cre^Tg+^* or *Rubcn^fl/fl^, Myh6-Cre^Tg+^* compared to their littermate controls (Figure 2C and 2D).

Taken together, these in vivo measurements of cardiac and endothelial functions in cell-specific depleted mice supports the transcriptomic analysis in the full body *Rubcn* deficient mice, strongly suggesting that the endothelial cell compartment plays a predominant role in the early onset of cardiac dysfunction induced by deficiencies in non-canonical autophagy.

### Rubicon deficiency leads to an increase in cardiac vasculature leakiness

To further assess whether non-canonical autophagy deficiency could perturb endothelial barrier functions throughout the whole heart, we adapted a 3D visualization of coronary vasculature approach by quantitative light sheet fluorescence microscopy^30^. For this purpose, we used a simultaneous intravenous injection of fluorescent anti-CD31 antibodies with fluorescently labelled 70kDa-Dextran (Figure S4A). As a marker of cardiac vascular permeability, mutual information analysis quantified Pearson correlation coefficients between CD31-Alexa647 and Dextran-TRITC, over 1000 bootstrap selections for 30 distinct morphological cardiac regions located either in the left ventricle, interventricular septum, or the right ventricle (Figure S4B and S4C). While averaging the Pearson correlation coefficients quantified over all cardiac regions did not show any changes between the genotypes (Figure 3B and 3E), a significant decrease in cardiac vascular permeability was observed in the right ventricle regions in both 2-month-old *Rubcn^-/-^* and *Atg16l1^DWD^ ^ki^* mice versus their littermate controls (Figure 3C and 3F). Our data suggest that non-canonical autophagy deficiency led to a CV phenotype which shared two pathophysiologic features for the very complex and heterogenous heart failures with mildly reduced ejection fraction in humans: right ventricular and coronary microvascular endothelial dysfunctions^31–33^.

**Figure 3.**
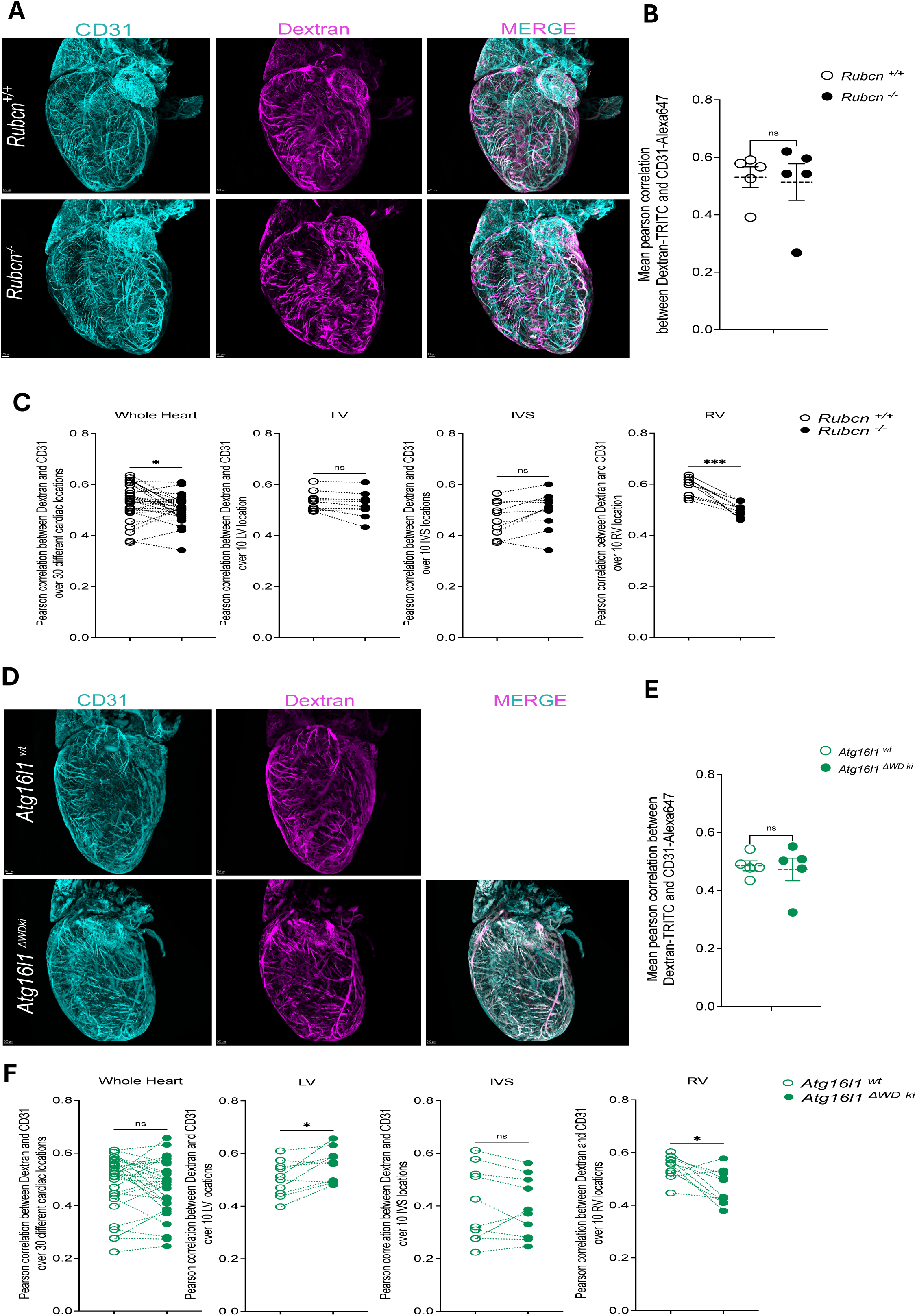
Full body deficiency in Rubicon and the WD40 domain of ATG16L1 in mice led to an increase in right ventricle vascular leakiness. **A** and **D.** 3D renderings of representative cardiac volume vasculature injected with Dextran70kDa-TRITC (Magenta) after ranking the orientation responses of pathway operators (RORPO) processing for vessel enhancement of CD31-Alexa647 (Turquoise) fluorescent signals in (**A**) 2-month-old *Rubcn^+/+^* mouse and *Rubcn^-/-^*mouse; and (**D**) 2-month-old *Atg16l1^wt^* mice n=4 and *Atg16l1^DWD^ ^ki^* mice n=4; Scale bar are values in mm. **B** and **E.** Mean of pearson correlation between CD31-Alexa647 and Dextran-TRITC quantified by mutual information analysis over 1000 boostrap selection for each tissue sub-volume cropped specifically at 30 different regions of interests for cardiac volumes of (**B**) 2-month-old *Rubcn^+/+^* mice n=4 and *Rubcn^-/-^* mice n=4; (**E**) 2-month-old *Atg16l1^wt^* mice n=4 and *Atg16l1^DWD^ ^ki^* mice n=4 . Values are expressed as mean±SEM. Significance was calculated for the pearson correlation with an unpaired Student *t* test. **C** and **F.** Pearson correlation coefficients between CD31-Alexa647 and Dextran-TRITC quantified by mutual information analysis over 1000 boostrap selection for each tissue sub-volume cropped either at 30 different regions of interests over the whole heart volumes, or at 10 different regions of interests over the left ventricle (LV), or the inter ventricular septum (IVS), or the right ventricle (RV) (**C**) in 2-month-old *Rubcn^+/+^* mice n=4 and *Rubcn^-/-^* mice n=4; (F) in 2-month-old *Atg16l1^wt^* mice n=4 and *Atg16l1^DWD^ ^ki^* mice n=4. Values are expressed as mean±SEM. Significance was calculated for the pearson correlation with an unpaired Student *t* test. *\*P<0.05*, *\*\*P<0.01*, *\*\*\*P<0.001*.

### Deficiency in Rubicon and the WD domain of ATG16L1 disrupts VEGFR2 recycling and nitric oxide pathway induced by shear stress

We then sought to explore the signaling pathway involved in the endothelial dysfunction induced by non-canonical autophagy deficiency *in vitro*. Endothelial cells are constantly exposed to fluid shear stress from blood flow which plays a critical role in vascular homeostasis and in determining endothelial functions. At physiological levels, endothelial cell responses to laminar shear stress were found to be mediated by a complex of proteins comprising VEGFR2, PECAM1 and VE-Cadherin^34^. We observed that laminar shear stress induced the recycling of the mechanosensor VEGFR2 on the plasma membranes of primary human endothelial aortic cells (HAEC) (Figure 4A).

**Figure 4.**
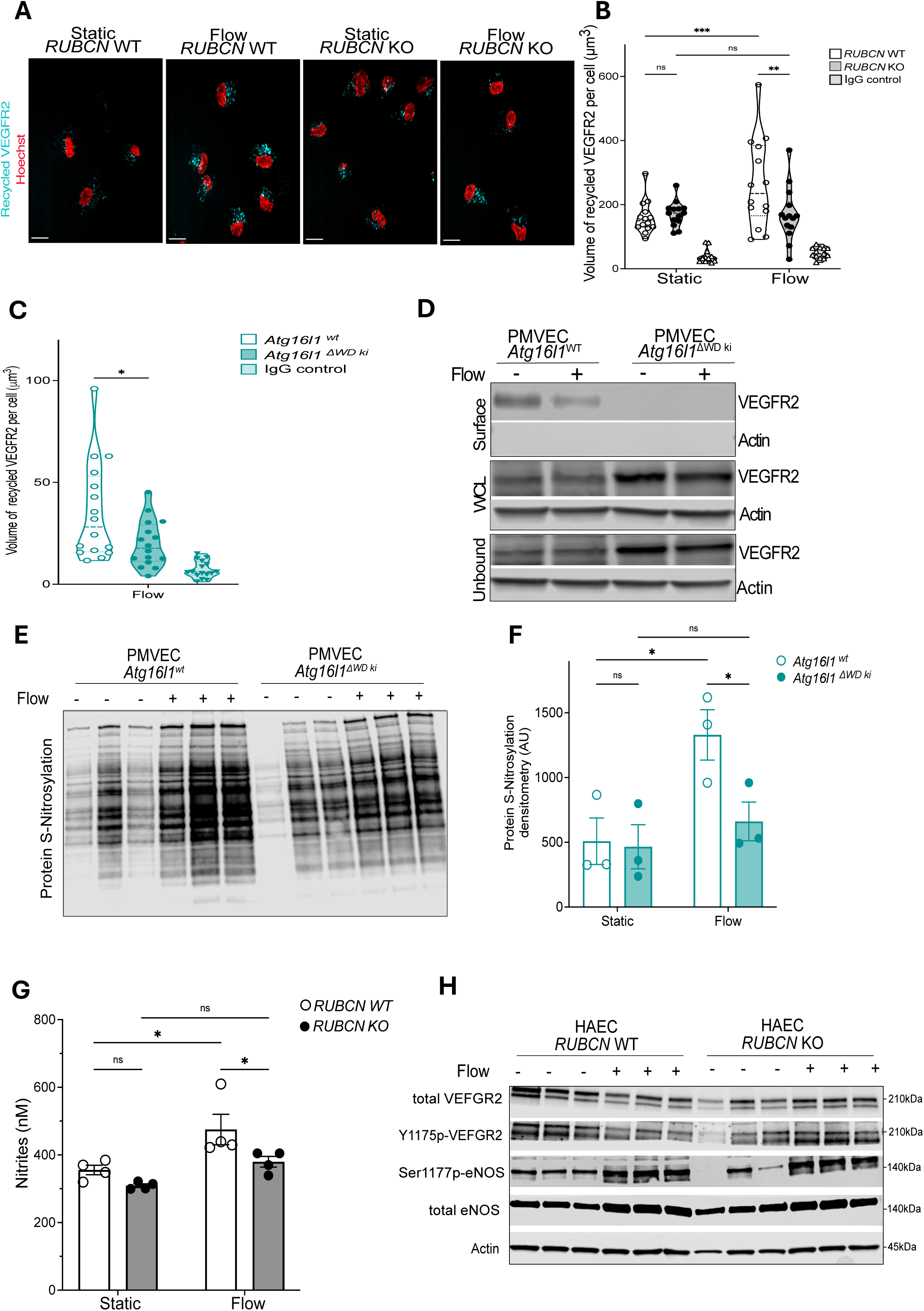
Deficiency in Rubicon and the WD40 domain of ATG16L1 in murine and human EC led to a decrease in VEGFR2 recycling and NO pathways which is not associated with S1177 eNOS and Y1175 VEGFR2 phosphorylation. **A**. Recycled VEGFR2 stained with Alexa674 (Cyan) and nuclei stained with Hoechst (Red) in human aortic endothelial cells (HAEC) *RUBCN* KO and *RUBCN* WT exposed to static or 5 dynes of laminar flow during 24h. Scale bars, 1mm. Images are representative of three independently performed experiments. **B.** Volume of recycled VEGFR2 per cell with each point representing the average volume for at least 3 cells per field view from HAEC *RUBCN* KO and *RUBCN* WT (n=3 independent replicates per group) exposed to static or 5 dynes of laminar flow during 24h. IgG controls as a control for VEGFR2 antibody. Values are expressed as mean±SEM. Significance was calculated for the Volume of recycled VEGFR2 with a 2-way ANOVA followed by tukey’s multiple comparison test (Flow exposure effect: *\*P<0.05*; Genotype effect: *\*\*\*\*P<0.0001;* Flow exposure Genotype effect: *\*P<0.05*). **C.** Volume of recycled VEGFR2 per cell with each point representing the average volume for at least 3 cells per field view from murine PMVEC isolated from 2-month-old *Atg16l1^wt^*mice n=3 and *Atg16l1^DWD^ ^ki^* mice n=3 then exposed to static or 5 dynes of laminar flow during 24h. Values are expressed as mean±SEM. Significance was calculated for the volume of VEGFR2 recycled per cell with an unpaired Student *t* test*. *P<0.05* **D.** Representative immunoblots of VEGFR2 and actin in isolated surface, whole-cell lysate (WCL) and biotin unbound fractions across murine PMVEC isolated from 2-month-old *Atg16l1^wt^* and *Atg16l1^DWD^ ^ki^* mice then exposed to static or 5 dynes of laminar flow during 24h. **E.** Immunoblots of total protein S-nitrosylation in whole cell lysate from PMVEC isolated from 2-month-old *Atg16l1^wt^* mice n=3 and *Atg16l1^DWD^ ^ki^* mice n=3, then exposed to static or 5 dynes of laminar flow during 24h. **F.** Total protein S-nitrosylation densitometry measured by western blot in PMVEC isolated from 2-month-old *Atg16l1^wt^* mice n=3 and *Atg16l1^DWD^ ^ki^* mice n=3 then exposed to static or 5 dynes of laminar flow during 24h. Values are expressed as mean±SEM. Significance was calculated for the densitometry with a 2-way ANOVA followed by tukey’s multiple comparison test. **G.** Nitrites measured in cell culture medium in HAEC *RUBCN* KO and *RUBCN* WT (n=3 independent replicates per group) exposed to static or 5 dynes of laminar flow during 24h.Values are expressed as mean±SEM. Significance was calculated for the nitrites with a 2-way ANOVA followed by tukey’s multiple comparison test (Flow exposure effect: *\*\*P<0.01*; Genotype effect: *\*P<0.05*). **H.** Immunoblots of total VEGFR2, Y1175-phosporylated-VEGFR2, Ser1177-phosphorylated-eNOS, total eNOS and b-actin proteins in whole cells lysates of HAEC *RUBCN* KO and *RUBCN* WT (n=3 independent replicates per group) exposed to static or 5 dynes of laminar flow during 24h.

We ablated Rubicon in HAEC using CRSPR-Cas9 and isolated microvascular endothelial cells from the lungs (PMVEC) of *Rubcn^-/-^* and *Atg16l1^DWD^ ^ki^* mice. We observed that laminar shear stress-induced VEGFR2 recycling was significantly dampened in endothelial cells deficient in Rubicon or the WD domain of ATG16L1 compared to controls (Figure 4A-C). Furthermore, the recycling on the plasma membrane of other mechanosensors such as PECAM-1, VE-cadherin or Piezo-1 remained unchanged between the genotypes in static or shear stress conditions (Figure S5A). In murine PMVEC we observed a reduction of VEFGR2 detected at the plasma membrane by surface biotin labeling in both static and shear stress conditions between cells deficient in Rubicon or the WD domain of ATG16L1 compared to controls (Figure 4D).

While adequate levels of bioactive nitric oxide (NO) produced by endothelial nitric oxide synthase (eNOS) are critical for the regulation of blood vessel vasodilatation, accumulating evidence has demonstrated that NO is often stabilized in the intravascular compartment by the formation of nitrites, S-nitrosothiols and S-nitrosylated proteins^35^. We assessed nitrites in the HAEC cell culture medium and total protein S-nitrosylation in cell lysates and found that deficiency in Rubicon or the WD domain of ATG16L1 in endothelial cells disrupted laminar shear stress-induced NO production (Figure 4F and 4G). Interestingly, non-canonical autophagy deficiency did not lead to any changes in the phosphorylation levels of the sites in eNOS (endothelial nitric oxide synthase) that are known to impact its activity (Ser 1179 and Thr495). No changes were observed in the levels of phosphorylation of VEGFR2 between the genotypes (Figure 4H). Taken together, these results revealed that a decrease in NO pathways *in vitro* are in line with endothelial dysfunctions induced by non-canonical autophagy deficiency *in vivo*. However no noticeable inflections in eNOS activity, the constitutive enzyme responsible for NO production in endothelial cells, were observed at the timeframe studied. This led us to investigate the upstream signaling components that could transduce a lack in recycling mechanosensors at the surface of plasma membrane into a decrease in NO pathways in endothelial cells.

### LANDO deficiency leads to a decrease in protein expression of hemoglobin subunit alpha (Hb-⍺) in endothelial cells

To identify and quantify the cell surface proteome that could be differentially affected by non-canonical autophagy depletion in endothelial cells, we performed surface biotin labeling coupled with unbiased TMT-proteomic analysis on Rubicon depleted HAEC (Figure S5B). This methodology had been previously used to describe numerous cell surface cargoes dependent on endosomal retrieval and recycling pathways^36^. While contaminations with intracellular fractions could not be excluded with this approach, we validated that the isolated biotinylated proteins through streptavidin affinity capture were enriched for protein expression related to components of plasma membrane (Figure S5C-E). Of a total of 903 quantified proteins, 3 proteins (HBA1, CXCR4 and TSPAN4) were differentially expressed from the surface of Rubicon-depleted HAEC under laminar shear stress compared to the control cells in four independent experiments (Figure 5A, 5B). Consistent with earlier observations in human umbilical vascular endothelial cells ^37^, the decrease in the chemokine receptor CXCR4 protein expression induced by laminar shear in HAECs was validated by western blot analysis in isolated surfaces, whole-cell lysates and biotin unbound fractions (Figure 5C).

**Figure 5.**
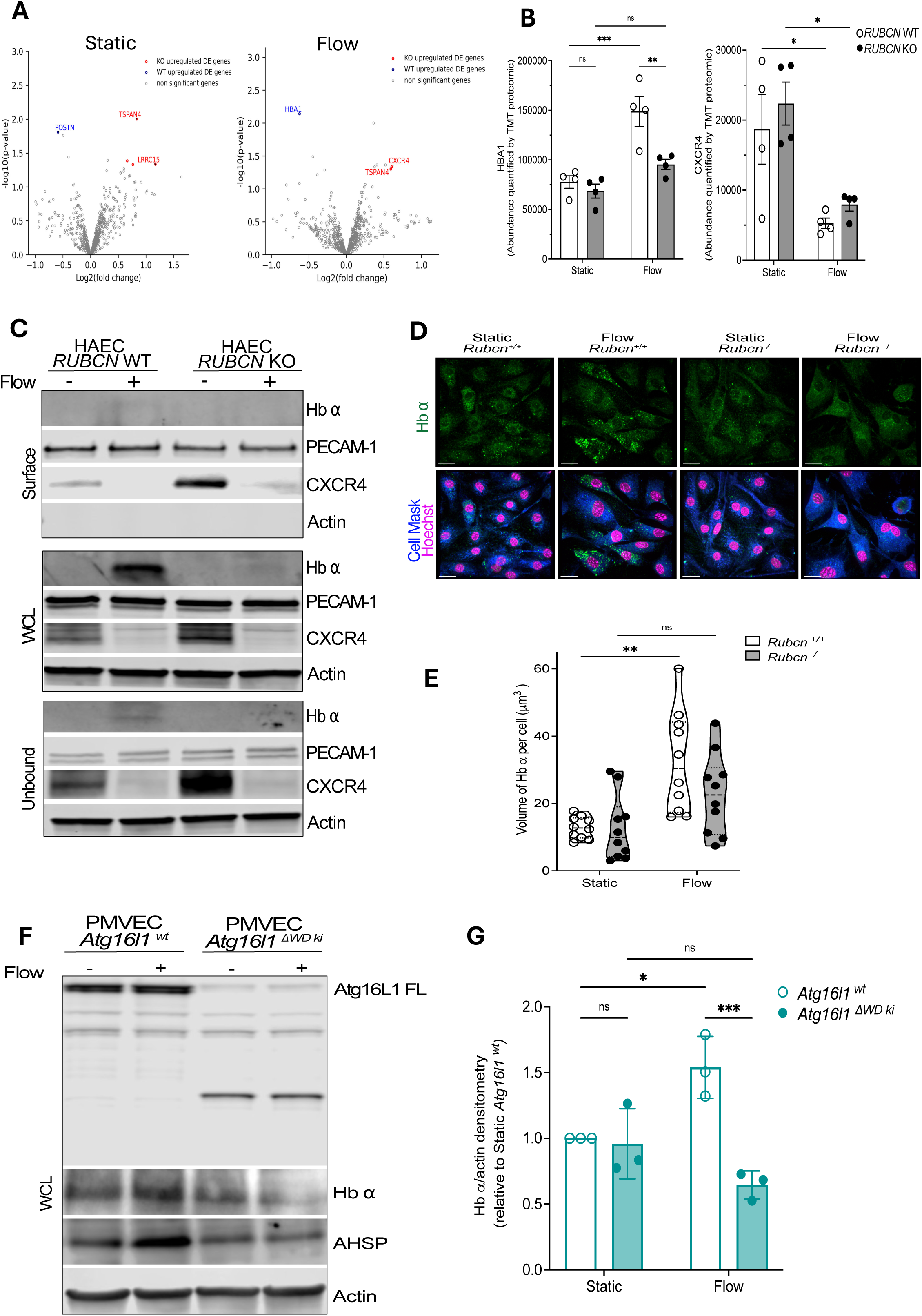
Deficiency in Rubicon and the WD40 domain of ATG16L1 in HAEC and murine pulmonary EC led to a decrease in hemoglobin subunit alpha protein expression. **A.** Volcano plots of differentially expressed surface proteins across HAEC *RUBCN* KO and *RUBCN* WT (n=4 independent replicates per group) in response to static or 5 dynes of laminar flow during 24h, with proteins considered as differentially expressed (fold change ³1.5 and *P<0.05*) colored in red (upregulated in HAEC *RUBCN* KO compared to *RUBCN* WT) or in blue (downregulated in HAEC *RUBCN* KO compared to *RUBCN* WT). **B.** Protein abundance of HBA1 and CXCR4 assessed by TMT proteomics in HAEC *RUBCN* KO and *RUBCN* WT (n=4 independent replicates per group) exposed to static or 5 dynes of laminar flow during 24h.Values are expressed as mean±SEM. Significance was calculated for protein abundance with a 2-way ANOVA followed by tukey’s multiple comparison test. **C.** Representative immunoblots of Hemoglobin subunit a (Hb a), PECAM-1, CXCR4 and actin in isolated surface, whole-cell lysate (WCL) and biotin unbound fractions across HAEC *Rubcn* KO and *Rubcn* WT in response to static or 5 dynes of laminar flow during 24h. **D.** Representative fluorescent images of intracellular Hb a stained in murine PMVEC isolated from 2-month-old *Rubcn^+/+^* mice and *Rubcn^-/-^* mice and exposed to static or 5 dynes of laminar flow during 24h. **E.** Volume of intracellular Hb a per cell with each point representing the average volume for at least 8 cells per field view from 2-month-old *Rubcn^+/+^*(n=3) and *Rubcn^-/-^*(n=3) mice exposed to static or 5 dynes of laminar flow during 24h. Values are expressed as mean±SEM. Significance was calculated for the Hb a volume of with a 2-way ANOVA followed by tukey’s multiple comparison test. **F.** Representative immunoblots of Atg16L1, Hb a, Alpha-Hemoglobin stabilizing protein (AHSP) and actin in whole-cell lysate (WCL) fraction across pulmonary PMVEC isolated from 2-month-old *Atg16l1^wt^*mice and *Atg16l1^DWD^ ^ki^* mice, then exposed to static or 5 dynes of laminar flow during 24h. **G.** Relative Hb a to actin densitometry measured by western blot in PMVEC isolated from 2-month-old *Atg16l1^wt^* mice n=3 and *Atg16l1^DWD^ ^ki^* mice n=3 then exposed to static or 5 dynes of laminar flow during 24h. Values are expressed as mean±SEM. Significance was calculated for the densitometry with a 2-way ANOVA followed by tukey’s multiple comparison test.

Strikingly, we observed an increase in hemoglobin subunit alpha (HBA1 or Hb-⍺) protein expression induced by laminar shear stress in control HAECs, which was completely dampened in *RUBCN*-deficient HAECs (Figure 5C). These immunoblot results validated the proteomic analysis in the whole-cell lysates of HAEC. Hb-⍺ is best known as an oxygen transporter in red blood cells. While previous reports have demonstrated that Hb-⍺ was also expressed in mouse and human arterial endothelial cells, where it controls NO production and vasodilatation^38,39^, little is known about the regulation of Hb-⍺ protein expression by laminar shear stress. IF staining of endothelial cells confirmed an increase in intracellular Hb-⍺ protein expression induced by laminar shear stress, which was blunted in murine PMVEC isolated from *Rubcn^-/-^*mice compared to the controls (Figure 5D-E). Similar results regarding Hb-⍺ protein expression and its chaperone ⍺-hemoglobin stabilizing protein (AHSP) were observed in whole cell lysates of murine PMVEC isolated from *Atg16l1^DWD^ ^ki^* mice versus the controls (Figure 5F-G). It is unlikely that Hb-⍺ is a cell surface protein, and therefore we suspect that our ability to detect it in our proteomic screen relates to an association with an unkown surface protein. Nevertheless, our data suggest that deficiency in Rubicon or the WD domain of ATG16L1 in endothelial cells can regulate shear stress induced Hb-⍺ protein upregulation.

### Identification of candidate genes related to the LANDO pathway associated with CV parameters in the Young Finns Study and the UK biobank

We next explored whether other candidate genes in LANDO and receptor recycling pathways beyond *RUBCN* and the WD domain ATG16L1, could be associated with CV parameters in humans and a modulation in Hb-⍺ protein expression. Genome wide association studies (GWAS) have shed light on genetic variants involved in pathways known to play a role in the physiopathology of CV parameters^40^. While increasingly larger sample sizes, multi-ancestry analyses, and inclusion of rare variants are improving GWAS power, reports suggesting the association of genetic variants present in loci harboring autophagy genes with CV parameters are rare^1^. To increase the power to detect associations of genetic variants in genes involved in LANDO with CV parameters, we used genetic expression studies to explore whether genetic variants that negatively or positive impact the expression of genes or proteins related to LANDO, were associated with CV parameters. A set of 42 genes were selected for their potential implication in the LANDO and receptor recycling pathways at different steps of mechanosensing, PI3P generation, LC3-conjugation, retromer and retriever complexes, and V-ATPase^41–43^ (Figure 6A). Selection of genetic variants in those 42 genes was based on 3 criteria: (1) their presence in the coding regions or the regulatory regions excluding the intronic, non-coding, intergenic and synonymous regions, and (2) significant changes observed in cis-expression quantitative trait loci (eQTL) from 2 public database in whole-blood (n=31684) and from the Genotype-Tissue Expression project (GTEX v8) across 10 neuro-cardiovascular tissues (n=838), or (3) significant changes in protein quantitative trait loci (pQTL) in the cortical brain regions (n=330), the only pQTL database in neuro-cardiovascular tissues where genetic variants in LANDO pathway gene loci were found significant until 2023 (Table S9).

**Figure 6.**
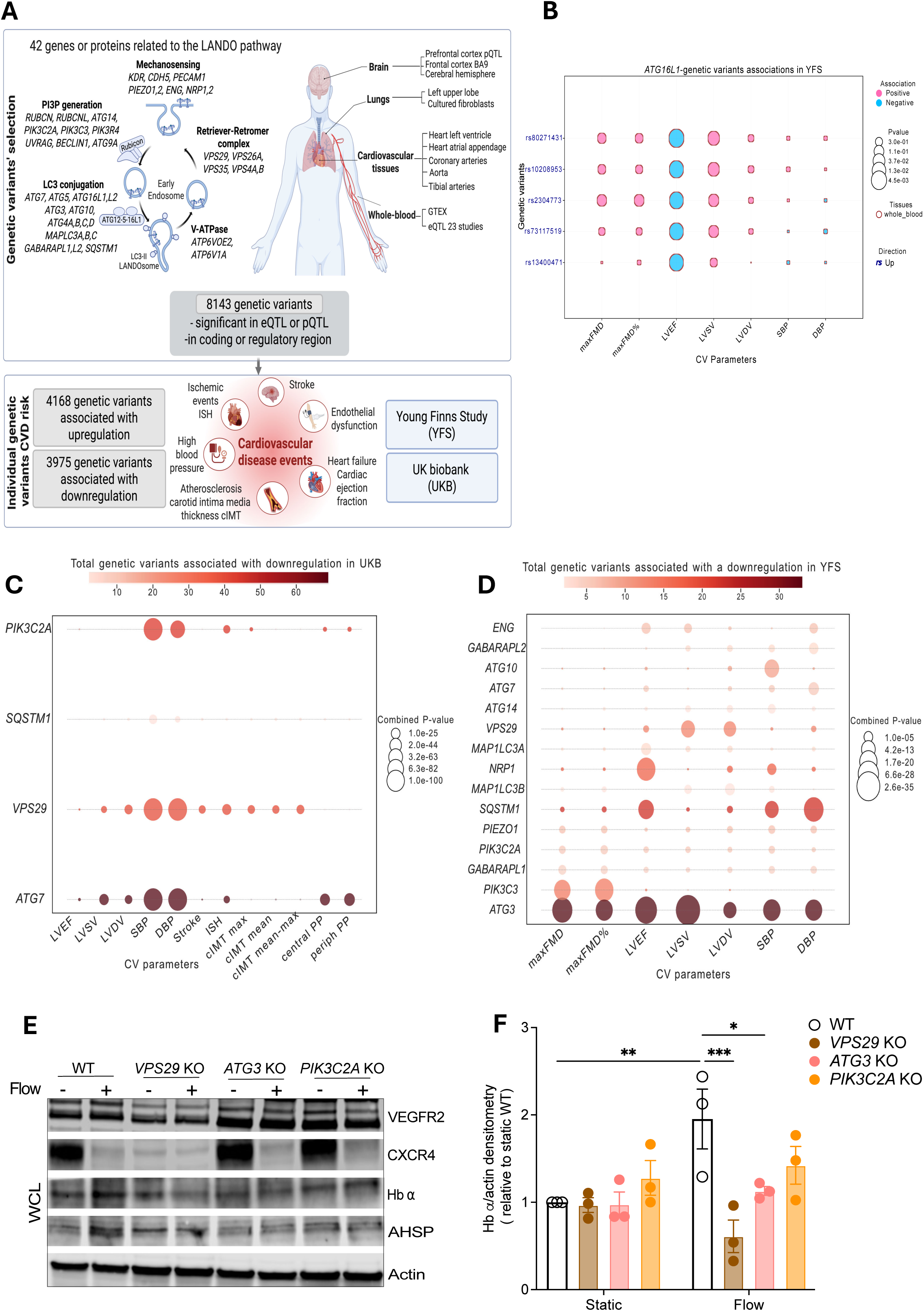
15 genetic loci and several candidate genes related to Rubicon and ATG16L1 pathway were associated with cardiovascular parameters in the Young Finns study and the UK biobank study. **A.** Workflow depicting a first step of genetic variants selection in 42 genes or proteins related to the LANDO pathway with a significant quantitative trait loci (QTL) in 11 CV related tissues from the genotype-tissue expression project; (GTEX n=838 individual tissue samples), the eQTLGen Consortium (n=31684 whole-blood samples) and pQTL (330 post-mortem dorsolateral prefrontal cortex), then a second step of individual genetic variants association analyses with cardiovascular (CV) parameters in the Young Finns Study and the UK Biobank study. **B.** Bubble plot exhibiting a single genetic variants analysis in *ATG16L1* loci associated with 7 CV parameters (Flow-mediated dilatation: FMD; Left ventricle ejection fraction: LVEFD; Left ventricle Systolic Volume; Left ventricle diastolic Volume: LVDV; Systolic blood pressure: SBP and Diastolic blood pressure: DBP) in the Young Finns Study. The dark blue color font indicates that the genetic variants in *ATG16L1* loci were significantly associated with an upregulation of ATG16L1 mRNA in the whole blood. The bubble size indicates the P-value for the association and the color bar indicates the direction of the association (positive in pink or negative in blue). **C** and **D.** Bubble plot exhibiting combined genetic variants analysis associated with CV parameters in (**C**) the UK biobank study or in (**D**) Young Finns study. The bubble size indicates the combined P-value for the association and the color bar indicated the number of total genetic variants associated with a downregulation for each gene or protein related to ATG16L1 and Rubicon pathway. **E.** Representative immunoblots of VEGFR2, CXCR4, Hb a, Alpha-Hemoglobin stabilizing protein (AHSP) and actin in whole-cell lysate (WCL) fraction across HAEC *VPS29* KO, *ATG3* KO and *PIK3C2A* and WT in response to static or 5 dynes of laminar flow during 24h. Relative Hb a to actin densitometry measured by western blot in HAEC *VPS29* KO, *ATG3* KO and *PIK3C2A* and WT exposed to static or 5 dynes of laminar flow during 24h (n=3 independent replicates per group). Values are expressed as mean±SEM. Significance was calculated for the densitometry with a 2-way ANOVA followed by tukey’s multiple comparison test.

Single variant analyses were first tested in 402137 individuals of the UK Biobank (UKB) for association with cardiac function parameters (LVEF, LVSV, LVDV), blood pressure parameters (SBP, DBP), CV events (stroke, ischemic heart disease, venous thromboembolism), carotid Intima-media thickness (cIMT) parameters that assessed atherosclerosis and 2 proxy parameters of endothelial function, the central and peripheric pulse pressures. Single variant analyses were next tested in about 2000 individuals of the Young Finns Study (YFS) for association with the same cardiac and blood pressure parameters described for the UKB, expanded with direct endothelial function parameters (Max FMD raw and Max FMD in percentage). As a first line of analyses, we examined whether genetic variants in *RUBCN* or *ATG16L1* loci that were in eQTLs in neuro-cardiovascular tissues, were also associated with CV parameters. Only in the YFS, we observed that 5 genetic variants (P<1×10⁻²) in *ATG16L1* locus, which minor alleles were associated with upregulated mRNA expression levels in whole-blood, showed associations consistent with CV regulation: increased endothelial function, and lower cardiac functions (lower LVEF and higher LVDV or LVSV) as described in Figure 6B. These results suggested that increased expression levels of *ATG16L1*, a gene shared in canonical and non-canonical autophagy, is associated with a beneficial effect on endothelial functions, which is consistent with our mice data on the WD domain of ATG16L1. However, these genetic variants in *ATG16L1* were not associated with beneficial effects on cardiac functions. No genetic variant in the *RUBCN* locus were found associated with CV parameters in the YFS or the UKB

Next, we sought to identify candidate genes related to the LANDO pathway that showed the strongest pattern of genetic association with CV parameters. We aggregated genetic variants signals to genes across twelve CV parameters in the UKB and seven CV parameters in the YFS (Figure 6C-6D). This approach allowed us to identify 12 novel loci in which genetic variants present in genes related to non-canonical autophagy were associated with CV parameters. For each gene, the list of genetic variants and their association with CV parameters are described Figure S7-S14. Three genes involved in mechanosensing (*ENG*, *NRP1*, PIEZO1) and 2 autophagy genes (*ATG7*, *SQSTM1*) served as positive controls since their lower expression had previously been associated with endothelial and/or cardiac dysfunctions^44–48^. Two genes *PIK3C2A* and *VPS29* overlapped between the combined analysis in the YFS and the UKB, and 1 gene, *ATG3* had the strongest associations for all CV parameters tested in the YFS.

To validate these results *in vitro*, *ATG3*, *PIK3C2A*, and *VPS29* were silenced in HAEC and Hb-⍺ protein expression levels were assessed under static or laminar shear stress. We found that, similar to *RUBCN* and the WD domain of ATG16L1, *VPS29* and *ATG3*-deficiency in endothelial cells prevented shear stress-induced Hb-⍺ protein upregulation (Figure 6E-F).

## Discussion

Our data using mice deficient in Rubicon or the WD domain of ATG16L1, murine and human endothelial cells, and a candidate pathway approach in humans consistently revealed that non-canonical functions of autophagy proteins play a pivotal role in the pathophysiology of endothelial and cardiac dysfunction.

Decline in NO bioavailability or production is widely recognized as a key mechanism involved in endothelial and cardiac dysfunctions^49^. Surprisingly, using an unbiased proteomic approach, we discovered that a decrease in intracellular Hb-⍺ protein expression was associated with lower NO bioavailability in non-canonical autophagy-deficient endothelial cells. The laboratories of Isakson and Weiss described a non-erythroid expression of Hb-⍺ mainly in endothelial cells at the myoendothelial junctions from small resistance arteries^38,50^. It has also been demonstrated that deficiency in Hb-⍺ in endothelial cells displayed an improved endothelial function. Through its heme-associated iron, Hb-⍺ could directly bind eNOS and degrade NO in endothelial cells^38,39^. Our data provide a physiological context in which Hb-⍺ is regulated in endothelial cells. We found that laminar shear stress robustly upregulated intracellular Hb-⍺ protein expression in endothelial cells in microvascular pulmonary vessels but also in large arteries such as aorta where Hb-⍺ had not been previously detected.

Recent gain-of-function and chromatin immunoprecipitation studies in HAECs have revealed the presence in the *HBA1* promotor region of transactivation binding sites for the flow-induced transcription factor *KLF2* and *KLF4*^51^. In the present study, *KLF2* and *KLF4* mRNA expression levels were expectedly upregulated 24h after laminar shear stress without any change observed between the non-canonical autophagy genotypes. Our measurement of the intracellular expression of the Hb-⍺ chaperone protein AHSP, which mirrored that of Hb-⍺, would favor a post-transcriptional degradation or regulation of Hb-⍺ in non-canonical autophagy deficient endothelial cells^52^. Furthermore, our data indicate that proteins involved in LANDO and receptor recycling pathways such as Rubicon, the WD domain of ATG16L1, and VPS29 were equally implicated in the regulation of shear stress-induced Hb-⍺ protein upregulation. Fundamental studies assessing reduction and oxidation of heme-associated iron carried by Hb-⍺ alongside a dynamic exploration of transcriptional and post-transcriptional regulation of *HBA1* related genes/proteins will be needed and may better define how Hb-⍺ expression levels are balanced during laminar shear stress but were beyond the scope of the present study.

Previous studies have indicated that deficiency in Rubicon, a negative regulator of canonical autophagy, displayed beneficial effects on cardiometabolic events including aging, lipopolysaccharide-induced stroke or cardiac pressure overload. Those results supported the hypothesis that canonical autophagy deficiency could be causal or primarily contributive to CVD^14^. A similar CV phenotype observed in mice deficient for Rubicon or the WD40 domain of ATG16L1 provide evidence that LANDO deficiency may initiate endothelial and cardiac dysfunction. In contrast with earlier negative data published using other mouse backgrounds, our use of a pure C57bl6 background, which is known to be sensitive to detect the onset of CV events, support our methodological approach^53^. Furthermore, our results emphasize a new layer of complexity regarding Rubicon function depending on its cell-type expression. Our observations that tissue-specific Rubicon deficiency could alter cardiac functions in vivo in endothelial cells at an early age, and in cardiomyocytes in older mice, are in line with data published by Yoshimori and colleagues, displaying a role of age-dependent loss of Rubicon in adipose tissue on metabolic disorders^54^.

While the circumstances in humans under which endothelial dysfunction initiates heart failure remain unclear, many clinical trials are testing whether therapies targeting endothelial cells could improve cardiac functions and CVD outcomes^55,56^. Peripheric endothelial dysfunction detected non-invasively by FMD in human brachial arteries have shown to be strongly correlated with coronary artery dysfunction assessed invasively by cardiac catheterization^57^. Both brachial and coronary endothelial dysfunctions precede clinical manifestations of CVD in humans and have regained interest regarding their prognosis values^55^. Interestingly, Rubicon joins the short list of 6 targets know so far, for which a sole genetic modification in endothelial cells were sufficient to induce cardiac dysfunction at baseline, without any nutritional, chemical or surgical cardiac dysfunction exacerbations^58^. Among those targets, mice with endothelial depletion of PECAM-1 exhibited LV dilation and systolic dysfunction, mainly explained by a lack of paracrine communication between endothelial cells and cardiomyocytes^59^.

In humans, heart failure with preserved ejection fraction (HFpEF), a major public health challenge which mortality is increasing in the general population, has been lately observed in younger patients <65 years of age and associated with impaired coronary microvasculature dysfunction^31,33,60,61^. Since obesity, diabetes and hypertension are the most prevalent comorbidities observed in HFpEF patients at higher risk of mortality, systemic inflammation has been hypothesized as the main trigger for endothelial dysfunction during the pathogenesis of HFpEF^32^. In the present study, no comorbidities, tissue fibrosis, systemic inflammation or immune cells infiltration were associated with endothelial and cardiac dysfunctions induced by non-canonical autophagy deficiency at baseline levels. Future studies exploring several nutritional aggravations of cardiac dysfunction will be necessary to better mimic human HFpEF and define whether non-canonical autophagy deficient mice could be valuable for pre-clinical assessment of novel therapeutic targeting endothelial dysfunction against heart failure.

Importantly, we identified many genetic variants with effects on genes related to LANDO and receptor recycling pathways which were associated with multiple traits of CV events in 2 independent cohorts, representing the general adult population in the UKB and younger individuals in the YFS. Our genomic analyses highlight genetic variants in genes involved in LANDO and receptor recycling that are associated with either increased risk or beneficial effects on CV parameters.

Our study is not without limitations. The UKB study has a well-documented healthy bias, meaning that our results likely underestimate the true effect on CV parameters^62^. Our genomic analysis began in the YFS with a list of 10 genes related to the LANDO pathway that was not sufficient to detect significant genetic associations with one CV trait. Our primary focus for this analysis was then on gene-based associations with a set of genetic variants in 42 genes selected for their potential implication in the LANDO and receptor recycling pathways. These gene-based findings have been experimentally validated in vitro, providing stronger supporting evidence. However, to improve our approach, the minimal number of genes, genetic variants, tissues, CV traits, the cohort size, the pathological and clinical features of the cohort, along with the optimum combinations should be further tested and fine-tuned. Only a few genes related to LANDO, whose functions are completely independent of canonical autophagy have been reported so far. Future studies exploring each genetic variant described in the present study in other ancestry-specific populations and their functional consequences at a protein level promise to improve resolution to identify independent components of non-canonical autophagy pathways. Furthermore, ongoing (CRISPR)-cas9-based screens in different cell types will better delineate which genes could be uniquely implicated in LANDO and CV disease.

## Conclusions

Our study identifies LANDO deficiency in endothelial cells as a pathway that fuels endothelial and cardiac dysfunction. The discovery that LANDO-related proteins regulated endothelial VEGFR2 and intracellular Hb-⍺ under shear stress offers new mechanistic insights and potential target for therapeutic intervention on early hallmarks of CV disease.

## Acknowledgments

We would like to thank the Hartwell Center at St. Jude for total RNA next-generation sequencing and Shahinur Alam, Richard Cross and Greig Lennon (SJCRH) for technical assistance, and Emilio Boada Romero (SJCRH), Zhenrui Li for thoughtful insights and discussions.

## Expanded Methods

### Experimental animals

The characterization of all genotypes was performed using genomic DNA and PCR with the primers listed in Table S10.

### Endothelial function measurement in vivo

To mimic flow-mediated dilatation measurement in the brachial artery in humans, endothelial-dependent vasodilatation assessment in mice femoral artery was adapted and developed from a method published previously^1–3^. The animal procedures were approved by the St Jude Children’s Research Hospital Committee according to guidelines. Males 8-12 weeks-old from C57Bl6/6N background mice were anesthetized with 2–3% isoflurane in an induction chamber and maintained with 1.0–1.5% isoflurane delivered via 100% O_2_ mask inhalation. The mice were placed in supine position on a controlled heating pad. The experimental setup is depicted in Figure S1A. The body temperature was maintained at 37°C and electrocardiogram limb electrodes were placed. After gently removing the fur in the left hindlimb region, pre-warmed ultrasound transmission gel was applied in the proximal inner thigh, and an ultrasound probe was manually aligned to the visible femoral artery as seen in Figure S1B. A high-frequency ultrasound device (Vevo 3100, Visual Sonics, Toronto, Canada) was used for ultrasound imaging with a high-frequency linear array probe MX-700 (29–71MHz). To properly image the femoral artery and identify the pulsatile blood flow pattern, the Duplex ultrasound mode was adjusted to the following parameters: B-Mode: Zoom 1.3x; Color-Mode: Beam Stream +15°; PW-Mode Frequency: 3-5Hz; Gate size: 0.26-0.30mm; PW-Mode Doppler-angle: -18-23°; Sensitivity:4. was adapted by using a Medium Occlusion cuff (#OCC-M, Kent Scientific Corporation, Torrington, CT, USA), which was placed around the lower limb and attached to a non-invasive blood pressure CODA® monitor (Kent Scientific Corporation, Torrington, CT, USA) as seen in Figure S1A. A special version of the CODA monitor has been developed by Kent Scientific to automatically induce a stable and reliable cuff occlusion at a pressure of 300mHg for 5min then to automatically release this pressure for 10min. Femoral artery occlusion in B-mode as seen in Figure S1B was confirmed in Color-Mode and PW-Mode. Measurement of femoral artery diameter were continuously recorded for 3 min every 30 seconds and analyzed using VEVO Lab ultrasound imaging analysis software, with VEVO Vasc Module (Visual Sonics, Toronto, Canada). Flow Mediated dilation (FMD) was determined as Δ% in average diameter change during reactive hyperemia as compared to baseline pre-ischemic values: [(Diameter post-ischemic – Diameter baseline) ⁄ Diameter baseline] *100. All diameter readings were taken at end diastole to limit the influence of potential differences in vascular compliance on diameter measurements.

### Conventional echocardiography

Transthoracic echocardiography was performed using a high-frequency ultrasound device (Vevo 3100, Visual Sonics, Toronto, Canada) equipped with a linear array probe MS-550D (25–55MHz). During the procedure, the mice were anesthetized with 2–3% isoflurane in an induction chamber and maintained with 1.0–1.5% isoflurane delivered via 100% O_2_ mask inhalation. The mice were placed in supine position on a controlled heating pad. The body temperature was maintained at 37°C and electrocardiogram limb electrodes were placed. The anterior chest and abdomen of mice were depilated using Nair depilatory cream (Church and Dwight Co., Ewing, NJ, USA). The heart rate, respiration frequency, and body temperature were continuously monitored. Heart rate was maintained at a maximum of 430 beats per minute. All ECG measurements were the average of 3 cardiac cycles.

The following parameters were measured from the LV mid-papillary level in the parasternal short-axis view with 2D M-mode imaging: interventricular septum thicknesses at end diastole and end-systole (IVSd and IVSs, respectively); LV internal diameters at end-diastole and end-systole (LVIDd, LVIDs); LV posterior wall thicknesses at end-diastole and end-systole (LVPWd, LVPWs); and stroke volume (SV), which was derived by the formula: EDV-ESV, where EDV and ESV are the end-diastolic and end systolic volumes. The LV systolic function was estimated by the ejection fraction (LVEF), which was derived by the formula: SV/EDV. All imaging and measurements were performed by a senior imaging technician who was blinded to treatment arm.

### Tail-cuff blood pressure recordings

The CODA noninvasive blood pressure acquisition system for mice (Kent Scientific, Torrington, CT, USA) was used for all tail-cuff measurements. This system uses Volume Pressure Recording (VPR) to detect blood pressure based on volume changes in the tail. CODA system was factory calibrated, and standard settings and recommendations were used as follows. Patency of the occlusion and VPR cuffs was checked routinely before the start of the experiments. The blood pressure measurement experiments were conducted in a designated quiet area (22±2°C), where mice acclimatized for a 1-hour period before experiments began. For all, mice were encouraged to walk into the restraint tubes, which end holders were adjusted to prevent excessive movement. The occlusion cuff was placed at the base of the tail and the VPR sensor cuff was placed adjacent to the occlusion cuff. Heating pads were preheated to 33 to 35°C. The mice were warmed for 5 minutes before and during blood pressure recordings. To measure blood pressure, the occlusion cuff is inflated to 250mm Hg and deflated over 20s. The VPR sensor cuff detects changes in the tail volume as the blood returns to the tail during the occlusion cuff deflation. Each recording session consisted of 5 to 5 inflation and deflation cycles per set, of which the first 2 cycles were “acclimation” cycles and were not used in the analysis, whereas the following cycles were used.

### RNA isolation and bulk RNA seq

Mouse cardiac LV apexes were collected and stored in RNA later (Ambion, Cat#7023, TX, USA). Approximately 20mg of cardiac tissue was dissociated in RLT buffer (Qiagen, Germantown, MD, USA) with a hard Tissue Homogenizing Mix 2.8 mm Ceramic beads using a Bead Ruptor 24 (OMNI international, Kennesaw GA, USA) programmed for cardiac tissues: 4 cycles of 45s at 5.65m/s, with 30s dwell time between cycles. The tissue homogenate was centrifuged for 15 min at 21,000 g at 4 °C then supernatants were carefully transferred into new tubes. Total RNA was extracted using RNeasy Fibrous Tissue Mini Kit (Qiagen, Cat#74704, Germantown, MD, USA).

RNA was quantified using the Quant-iT RiboGreen RNA assay (ThermoFisher) and quality checked by the 4200 TapeStation High Sensitivity RNA ScreenTape assay (Agilent) prior to library generation. Libraries were prepared from total RNA with the TruSeq Stranded Total RNA Library Prep Kit according to the manufacturer’s instructions (Illumina, PN 20020599). Libraries were analyzed for insert size distribution using the 5300 Fragment Analyzer NGS fragment kit (Agilent). Libraries were quantified using the Quant-iT PicoGreen ds DNA assay (ThermoFisher). Paired end 100 cycle sequencing was performed on a NovaSeq 6000 (Illumina).

Paired-end RNA-seq reads were aligned to the mouse mm10 reference genome using STAR^4^, with gene-level quantification performed by RSEM^5^, focusing on protein-coding genes with appreciable expression levels. Differential expression analysis was conducted with edgeR. Low-expressed genes were filtered out, and normalization factors were applied to adjust for differences in library size. Data were transformed with voom to stabilize the mean-variance relationship, and contrasts were applied to identify differentially expressed (DE) genes. Linear modeling and eBayes moderation were used to enhance statistical power in detecting expression differences. Following differential expression analysis, volcano plots were generated, with a significance cutoff of p-value < 0.05 and fold change > 2 to highlight DE genes. Pathway enrichment was performed with GSEApy on KO vs WT contrasts using two approaches: Enrichr (over-representation of up/down DE lists) and pre-ranked GSEA. Significant pathways were defined as adjusted p < 0.05 (Enrichr) or FDR q < 0.25 (pre-ranked GSEA), with NES sign indicating direction (positive = KO, negative = WT).

### Blood chemistry

For blood chemistry analysis, blood was collected from the venous sinus via retro-orbital bleeding after euthanizing the mice and allowed to clot for 30-60 minutes. The samples were centrifuged at 5700 rpm for 10 minutes to harvest the serum and pipetted into a Horiba bio cup. The serum samples were diluted (1:2) into ultrapure Millipore water. A Horiba ABX Pentra 400 chemistry analyzer was used to detect albumin, alkaline phosphatase, alanine aminotransferase, amylase, BUN, calcium, creatinine, globulin, glucose, phosphorous, potassium, sodium, total bilirubin and total protein in serum using manufacturer-provided reagents per the manufacturer’s protocol.

### Light sheet microscopy of cardiac vasculature leakiness

For light sheet experiments, mice received intravenous (i.v) tail vein injection 10min before sacrifice with 5ug of anti-mouse CD31 (clone MEC13.3) conjugated with AlexaFluor (AF)-647 (Biolegend #102516) and with 10ug 70kDa-Dextran conjugated with Tetramethylrhodamine (ThermoScientific #D1818) in PBS in a total volume of 100µl per animal. After sacrifice, hearts were excised and subjected to following tissue processing as described in Figure S4A.

#### Tissue clearing

Perfusion fixed hearts were immersed in 4% paraformaldehyde (w/v) (Polysciences; cat#18814-20) in PBS overnight at 4°C. Samples were dehydrated through a series of methanol/ddH_2_O solutions (50%, 75%, 100%, 100%), each for 1 hour with shaking, and then delipidated overnight in a solution of 66% dichloromethane/33% methanol. Samples were then washed twice in 100% methanol and bleached overnight at 4°C in freshly prepared 5% hydrogen peroxide/methanol. Finally, samples were rehydrated through a series of methanol/ddH_2_0 solutions (75%, 50%, PBS), each for 1 hour with shaking, followed by two, one-hour washes of PBS+2%Triton. All steps were performed at room temperature except as noted.

#### Immunolabeling

All steps were performed at 37°C. Cleared hearts were blocked in a solution of 10% goat serum, 2% Triton X-100, and 20% DMSO in PBS for 24 hours with shaking and then incubated for 4 days with directly conjugated primary antibodies at CD31 directly labeled with AlexaFluor647 (Biolegend #102516) is and used at 5µg/mL in 3% goat serum, 2% Triton X-100, 10% DMSO and 1% Glycine in PBS. Samples were then washed 5x in PBS plus 0.1% Tween (v/v) over the course of 24 hours.

#### Refractive index matching

Immunostained hearts were dehydrated through a series of methanol/ddH_2_O solutions (50%, 75%, 100%, 100%), each for 1 hour with shaking. For refractive index matching, samples were incubated with ethyl cinnamate (ECi) for at least 4 hours prior to imaging. (Tubes were filled to the brim to prevent oxidation). All steps were performed at room temperature.

#### Light sheet fluorescence microscopy

RI-matched hearts were imaged on a Zeiss Light Sheet 7 with two, pco.edge mono sCMOS cameras using Zen software. Samples were suspended in the ‘5x’ imaging chamber and glued to the sample holder using Loctite Super Glue Professional. Imaging was performed using simultaneous excitation with 488 nm and 647 nm lasers through 5x/0.1 NA light sheet forming objectives with pivot scan, dual side excitation, and online dual-side mean fusion. Emissions were collected with a 5x/0.16 NA detection objective and passed through 575-615 and LP 660 emission filters. Camera exposure time was 100 msec and pixel size was 0.938 µm/pixel. Z-steps used 4.0 µm spacing based on a predicted gaussian sheet waist of 8.42 µm. Hearts were oriented such that the detection objective faced the heart’s anterior surface. Volumetric tiles were acquired in ‘reading’ mode with 10% overlap and saved in CZI format.

#### Image stitching and rendering

CZI files were converted into Imaris format using the Imaris File Converter 9.9.4 (Bitplane) and then stitched using Imaris Stitcher 9.9.4 (Bitplane). 3D renderings were created in Imaris 10.0.1 (Bitplane), as described in Figure S4B.

### Image registration, pre-processing and analysis

To register the 3D cardiac volumes, we used the semi-automated software package BigWarp^6^ with 30 cardiac morphological landmarks over 3 different Z-stacks (10 in the Left Ventricle, 10 in the intraventricular Septum and 10 in the right ventricles as described in Figure S4C.

Samples TBB3339 and MAC1721 were used as the fixed reference hearts. Rather than warp whole image volumes, we transformed the coordinate space of each cardiac volume using these 30 manually selected control points under a thin plate spline transformation. Control points were selected in-place for all hearts, however the registration transformations for the two anatomically inverted samples (TBB3341, TBB3623) were calculated using points reflected over the x-axis. For each of the 30 control points, there were minor variations in the exact (x,y,z) reference coordinate used between sample registrations (reference point SD = 141 µm for MAC, and 90 µm for TBB in any given direction). To standardize the sub-volume selection across all cardiac samples, the mean of each reference coordinate, taken across all registration pairs, was used as a fixed landmark. These landmark coordinates were then warped into the corresponding anatomical space of each cardiac volume. 512 x 512 x 128 voxel (∼ 480 x 480 x 514 µm^3) regions of interest (ROIs) were cropped around each landmark, then up-sampled by 4x along the z-axis using B-spline interpolation to create a roughly isotropic volume. Colocalization of the fluorescently-labeled 70kDa-Dextran and anti-CD31 was measured through mutual information and Pearson’s correlation between imaging channels, both estimated using random subsamples of 51 x 51 x 51 contiguous voxels, bootstrapped 1000 times for each ROI (see Code availability statement).

#### Whole heart visualization

To enhance the vasculature signals and visualize the whole hearts (Figures 3A and 3D), we used a method called Ranking Responses of Robust Path Operators (RORPO). This technique is a low-level curvilinear detection filter that works using paths, a set of connected pixels, and is non-linear and global^7^. Before running RORPO, the original images were downsampled by 8x along all three dimensions to reduce computational costs.

The RORPO algorithm takes in the following parameters - scale (30.0) which specifies the unit length of path being used for detection, dilation size (0) which is the limit set for tolerating noise in the path, number of scales (3) that tells you the number of times to scale the unit path detector in order to create a list of scaled unit detectors and the scaling factor (1.5) which is the factor for creating a geometric sequence of scales for each scaled unit detector in the list. The values used for our experiments are in parenthesis. We used the Python bindings for RORPO, pyRORPO (Github repo link available in the Code availability statement); and the final images were rendered using the software Imaris.

### Isolation and culture of endothelial cells from murine lungs

Three–6-week-old mice were used for pulmonary microvascular endothelial cells (PMVEC) isolation as previously described with modifications^8,9^.

#### Isolation and plating lung cells

Ten minutes before sacrifice, heparin sodium (25µl/mouse, 1000U/ml, Fresenius Kabi, Cat#504013, IL, USA) was injected in the hindlimb intramuscularly. After transcardial perfusion (25G needle) of 15ml ice-cold serum-free DMEM (ThermoScientific; Cat#11965092), the lungs, one lob at a time, were collected in 20ml of ice-cold basal medium (DMEM, 20% FBS, 25mM HEPES, 1x PEST), quickly agitated to remove traces of blood, washed in HBSS (ThermoScientific; Cat#14175095), shredded with sterilized scissors in an Eppendorf tube then digested in a 10cm dish with collagenase I (3m/ml in HBSS; Gibco, Cat#17100-017) at 37°C for 45min under gentle agitation 80rpm. The minced tissue was mixed with 20ml basal medium to stop the digestion and further triturated (10 passages in a 20ml Syringe attached with 20G needle). Cell suspension was filtered through a 70-µm cell strainer then centrifuged at 400g for 8min. Cell pellets were resuspended in 20mL complete medium (45mg endothelial cell supplement, 1x NEAA, 1x Sodium pyruvate, 45mg Heparin Sulfate in basal medium), transferred in T75 flask CellBind treated (Corning, Cat#3290) and maintained at 37°C in a humidified 5% CO_2_ chamber. The following day, the attached cells were washed twice with 10mL of HBSS then 20mL complete medium were added, changing the media every 2 days.

#### Isolation and plating CD31 cells

When 95% confluency was reached, cells were washed twice with HBSS, detached with Accumax (StemCellTechnologies, Cat#07921) for 5min at 37°C, resuspended in basal medium then pelleted at 400g for 4min. Cell pellets were resuspended in 2ml basal medium then transferred in a 5ml polystyrene round bottom tube. An appropriate volume of sheep anti-rat Dynabeads (30ul per 2ml cell suspension) pre-coated overnight with purified rat anti-mouse CD31 antibody (BD; Cat#553370; clone MEC13.3; 0.5mg/ml) was used. The cells were incubated 10min at room temperature on an end-over-end rotator at 30rpm then applied to a magnetic separator for 2min. According to the manufacturer’s instructions, the cells suspension was washed 5 times with 3ml of basal medium. The CD31 positive enriched cells were resuspended in 20ml complete medium, transferred in a T75 flask CellBind treated (Corning, Cat#3290) and maintained at 37°C in a humidified 5% CO_2_ chamber. The following day, the attached cells were washed twice with 10mL of HBSS then 20mL complete medium were added, changing the media every 2 days.

#### Isolation and plating double positive CD31-CD102 positive cells

When 90% confluency was reached, cells were washed, detached and resuspended as described above for CD31 positive enriched cells. Sheep anti-rat Dynabeads (30ul per 2ml cell suspension) pre-coated overnight with purified rat anti-mouse CD102 or ICAM2 antibody (BD; Cat#553326; 0.5mg/ml) were used. After the magnetic separation and washing steps, double positive CD31-CD102 positive cells or PMVEC, were resuspended in 20mL complete medium then transferred in 2x 10cm dishes precoated 1h with 0.2% gelatin (Sigma, Cat#G1393 diluted in PBS). The following day, the attached cells were washed twice with HBSS then 10mL complete medium were added, changing the media every 2 days.

### Culture of human aortic endothelial cells

Human aortic endothelial cells (HAEC) were purchased at ATCC (Cat#PCS-100-011), plated on fibronectin coated plates (1µg/cm^2^, Corning; Cat# 354008) and maintained in EBM-2 medium (Lonza, Cat# CC-3156) completed with Endothelial Cell growth medium-2 bullet kit (Lonza, Cat# CC-3162) in a humidified 5% CO_2_ chamber at 37°C. Medium was changed every 3 days. The cells were used mainly in passage 4 for the study.

### Generation of *RUBCN* KO in HAEC

*RUBCN* KO in HAEC were generated by CRISPR-Cas9 technology using the LentiCRISPRv2GFP system (Addgene, 82416), where the GFP was replaced by RFP and carrying gRNA targeting human RUBCN (5’–GCTAAGTGACGCTCATGTCA-3’). The transduced cells (RFP+) were cell sorted by flow cytometry (St. Jude Flow Cytometry and Cell Sorting Shared Resource) into five days post lentiviral transduction. Cells were screened for the desired targeted modification (out-of-frame indels) via targeted deep sequencing using gene specific primers with partial Illumina adapter overhangs (CAGE3132.RUBCN.F–5’TGTTTCTGATCTGCCTTCACCCTCC-3’ and CAGE3132.RUBCN.R–5’ GGGCTACATCCTGCTGAGCTCCAGGG-3’, overhangs not shown) as previously described^10^.

### *VPS29*, *ATG3*, *PIK3C2A* knockdown by using siRNA approach in HAEC

A total of 200,000 HAEC were seeded in antibiotic-free medium 24 h before transfection to achieve 60–80% confluency at the time of transfection. For each 100-mm culture dish, *VPS29*, *ATG3* or *PIK3C2A* siRNA (final concentration 100pmol, siGENOME Human siRNA, SMARTPool, Horizon Discovery) and Lipofectamine RNAiMAX (30 µl per dish, Thermo Fisher, cat#13778150) were separately diluted in Opti-MEM Reduced-Serum Medium (ThermoFisher, cat#31985070). Both tubes were inverted by hand for 2–3 s. Then, the diluted siRNA was combined and incubated with the diluted Lipofectamine RNAiMAX reagent for 15 min at room temperature to form siRNA–lipid complexes. The mixture was added dropwise to cells, followed by gentle swirling. Cell culture medium should be refreshed after 6h transfection at 37 °C in a 5% CO2 humidified atmosphere to avoid cytotoxicity. Control transfections included non-targeting siRNA (WT, Horizon Discovery). Finally, the cells were collected after 48h for VPS29, ATG3, PIK3C2A protein expression detection and further shear stress studies.

### Shear stress studies

#### For immunofluorescence assays

Murine PMVEC or HAEC were grown to confluent monolayers in 0.2% gelatin pre-coated µ-slide-IV 0.4-luer (Ibidi, cat#80606). Laminar shear stress was generated through a parallel flow system (Ibidi, cat#10902) at 5dyn/cm^2^. The system was maintained at 37 °C and ventilated with 95% humidified air containing 5% CO_2_.

#### For western-blotting, nitrites, protein S-nitrosylation and proteomic assays

Endothelial cells were seeded in the periphery of modified 100-mm culture dishes that were made by bonding the bottoms of 60-mm culture dishes upside down into the center of 100-mm culture dishes using clear epoxy resin (Gorilla, 0.85oz syringe cat#4200101-2,) as previously described and adapted (Dela Paz et al. JCell Science 2012; Choi et al. CircRes2017). Dishes were sterilized under UV light for 30 minutes and then coated with 0.2% gelatin for 1h before plating the cells in their complete medium. When 95% confluency was reached, HAECs and murine PMVECs were starved for 24h in a starvation medium (DMEM-F12 without phenol red, HEPES 25mM, PEST 1%) completed either with 0.5% FBS for the HAEC or with 0.5%BSA for the PMVEC. Confluent cell monolayers were subjected to orbital shear stress for 24 hours in the starvation media using an orbital shaker at 210rpm (Microyn, Cat# MT-201-BD) in a humidified 5% CO_2_ chamber at 37°C.

### Recycling assay

Receptor recycling assays on murine PMVEC or HAEC were performed as previously described and adapted^11,12^. EC were plated on 0.2% gelatin pre-coated 4-well chamber slide ibiTreat 1.5m polymer coverslide (Ibidi, cat#80426) at a density of 50000 cells per well and maintained in their complete medium. Static conditions or laminar shear stress was generated by orbital shaker for 24h. Any unspecific binding to Fc receptors were avoided by incubating the cells in pre-warmed recycling medium (EGM-2 with 10% donkey serum; Sigma; cat# S30-M) for 15 min at 37°C. Then, monoclonal antibody against VEGFR2, PECAM1, VE-Cadherin or Piezo-1 (Table S11) were added to the cells in EGM-2 with 1% donkey serum for 1h at 37°C. Next, the primary antibody was stripped off the cell surface using 2 washes of ice-cold stripping medium (EGM2 at pH 2.0) of 1min each then 2 washes of PBS. Cells were incubated again in the recycling medium for 1h at 37°C under static or laminar shear stress conditions, which was then provided with fluorescently labelled secondary antibody (Table S11) in 1% donkey serum for 1h at 37°C. Cells were again stripped off the cell surface by acid stripping. Cells were then fixed with 4% paraformaldehyde for 15min at room temperature, washed 3x with PBS, then incubated with Hoechst 33342 nucleic acid staining solution (Table S11). Slides were imaged with Marianas spinning disk confocal microscope (Intelligent Imaging Innovations) equipped with a 100X 1.3NA objective and EMCCD camera. Images were analyzed either using Imaris 10.0.1 (Bitplane) or Slidebook 7 software (Intelligent Imaging Innovations). The receptor recycling assay was quantified as the fluorescent signal divided by the total number of cells or nuclei present in the field to generate a measurement of fluorescent area per cell.

### Determination of NO metabolites and protein S-nitrosylation

For nitrites and nitrates analysis, after shear stress or static exposure, the cell media were quickly cleared by centrifugation, snap-frozen and conserved at -80°C. Nitrate and nitrate levels were determined using a fluorimetric assay kit (Cayman; Cat#780051, Ann Arbor, MI, USA). Protein S-nitrosylation was detected using a S-Nitrosylated Protein Detection kit or Biotin Switch assay (Cayman, Item# 10006518, Ann Arbor, MI, USA).

### Immunoblot analysis

Protein extract from human aortic endothelial cells or murine PMVEC were prepared in an isotonic lysis buffer containing Tris-HCl (pH 7.5; 50 mmol/L), NaCl (150 mmol/L), NaF (100 mmol/L), Na_4_P_2_O_7_ (15mmol/L), Na_3_VO_4_ (2mmol/L), leupeptin (2mg/mL), pepstatin A (2mg/mL), trypsin inhibitor (10mg/mL), Phenylmethylsulphonyl fluoride (PMSF;44mg/mL), and Triton X-100 (1%vol/vol), phosphatase inhibitor cocktail (Roche, USA), left on ice for 10minutes, and centrifuged at 10 000g for 10 minutes (Fleming et al. CircRes 2001). For flow experiments, endothelial cells were directly lysed in 500ul of 2x SDS sample buffer (Biorad). The lysates were heated at 99°C for 10 min followed by centrifugation at 15000g for 10 min and the supernatants were collected. Proteins were separated by SDS-PAGE on 4-20% gradient gels (BioRad) and transferred to nitrocellulose membrane, which was blocked with 3% BSA-TBS and incubated with specific primary antibody. An Odyssey Fc imager (LI-COR) was used as detection system. Proteins were detected with the following antibodies total VEGFR2 (CST,2479), Y1175p-VEGFR2 (CST,2478), total eNOS (CST,9572), Ser1177p-eNOS (CST,9571) for HAEC and Ser1177p-eNOS (BD,612392) for murine PMVEC, Pecam1 (CST, 77699), CXCR4 (R&D,MAB21651-100), Actin (Santa-Cruz,sc-47778-HRP), Hemoglobin alpha (Abcam,ab174536), Atg16L1(MBL,PM040), Rubicon (CST,8465), AHSP (Abcam, ab180861).

### Surface protein isolation

For surface protein isolation summarized Figure S5B, HAEC were grown on in the periphery of modified 100-mm culture dishes gelatin coated plates to confluency. The Pierce™ Cell Surface Protein Isolation Kit (ThermoScientific, #A44390) was used according to the package instructions with the following modifications. Following shear stress or static exposure, cells were washed twice quickly with iced-cold HBSS with Ca^2+^ and Mg^2+^ (ThermoScientific, # 2402117) before incubating for 30min at 4°C with 3m/dish EZ-Link Sulfo-NHS-SS-Biotin (final concentration 0.25mg/ml in iced-cold HBSS with Ca^2+^ and Mg^2+^). Cells were washed twice with cold TBS1X furnished in the kit then detached with 1ml Accumax (StemCell Technologies, #07921), then blocked with stopping buffer (0.5% FBS in TBS1X). Cells were pelleted, washed twice in cold TBS1X then lysed in 250μL lysis buffer and sonicated briefly, 5 pulses at the lowest setting (10%) for 1 s each on ice (+ add Sonicator info). Lysates were incubated on ice for an additional 30 min before spinning down at 15000g for 5min at 4°C. Then, 50μL of cell lysate was separated for the whole-cell lysate fraction and 200μL of cell lysate was added to each column prepared with 400μL NeutrAvidin Agarose slurry (ThermoScientific #29201) and placed in an end-over-end rotator for 1h. The first flow through fraction was collected as the intracellular fraction or unbound fraction, then the column was washed 3 times. 200μL of Elution buffer with Dithiothreitol (DTT) was then added to the columns, which were rotated for 1 h at room temperature to elute the biotinylated proteins or the surface fraction. For western blotting, 15μL of each fraction was mixed with 5μL 4x Laemmli, with a total of 20μL loaded on the gel after boiling for 5 min.

### Proteomic analysis

Cells were lysed and digested into peptides. After desalting, the peptides are labeled with TMT reagents. The reagents consisted of three groups: an amine-specific active ester group that reacts with free alpha N-termini of peptides and the epsilon amino groups of lysine residues, a balance group, and a reporter group. The isobaric labels, defined by a different distribution of isotopes (^13^C, ^15^N) between the reporter and balance groups, allowed each labeled peptide to coelute during chromatography and to ionize at the same mass-to-charge in MS scans. The labeled samples were equally mixed and further fractionated by neutral pH reverse phase liquid chromatography. In general, 10 fractions were collected and further analyzed by low pH reverse phase LC-MS/MS. During ion fragmentation, the TMT regents were cleaved to produce reporter ions for quantification.

The collected data were searched against a database to identify peptides. While the peptides were identified by MS/MS, the quantification was achieved by the fragmented reporter ions in the same MS/MS scans. The peptide quantification data was then corrected for mixing errors and summarized to derive protein quantification results. Differential expression was performed with limma (empirical-Bayes moderated linear models), using KO vs WT contrasts separately for the static and flow experiments; results were visualized with volcano plots, highlighting proteins with p < 0.05 and |log2FC| ≥ 0.58 (∼1.5-fold).

### Immunofluorescence of hemoglobin alpha

Endothelial cells were grown to confluent monolayers in 0.2% gelatin pre-coated µ-slide-IV 0.4-luer (Ibidi, cat#80606). Directly after shear stress or static exposure, cells were fixed in 4% PFA in PBS for 10 min at room temperature. Fixed cells were permeabilized in a blocking buffer (0.5% triton-X100, 4%BSA, 0.01% Sodium Deoxycholate in PBS) for 45 min at room temperature. The permeabilized cells were then incubated with 5 μg/ml Hemoglobin subunit alpha antibody (Abcam, Ab174536) prepared in a diluent buffer (blocking buffer diluted 1:2) overnight at 4°C. Cells were washed for 5 min, three times in PBS and then incubated in Alexa plus 647 anti-rabbit secondary antibody 1:50, and Cell mask (Table S11) 1:5000 for 1 h at room temperature, protected from light. Cells were then washed for 5 min, three times before Hoechst 33342 nuclear staining solution. Slides were imaged with Marianas spinning disk confocal microscope (Intelligent Imaging Innovations) equipped with a 100X 1.3NA objective and EMCCD camera. Images were analyzed using Imaris 10.0.1 (Bitplane) and quantified by measuring the total of Hb-⍺ positive staining divided by the total number of cells per 100× field of view.

### Genetic variants’ selection for genomic analysis

A selection of genetic variants in 42 genes related to LANDO pathway was based on their presence in the coding regions or the regulatory regions which was assessed in Ensembl using Variant effect predictor (VEP). In VEP, two set of filters were applied: (1) Variant’s consequences: 3’UTR, 5’UTR, missense, regulatory region, splice region, TF binding site.

Intronic, non-coding, intergenic, synonymous, downstream, upstream consequences were filtered out; (2) Genetic variants biotypes: CTCF binding sites, enhancer, promoter, open chromatin, protein coding, processes transcript. Non-sense mediated decay, retained introns, miRNA, LnRNA, other RNAs were filtered out. Secondly, we included genetic variants with significant changes observed in cis-expression quantitative trait loci (eQTL) from two public database in whole-blood^13^ (n=31684) and GTEX v8 across 10 neuro-cardiovascular tissues (n=838)^14^. We also included genetic variants with significant changes in protein quantitative trait loci (pQTL) in the cortical brain regions^15^ (n=330), the only pQTL database in neuro-cardiovascular tissues where genetic variants in LANDO pathway related genes loci were significantly associated with protein levels (until 2023). A brief description of the two eQTL and one pQTL datasets are found in Table S9.

We identified a total of 8143 genetic variants with significant changes in either the 2 cis-eQTL or the pQTL analyses (4168 genetic variants associated with an upregulation and 3975 associated with a downregulation of LANDO pathway related genes or proteins expression levels; Supplementary Excel Table 1A-1B).

### UK Biobank

The UK Biobank study (UKB) is a population-based study that has been described previously^16,17^. Briefly, between 2006 and 2010, approximately 500,000 individuals were recruited through 22 centres across the UK. Each participant underwent a physical examination, answered extensive questionnaires on their health, medication use and lifestyle and provided blood samples for genetic and biochemical analyses. Touchscreen questionnaires were followed up in some domains with in- person questionnaires (especially for medication use). A subset of the study was invited for a follow-up imaging visit. Informed written consent was provided by all study participants. This work was approved under the generic ethical approval for UK Biobank studies granted by the NHS National Research Ethics Service (approval letter dated 29 June 2021, Ref 21/NW/0157). This project used data from UK Biobank applications 71392 (PI. RJS). *UK Biobank (UKB) Phenotyping:* Cardiovascular (CV) events were based on self-report of ischemic heart disease (ISH, angina and/or heart attack, #6150), stroke (#6150) or venous thromboembolism (VTE, including deep vein thrombosis and/or pulmonary embolism, #6152). Ultrasound measurement of the carotid intima-media thickness (cIMT) was recorded at two angles on each side (#22671, 22672, 22674, 22675, 22677, 22678, 22680), and three measures were calculated: the average of 4 mean measures (2 for each of the left and right carotid arteries) was calculated for the mean (average of mean cIMT values [cIMT mean]); the maximum IMT (average of maximum cIMT values [cIMT max]), where the largest of the 4 maximum IMT measures was used; the mean of 4 maximum measures (2 for each of the left and right carotid arteries) was calculated [cIMT mean-max]), as described by Strawbridge et al.^18^. Cardiac MRI imaging, quality control and analyses have been described previously^19^. From this data, central and peripheral pulse pressure (#<u>12678</u> and <u>12676</u> respectively), left ventricle ejection fraction (#24103), left ventricle end systolic volume (#22422) and left ventricle end diastolic volume (#22421) were considered. The use of lipid-lowering or antihypertensive medication was self-reported (#6177 and 6153). Systolic and diastolic blood pressure (SBP and DBP, data field #4079 and 4080 respectively) were measured twice and the average computed. Prior to analyses, these measures were adjusted for hypertensive medication use (to estimate drug-naive values) as per Ehret et al.^20^ Body-mass index (#1001) was calculated by the central UKB team, from height and weight measurements. Waist and hip circumferences were measured (data fields #48 and 49 respectively), and waist: hip ratio calculated.

#### UKB Genotyping

Details of the blood storage, DNA extraction, genotyping, imputation and quality control have been described in detail elsewhere^21^. Briefly, the central UKB team conducted standard quality control on genetic data prior to and after imputation, and a clean dataset was provided to researchers. The central UKB team also computed pairwise relatedness and principle genetic components. For this study, only self-reported “white British” ancestry individuals were included and where related individuals were identified, only one of each pair were retained.

Single variant analyses: eQTLs were tested for association with ISH, stroke, VTE, SBP, DBP and cIMT mean, max and mean-max. cIMT were normalized by log transformation prior to analyses. Genetic association analyses were conducted using Plink 1.9 by logistic regression for dichotomic traits, and linear regression for quantitative traits, assuming an additive genetic model and adjusting for age, sex, population structure and genotyping chip. For pulse pressure and left ventricle measures, residuals were calculated and quantile normalized prior to analyses. The threshold for significance was set at P<1x10^-5^ (Bonferroni multiple testing correction divided by the number of independent variants tested). In total, 6891 variants were included in the analyses in up to 402,137 individuals.

#### Data availability

The UK Biobank study data is available by application to the central UK Biobank team (https://www.ukbiobank.ac.uk/enable-your-research).

### Young Finns Study

The Young Finns Study (YFS) is one of the largest prospective multicenter follow-up studies assessing CV risk factors from childhood to adulthood^22^. The study began in 1980 with 3,596 children and adolescents aged 3–18 years, randomly selected from five university hospitals in Finland (Turku, Tampere, Helsinki, Kuopio, and Oulu). The participants were regularly followed for over 40 years. The study was approved by the ethics committee of the Hospital District of Southwest Finland on 20 June 2017 (ETMK:68/1801/2017). All participants provided written informed consent, and the study was conducted in accordance with the Declaration of Helsinki. Data protection will be handled according to the current regulations.

#### Phenotypes

2007: Brachial artery ultrasound studies were performed to assess brachial endothelial function, the brachial artery baseline diameter and FMD using Sequoia 512 ultrasound mainframes (Acuson, Mountain View, CA, USA) with 13.0 MHz linear array transducer, as previously described^22,23^. Left brachial artery diameter was measured at rest and during reactive hyperemia to derive the percentage in arterial diameter relative to resting scan, and the highest of three measurements was chosen as the maximum FMD. Blood pressure from the right brachial artery was measured in the sitting position after a 5-minute rest with a random-zero sphygmomanometer (Hawksley & Sons Ltd, Lancin, United Kingdom) as described previously^24^. The average of 3 measurements was used in the analysis.

*2011:* The echocardiographic examinations were performed according to the American and European guidelines ^25,26^. Transthoracic echocardiograms were performed with Acuson Sequoia 512 (Acuson Mountain View, California, USA) ultrasonography, using a 3.5 MHz scanning frequency phased-array transducer. Analyses of the echo images were performed by a single observer. Both the sonographer and the observer were blinded to the subject’s details. Standard echocardiographic examinations were produced from the standardized image planes and modes: parasternal long and short axis in 2D and M mode and apical four chamber view^25^. Based on the recommendations to assess LV ejection fraction, the end-diastolic and end-systolic volume were measured from the apical 4-chamber view^25,26^. LV ejection fraction was calculated as follows: Ejection fraction (EF) = 100 × (LV end-diastolic volume-LV end-systolic volume) / LV end-diastolic volume.

#### Genotyping and genotype imputation

Genomic DNA was extracted from peripheral blood leukocytes using a commercially available kit and Qiagen BioRobot M48 Workstation according to the manufacturer’s instructions (Qiagen, Hilden, Germany). Genotyping was done using custom build Illumina Human 670 k BeadChip atWelcome Trust Sanger Institute. Genotypes were called using Illuminus clustering algorithm. Samples that failed Sanger genotyping pipeline quality control criteria (i.e., duplicated samples, heterozygosity, low call rate, or Sequenom fingerprint discrepancy) were excluded from the analysis. Similarly, samples with sex discrepancy, low genotyping call rate (<0.95) and possible relatedness (pi-hat>0.2) were excluded from the analysis. Short Nucleotide Polymorphisms (SNPs) were excluded based on Hardy-Weinberg equilibrium test (p ≤ 1e-06), failed missingness test (call rate<0.95) and failed frequency test (minor allele frequency<0.01). Overall, 546677 genotyped SNPs passed the quality control. Genotype imputation was performed using TOPMed Imputation Server and TOPMed r2 GRCh38 reference set.

#### Data availability

The YFS dataset comprises health related participant data and their use is therefore restricted under the regulations on professional secrecy (Act on the Openness of Government Activities, 612/1999) and on sensitive personal data (Personal Data Act, 523/1999, implementing the EU data protection directive 95/46/EC). Due to these legal restrictions, the Ethics Committee of the Hospital District of Southwest Finland has in 2016 stated that individual level data cannot be stored in public repositories or otherwise made publicly available. Data sharing outside the group is done in collaboration with YFS group and requires a data-sharing agreement with the understanding that collaborators will protect the data and not share it with any other parties. The list of all investigators that collaborate with the YFS group is displayed at the website of the YFS (http://youngfinnsstudy.utu.fi/). Investigators can submit an expression of interest to the chairman of the data sharing and publication committee (prof Mika Kähönen, Tampere University) in the case of clinical data and in genomic data to professor Terho Lehtimäki (Tampere University).

#### Statistical analysis

The association analysis was conducted using PLINK 2^27^ using additive models adjusted for age, sex and first five principal components of the genetic data.

### Genetic association analyses

In cases of the same genetic variant being simultaneously associated with bi-directional changes in QTL in different tissues (up and downregulation), this variant was removed from analysis (911 removed in total).

*For the UKB genomic analysis* the raw data are presented in Supplementary Excel Table (Table 2A-2F). A combined analysis of the genomic data was run separately for each directional change in QTLs (upregulation or downregulation; Supplementary Excel Table 2A and 2B).

Genetic variants ID (rsID) were first harmonized between the QTLs and the UKB datasets. For stroke and ischemic heart disease (ISH) events, odds ratios (OR) were converted to beta values using log^10^(OR). A stringent threshold for significance was set at P<5x10^-8^, meaning that a genetic variant should have passed this threshold in any of CV parameter or phenotype). A total of 320 genetic variants passed this threshold (166 associated with an upregulation and 154 with a downregulation of the LANDO pathway related genes or proteins expression levels).

#### Summary of genetic variants-CV trait associations for a specified gene

A DataFrame with columns [’Gene’,’CVParameter’,’Genetic variants’,’Tissue’,’Directionality’,’PValue’,’Beta’] filtered rows for the chosen gene. Each association were plotted as a circle at (Parameter, Genetic Variants), with circle area proportional to statistical significance (size = -log^10^(pValue) × 100), fill color encoding the direction of effect from Beta (hotpink for β ≥ 0, deepskyblue for β < 0), and edge color encoding the tissue of origin. Genetic variant tick labels on the y-axis are colorized by Directionality (mapped via (’Up’:’darkblue’,’Down’:’darkred’), while x-axis ticks show the parameter order (As seen for example in Figure 6B). To facilitate the visualization of the association of genetic variants with CV trait for a specified gene, we introduced a rule of tissue prioritization where the 5 CV tissues (Heart left ventricle, heart atrial appendage, coronary artery, aorta and tibial arteries) had the highest priority. If a genetic variant was present in 5 of the CV tissues, the p-values and relevant scores were displayed in priority (even though present in other tissues). If a genetic variant was absent in 5 of the CV tissues, the second level of priority was given to the whole blood data. If a genetic variant was absent in 5 of the CV tissues or whole blood, a mention other tissues was attributed, referring then to pulmonary and cerebral tissues.

#### Aggregation of genetic variants-level evidence to gene-level signals and visualization

For each gene, we first grouped the input table (retaining QTLs metadata such as rsID, study_origin, tissues, gene_id, Directionality) and, across predefined CV traits p-value fields (cIMT_meanmax, cIMT_mean, cIMT_max, dbp, sbp, stroke, P_ish) computed a combined p-value per gene using Fisher’s method (scipy.stats.combine_pvalues). We appended both the raw combined p-values and their −log^10^ transforms, replacing infinities with the column maximum, and recorded the number of contributing genetic variants as Total_genetic variants. After filtering (Total_genetic variants ≥ 2), we plotted two panels—genes inferred from “Upregulated genetic variants” and from “Downregulated genetic variants.” Each gene–parameter association were drawn as a circle whose area scaled with statistical support −log^10^ (PValue); circle face color encoded the genetic variants burden using a continuous colormap (Reds) on Total_genetic variants, with a horizontal colorbar placed above the plot (as seen in Figure 6C).

#### For YFS genomic analysis

The raw data are presented in Supplementary Excel Table (Table 3A-3G). Genetic variants rsIDs were first harmonized between the QTLs and the YFS information using a total of 6600 genetic variants with significant changes in QTLs, which were genotyped in the YFS (Excel Supplementary Table 3H; Genetic variants_annotation in the YFS). Since the YFS was used as a validation dataset in our genomic analysis, the focus for this cohort was on similarity on effect sizes and directions. Therefore, we considered genetic associations in the YFS as significant, those with P-value/number of tests (where number of tests are only those significant in the discovery cohort, the UKB) and then as suggestive if P-value<0.05 and consistent beta-values. A total of 285 genetic variants passed this threshold (141 associated with an upregulation and 144 with a downregulation of the LANDO pathway related genes or proteins expression levels).

### Code availability statement

Landmark coordinates were transformed for registration using the BigWarp Fiji macro: https://github.com/saalfeldlab/bigwarp/blob/master/scripts/Apply_Bigwarp_Xfm_csvPts.groovy. ROI extraction and upsampling was done using the Insight Toolkit (ITK 5.4.0) for Python. The code for correlational analyses can be found here: https://github.com/panlilio/corrutil.

The “develop” branch of the RORPO github repository was used to enhance the cardiac vasculature https://github.com/path-openings/RORPO/tree/develop. This particular branch was the one that was successfully compiled and therefore used to run visualization experiments.

---

**Figure S1:**
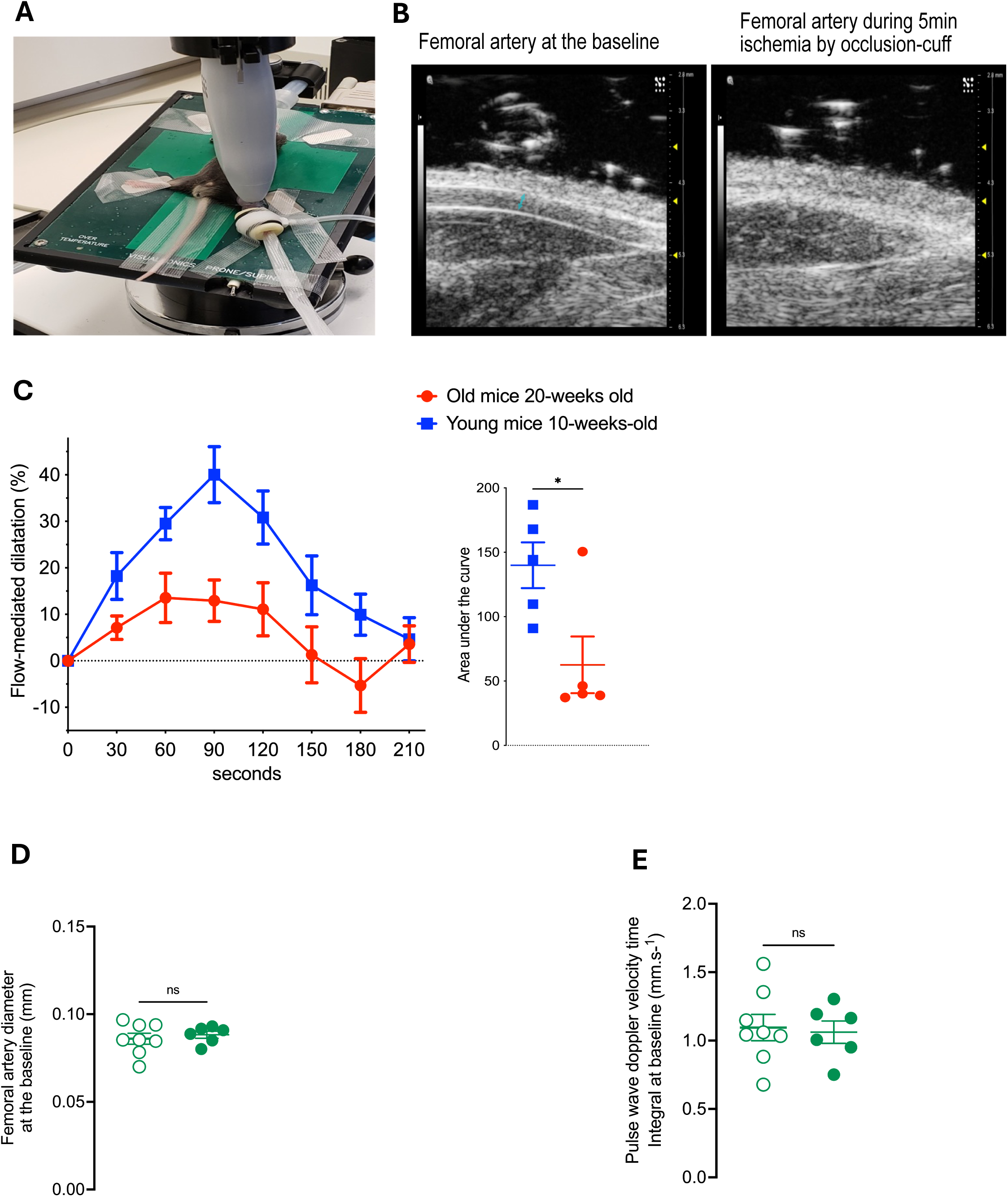
Endothelial function measurement by flow-mediated dilation. **A.** Picture depicting the most stable placement that allowed reliable and repetitive assessment of flow-mediated dilatation (FMD) measured in the femoral artery by ultrasound. Position of a mouse taped in supine position on a controlled heating pad is seen as well as the ultrasound probe and the position of the Occlusion cuff from the plethysmograph CODA monitor placed around the mouse lower limb. **B.** Representative ultrasound pictures of femoral artery diameter outlined by the blue arrow at the baseline and during ischemia. **C.** FMD measured in mice femoral artery during reactive hyperemia over 4min in response to 5 min ischemia in 10-week-old C57Bl6 mice n=5 and 20-week-old C57Bl6 mice n=5. Values are expressed as mean±SEM. Significance was calculated for the FMD with a 2-way ANOVA followed by Tukey’s multiple comparison test (Time effect: *\*\*\*P<0.001*; Age effect: *\*P<0.05;* Time x Age effect: *\*P<0.05*). Significance was calculated for the area under the curve with an unpaired Student *t* test *\*P<0.05*. **D.** Mean femoral artery diameter at the baseline and **E**. Pulse wave doppler velocity time integral quantified by ultrasound analysis Vevo Software over platform over measurements per mice in 2-month-old *Atg16l1^wt^* mice n=8 and *Atg16l1^!1WD^ ^ki^* mice n=6. Significance was calculated for the diameter and the pulse wave velocity with an unpaired Student *t* test.

**Figure S2:**
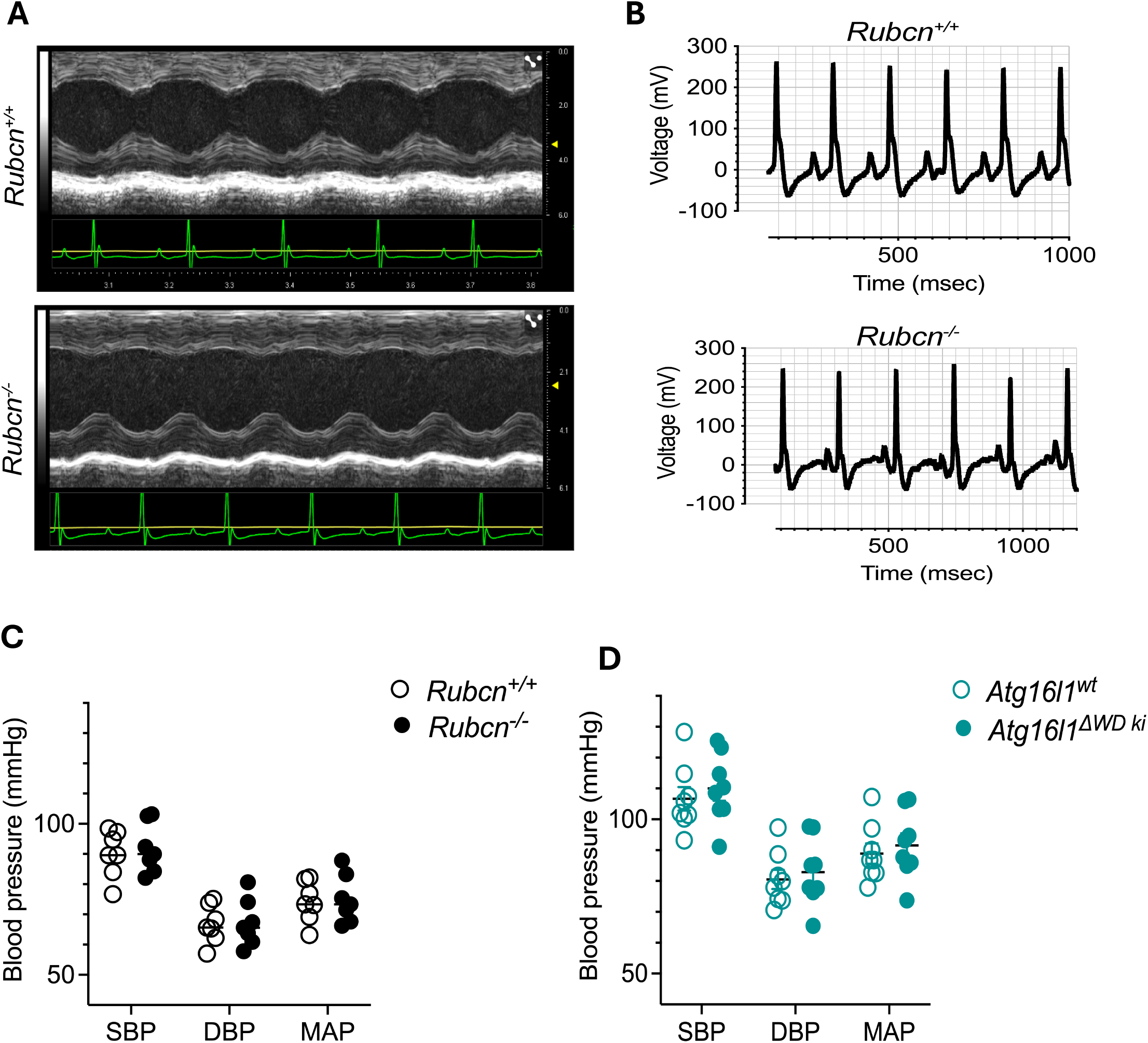
Cardiac function, ECG and blood pressure measurement. **A.** Representative ultrasound pictures of electrocardiogram in *Rubcn^+/+^* and *Rubcn^-/-^* mice**. B.** Representative electrocardiography (ECG) waveforms in *Rubcn^+/+^* and *Rubcn^-/-^* mice as a function of Voltage over time. **C** and **D** Systolic blood pressure (SBP), Diastolic blood pressure (DBP) and Mean Arterial blood pressure (MAP) in (**C**) 2-month-old *Rubcn^+/+^* mice n=7 and *Rubcn^-/-^* mice n=7 and (**D**) *Atg16l1^wt^* mice n=8 and *Atg16l1^!1WD^ ^ki^* mice n=8. Values are expressed as mean±SEM. Significance was calculated for the SBP, DBP and MAP with a 2-way ANOVA followed by sidak’s multiple comparison test.

**Figure S3:**
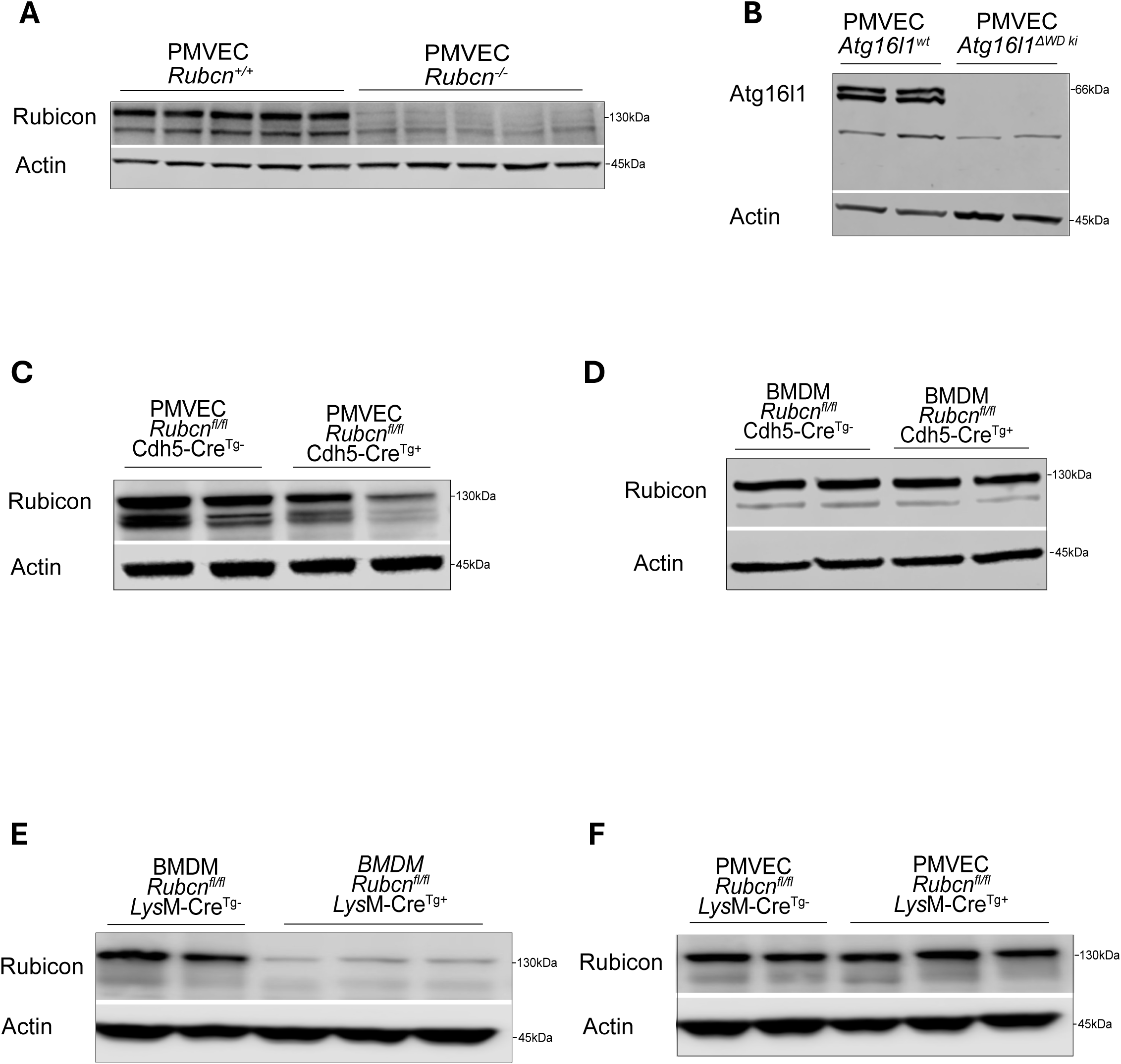
Confirmation mice model genotype by western blot A, C-F. Representative immunoblots of Rubicon and Actin in whole cell lysate from 2-month-old (**A**) *Rubcn^+/+^* and *Rubcn^-/-^* mice EC lung ; (**C**) *Rubcn^fl/fl^, Cdh5-Cre^Tg+^* and *Rubcn^fl/fl^, Cdh5-Cre^Tg-^* mice EC lung ; (**D**) *Rubcn^fl/fl^, Cdh5-Cre^Tg+^* and *Rubcn^fl/fl^, Cdh5-Cre^Tg-^* mice BMDM ; (**E**) *Rubcn^fl/fl^, LysM-Cre^Tg+^* and *Rubcn^fl/fl^, LysM-Cre^Tg-^* mice BMDM; (**F**) *Rubcn^fl/fl^, LysM-Cre^Tg+^* n=9 and *Rubcn^fl/fl^, LysM-Cre^Tg-^*mice EC lung; (**B**) Representative immunoblots of Atg16l1 and Actin in whole cell lysate from 2 month-old *Atg16l1^wt^* mice and *Atg16l1^!1WD^ ^ki^*

**Figure S4:**
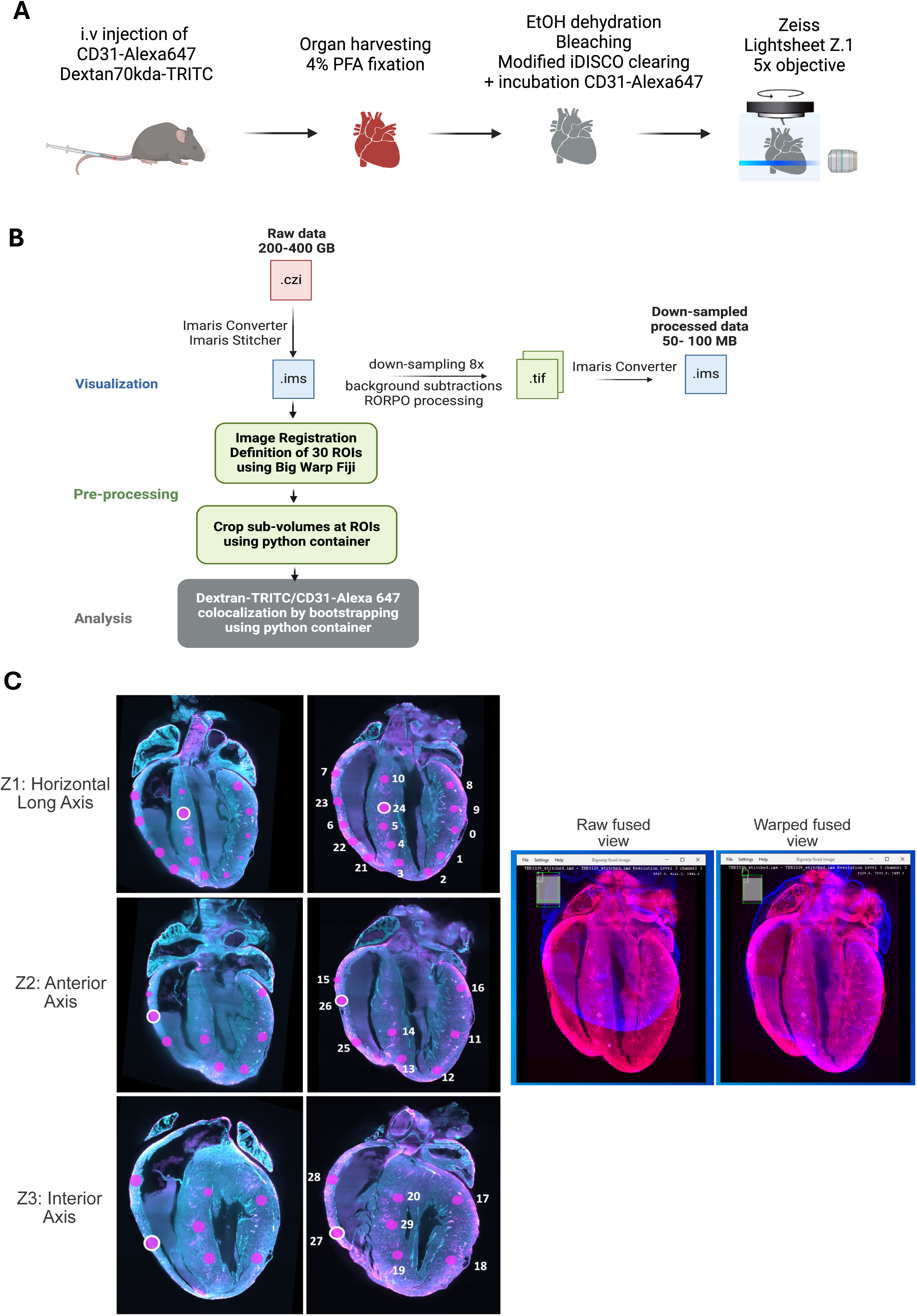
Light sheet microscopy imaging workflow and analysis. **A, B.** Workflow for light sheet microscopy imaging depicting (**A**) cardiac tissue clearing and immunolabelling (**B**) data visualization, pre-processing and image analysis. (**C**) Overlay pictures with CD31-Alexa647 (Turquoise) and Dextran70kDa-TRITC (Magenta) depicting on the left: image registration step using Big Wrap from Fiji software with representative location of 30 regions of Interest (ROI) placed over 3 different Z-steps of a cardiac volume ; on the right: and the raw fused view image of the heart volume at the anterior axis Z-step compared to warped fused aligned view.

**Figure S5:**
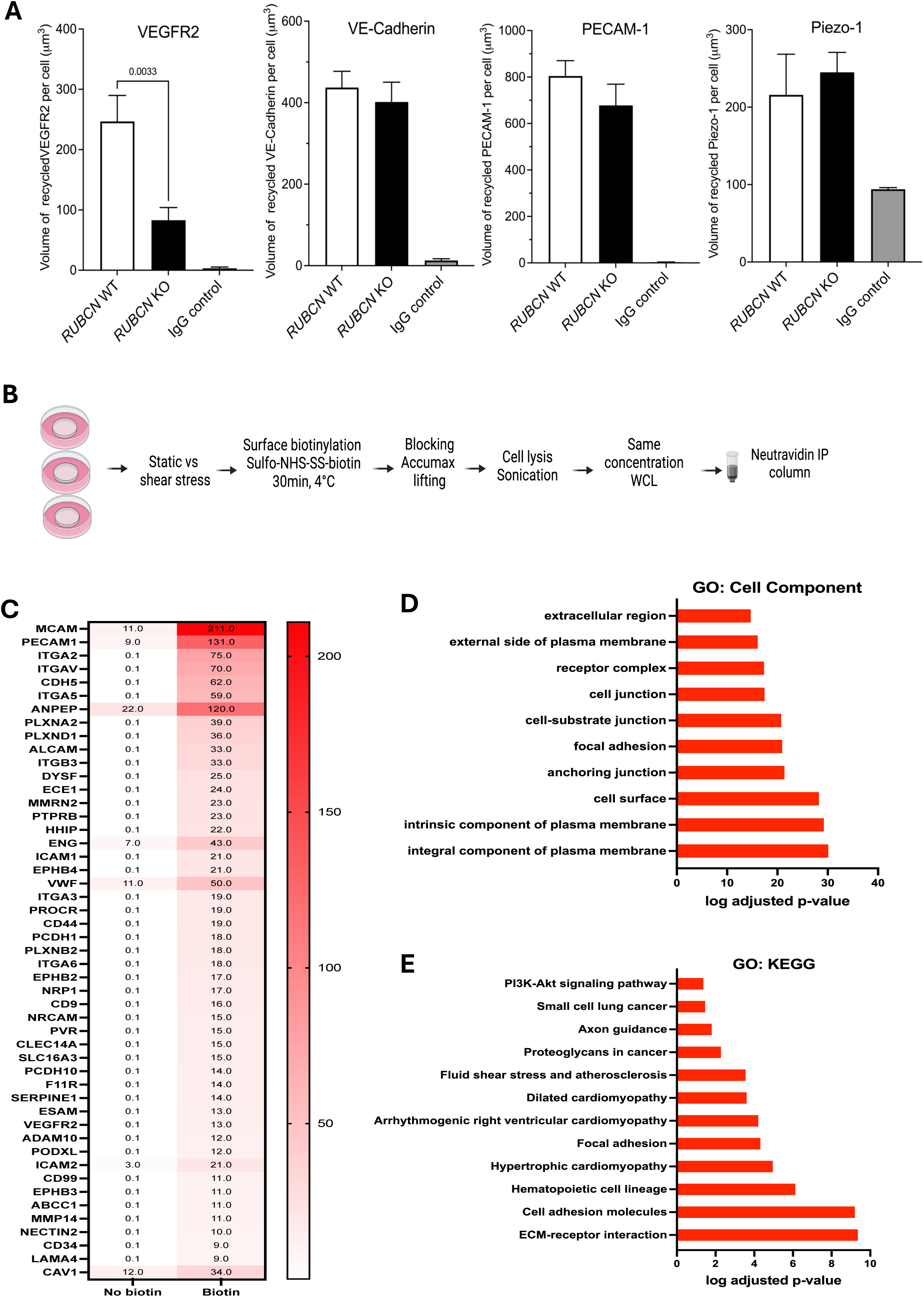
Workflow for surface proteomic assay and technique specificity. **A.** Volume of recycled VEGFR2, CD31, VE-cadherin, Piezo-1 per cell with each point representing the average volume for at least 3 cells per field view from HAEC *RUBCN* KO and *RUBCN* WT (n=3 independent replicates per group) exposed to 5 dynes of laminar flow during 24h. IgG controls as a control for antibodies. Values are expressed as mean±SEM. Significance was calculated for the volume of VEGFR2 recycled per cell with an unpaired Student *t* test. **B.** Workflow for surface proteomic assay. **C.** Heatmap displaying the top 49 proteins differentially expressed in HAEC when exposed to Sulfo-NHS-SS-biotin compared to no exposure. Numbers are representing the relative abundance of proteins assessed by proteomic. **D** and **E**. Histogram exhibiting pathway enrichment analysis of the proteins differentially expressed in HAEC when exposed to Sulfo-NHS-SS-biotin compared to no exposure showing (**D**) the top 10 Gene Ontology (GO) term Cellular_component and (**E**) Kyoto Encyclopedia of Genes and Genomes (KEGG) pathway.

**Figure S6:**
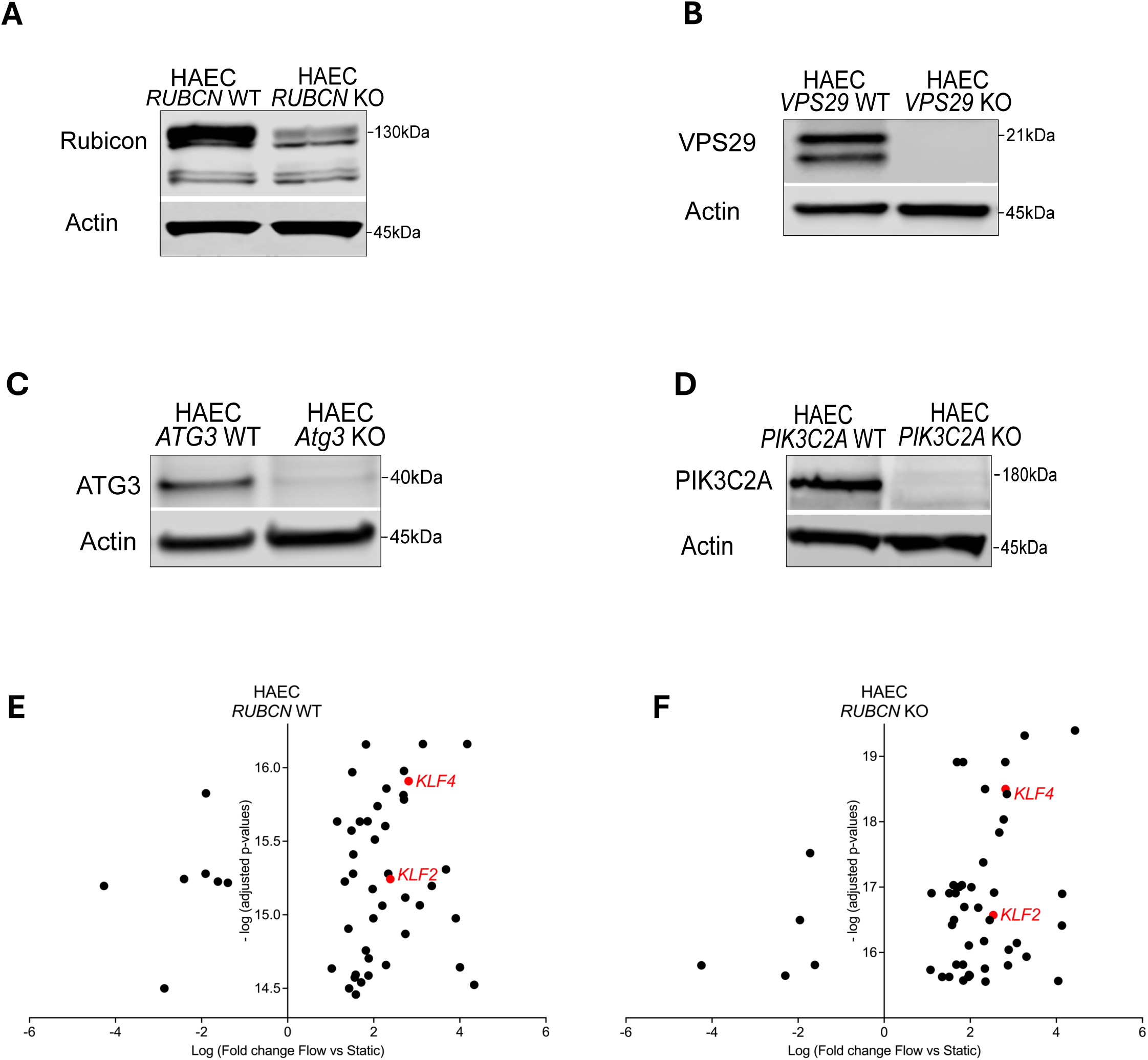
Confirmation of *RUBCN*, *VPS29, ATG3* and *PIK3C2A* knock-out in HAEC A-D. Representative immunoblots of Rubicon, VPS29, Atg3, PIK3C2A and Actin in whole cell lysate when *RUBCN*, *VPS29*, *ATG3* and *PIK3C2A* were knocked out in HAEC. **E**. Volcano plots displaying the top 50 mRNA differentially expressed in HAEC *Rubcn* WT when exposed to 5 dynes of laminar flow during 24h vs static conditions (n=4 independent replicates per group). **F**. Volcano plots displaying the top 50 mRNA differentially expressed in HAEC *RUBCN* KO or HAEC *RUBCN* WT when exposed to 5 dynes of laminar flow during 24h vs static conditions (n=4 independent replicates per group).

**Figure S7:**
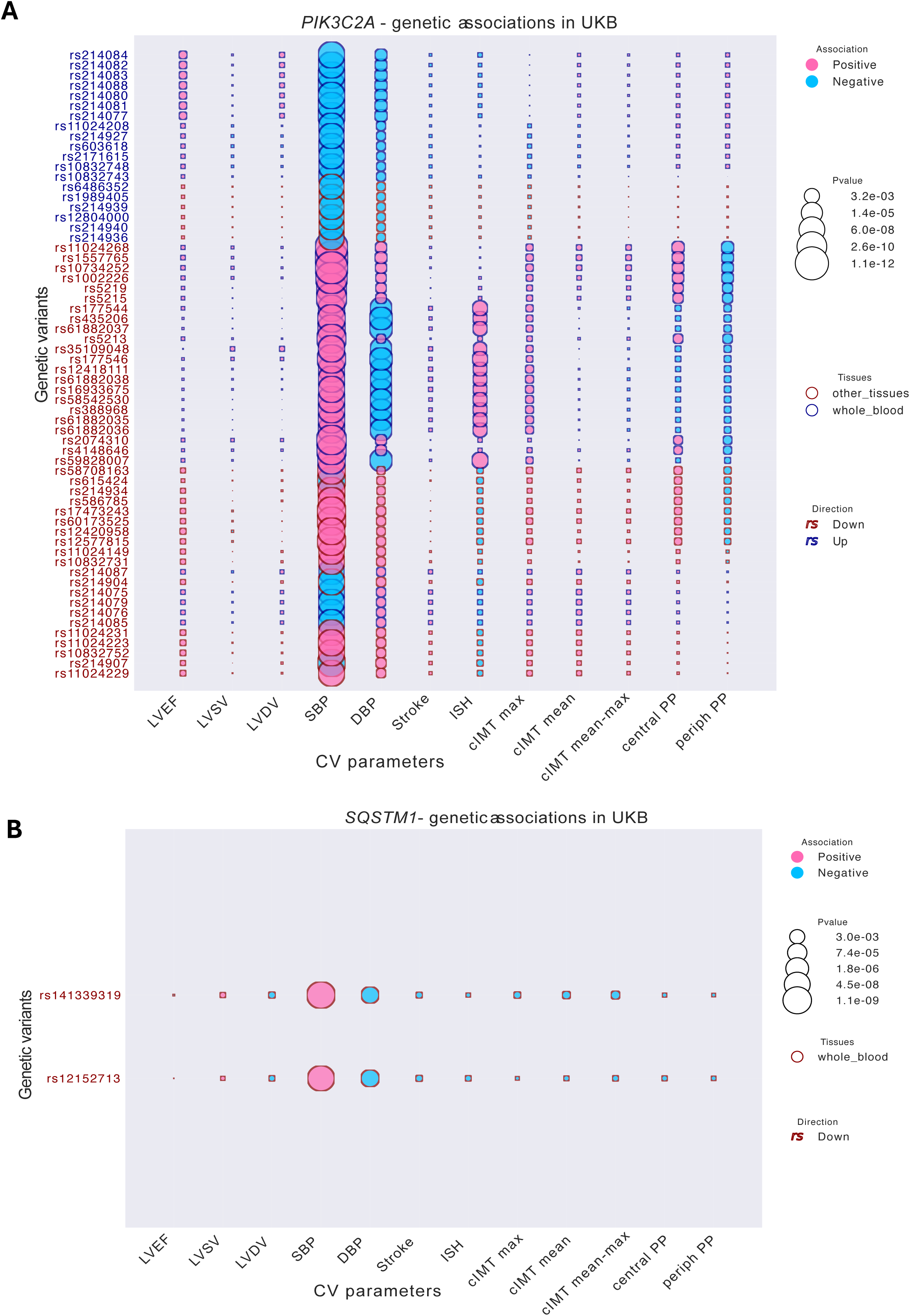
*PIK3C2A* and *SQSTM1* genetic variants in UK biobank. Bubble plot exhibiting a single SNPs analysis in (**A**) *PIK3C2A and* (**B**) *SQSTM1* loci associated with 12 cardiovascular parameters in the UK biobank. The dark blue color for the SNPs font indicates that the SNPs in genetic variants loci were significantly associated with an upregulation of the mRNA in the whole blood. The bubble size indicates the P-value for the association and the color bar indicates the direction of the association (positive in pink or negative in blue).

**Figure S8:**
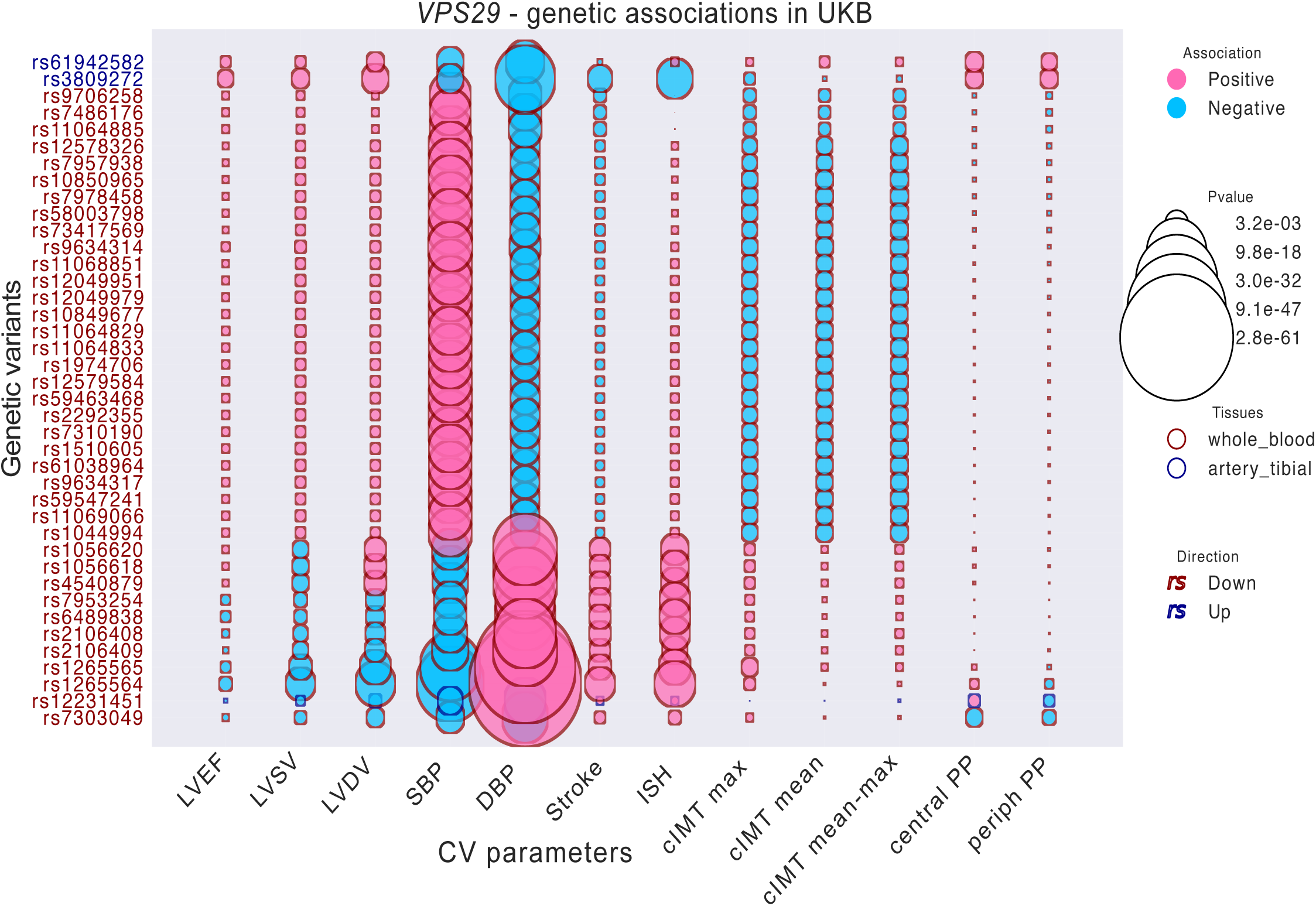
*VPS29*-genetic variants associations in UK biobank. Bubble plot exhibiting a single SNPs analysis in *VPS29* loci associated with 12 cardiovascular parameters in the UK biobank. The dark blue color for the SNPs font indicates that the SNPs in genetic variants loci were significantly associated with an upregulation of the mRNA in the whole blood. The bubble size indicates the P-value for the association and the color bar indicates the direction of the association (positive in pink or negative in blue).

**Figure S9:**
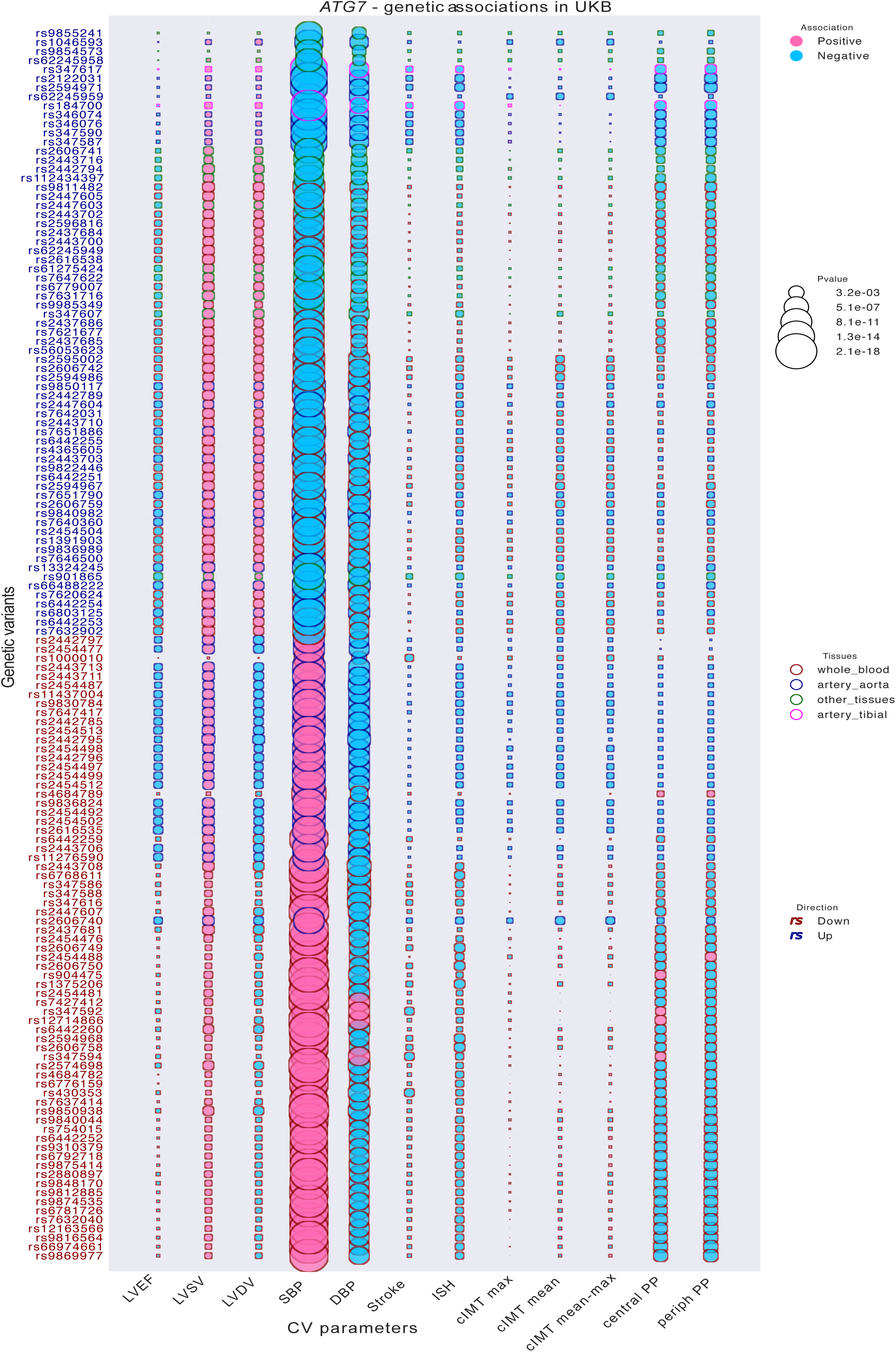
*ATG7*-genetic variants associations in UK biobank. Bubble plot exhibiting a single SNPs analysis in *ATG7* loci associated with 12 cardiovascular parameters in the UK biobank. The dark blue color for the SNPs font indicates that the SNPs in genetic variants loci were significantly associated with an upregulation of the mRNA in the whole blood. The bubble size indicates the P-value for the association and the color bar indicates the direction of the association (positive in pink or negative in blue).

**Figure S10:**
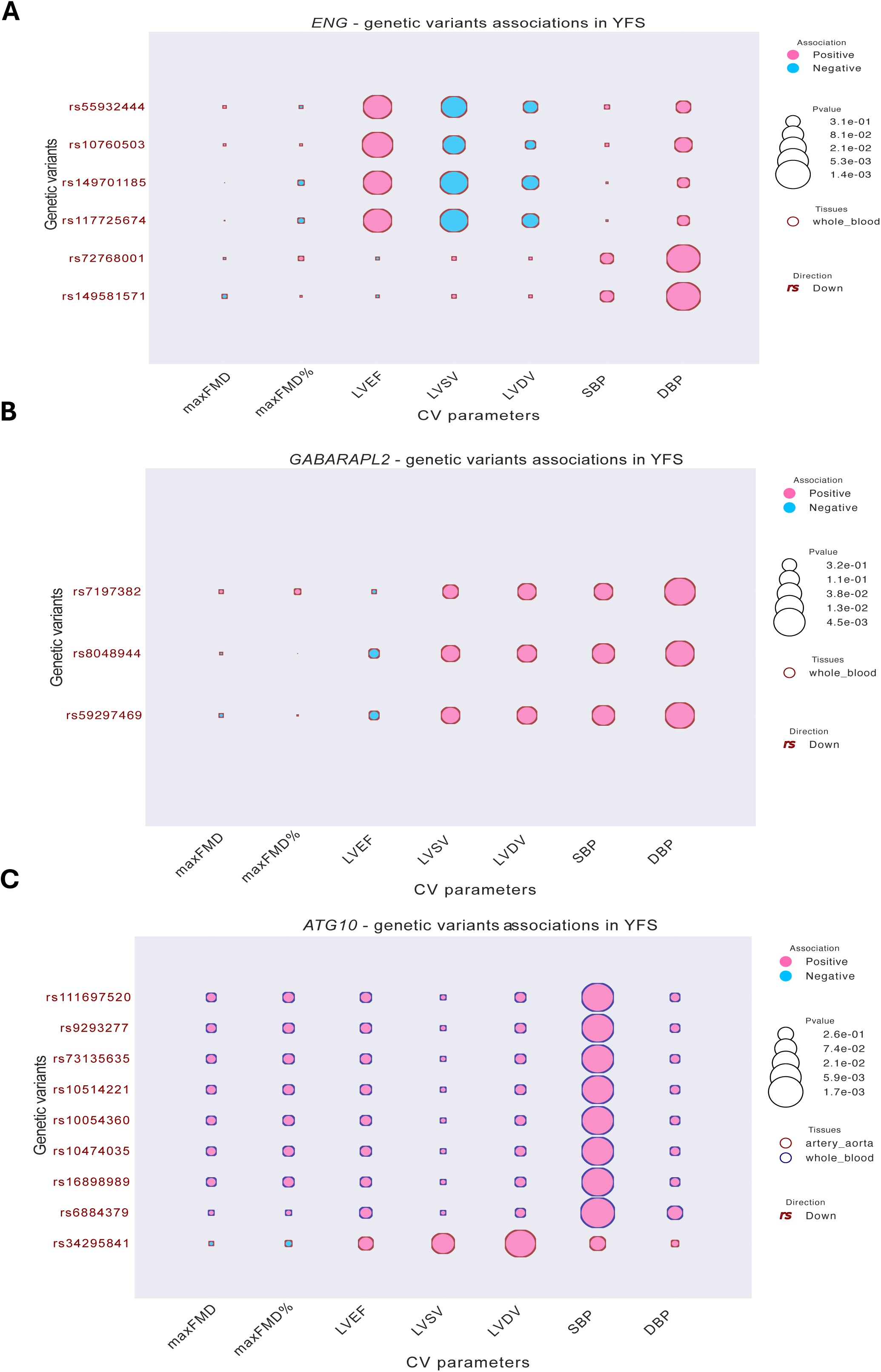
*ENG*, *GABARAPL2*, *ATG10* genetic variants associations in YFS. Bubble plot exhibiting genetic variants analysis in (**A**) *ENG*, (**B**) *GABARAPL2*, (**C**) *ATG10* loci associated with 7 cardiovascular parameters in the Young Finns Study. The dark blue color for the SNPs font indicates that the genetic variants were significantly associated with an upregulation of the mRNA in the whole blood. The bubble size indicates the P-value for the association and the color bar indicates the direction of the association (positive in pink or negative in blue).

**Figure S11:**
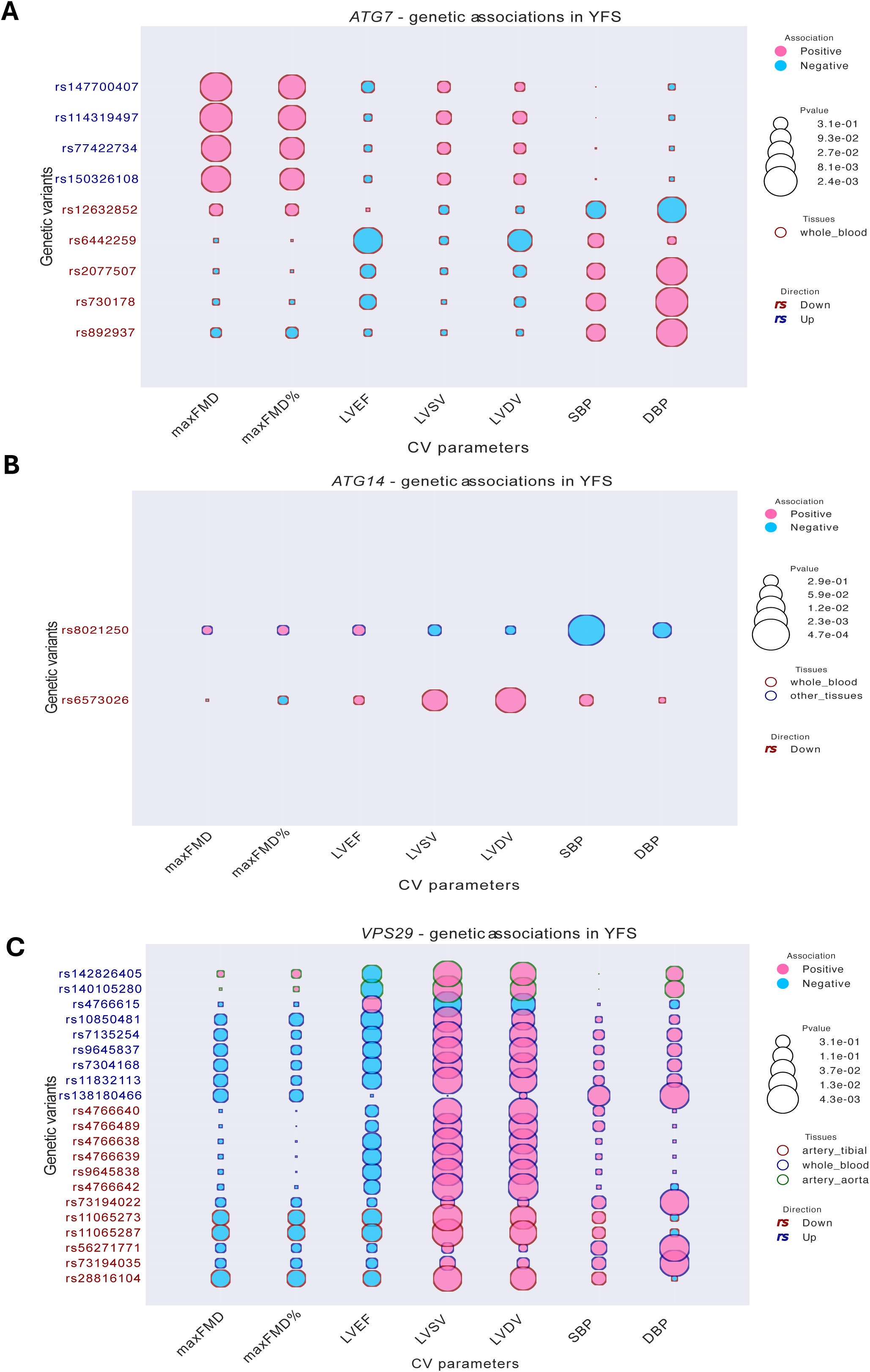
*ATG7*, *ATG14*, *VPS29* genetic variants associations in YFS. Bubble plot exhibiting genetic variants analysis in (**A**) *ATG7*, (**B**) *ATG14*, (**C**) *VPS29* loci associated with 7 cardiovascular parameters in the Young Finns Study. The dark blue color for the SNPs font indicates that the genetic variants were significantly associated with an upregulation of the mRNA in the whole blood. The bubble size indicates the P-value for the association and the color bar indicates the direction of the association (positive in pink or negative in blue).

**Figure S12:**
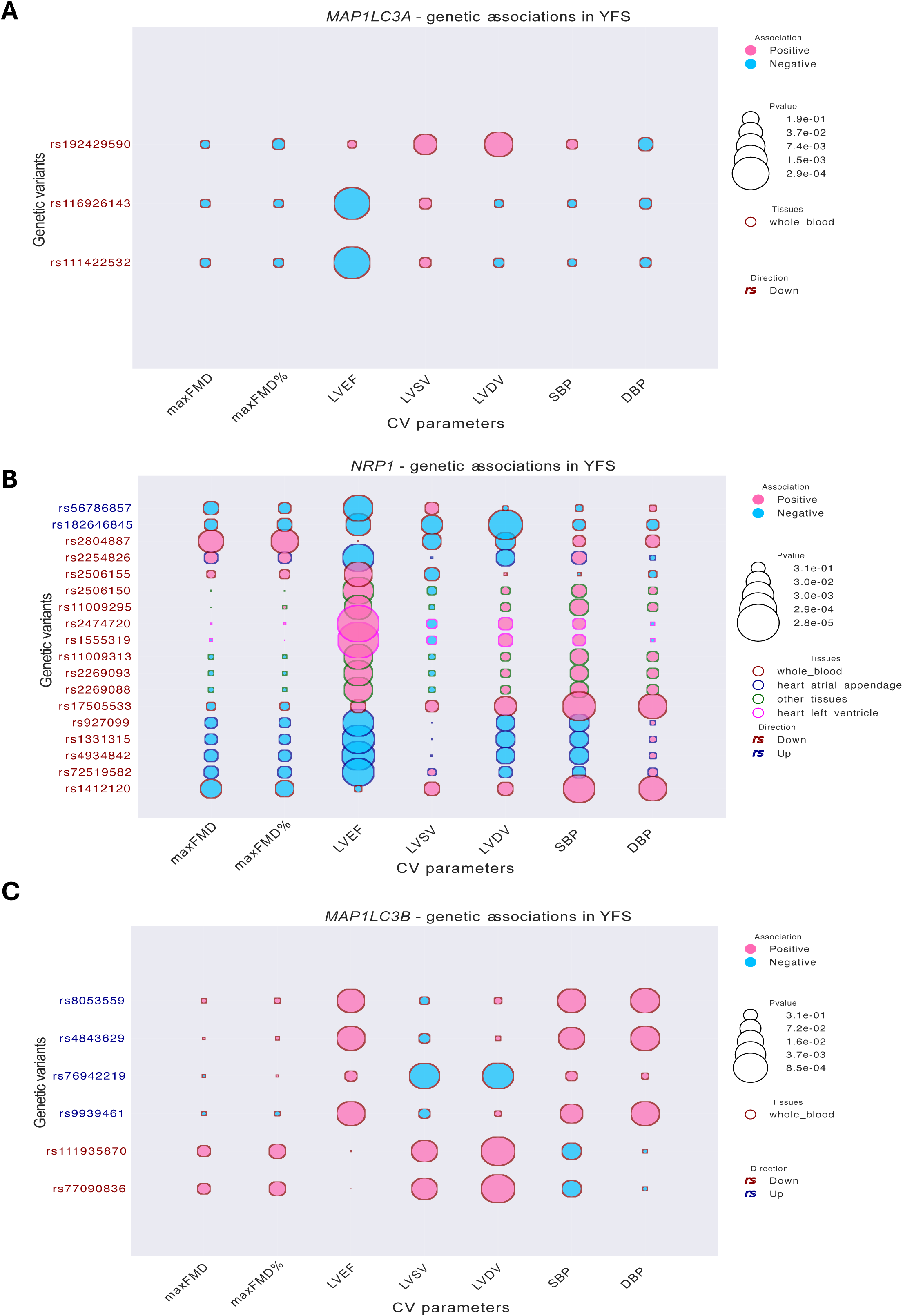
*MAP1LC3A*, *NRP1*, *MAP1LC3B* genetic variants associations in YFS. Bubble plot exhibiting genetic variants analysis in (**A**) *MAP1LC3A*, (**B**) *NRP1*, (**C**) *MAP1LC3B* loci associated with 7 cardiovascular parameters in the Young Finns Study. The dark blue color for the SNPs font indicates that the genetic variants were significantly associated with an upregulation of the mRNA in the whole blood. The bubble size indicates the P-value for the association and the color bar indicates the direction of the association (positive in pink or negative in blue).

**Figure S13:**
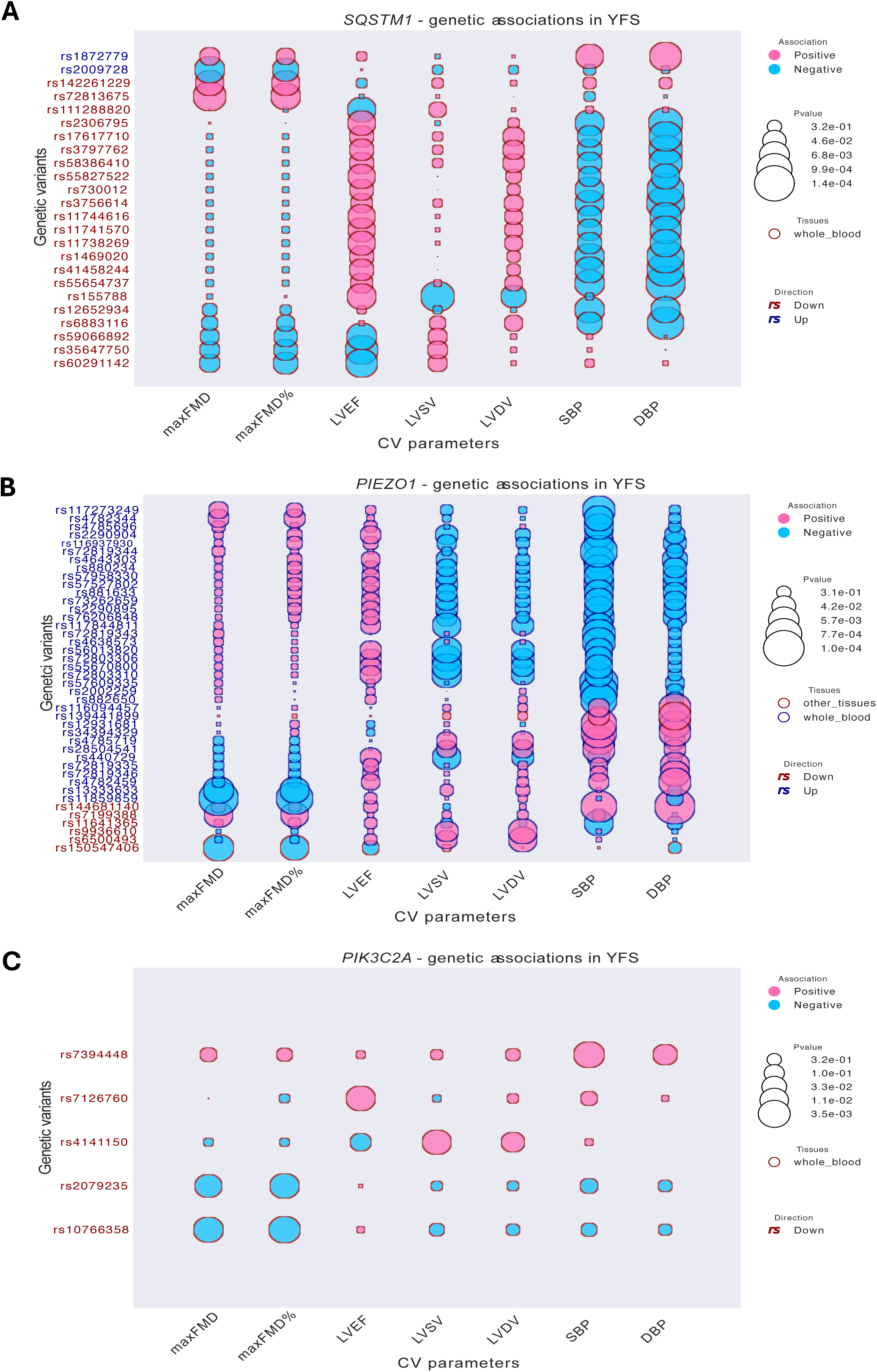
*SQSTM1*, *PIEZO1*, *PIK3C2A* genetic variants associations in YFS. Bubble plot exhibiting genetic variants analysis in (**A**) *SQSTM1*, (**B**) *PIEZO1*, (**C**) *PIK3C2A* loci associated with 7 cardiovascular parameters in the Young Finns Study. The dark blue color for the SNPs font indicates that the genetic variants were significantly associated with an upregulation of the mRNA in the whole blood. The bubble size indicates the P-value for the association and the color bar indicates the direction of the association (positive in pink or negative in blue).

**Figure S14:**
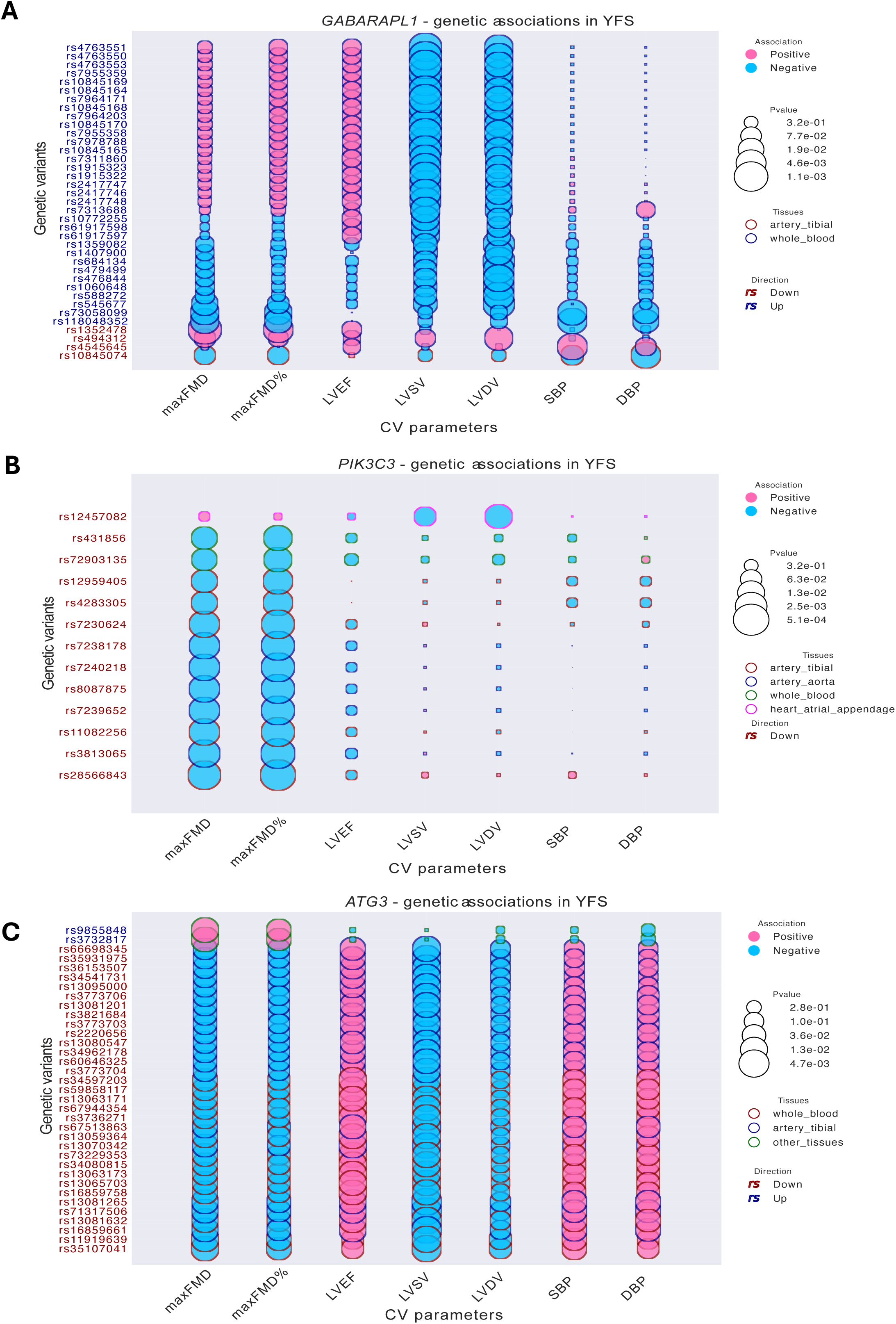
*GABARAPL1*, *PIK3C3*, *ATG3* genetic variants associations in YFS. Bubble plot exhibiting genetic variants analysis in (**A**) *GABARAPL1*, (**B**) *PIK3C3*, (**C**) *ATG3* loci associated with 7 cardiovascular parameters in the Young Finns Study. The dark blue color for the SNPs font indicates that the genetic variants were significantly associated with an upregulation of the mRNA in the whole blood. The bubble size indicates the P-value for the association and the color bar indicates the direction of the association (positive in pink or negative in blue).

**Figure S15:**
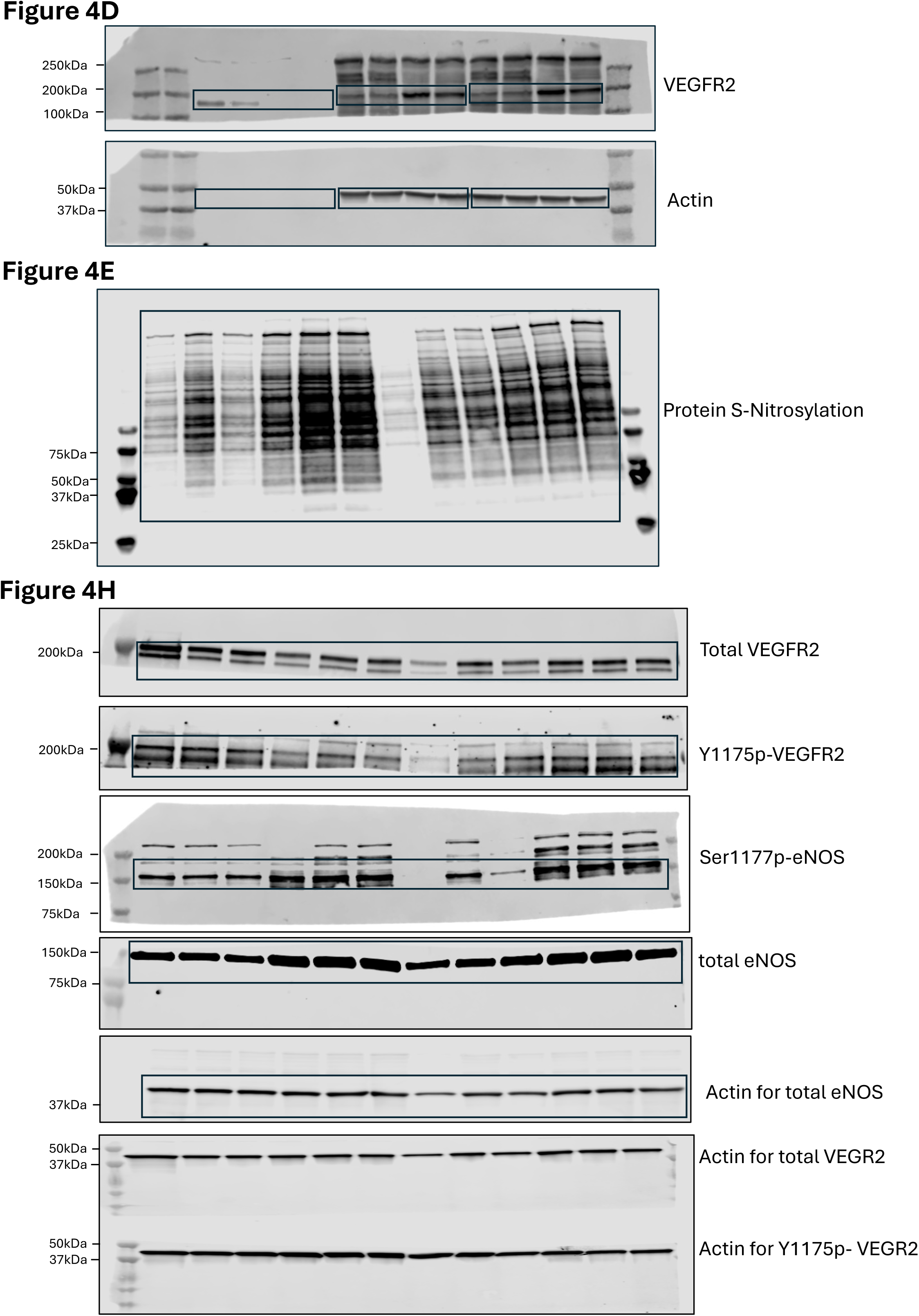
Unprocessed western blots for. **Figure 4** Uncropped blots from which figure 4 in this paper were derived from. Black boxes mark where western blots have been cropped. Molecular weights are indicated for each western blot.

**Figure S16:**
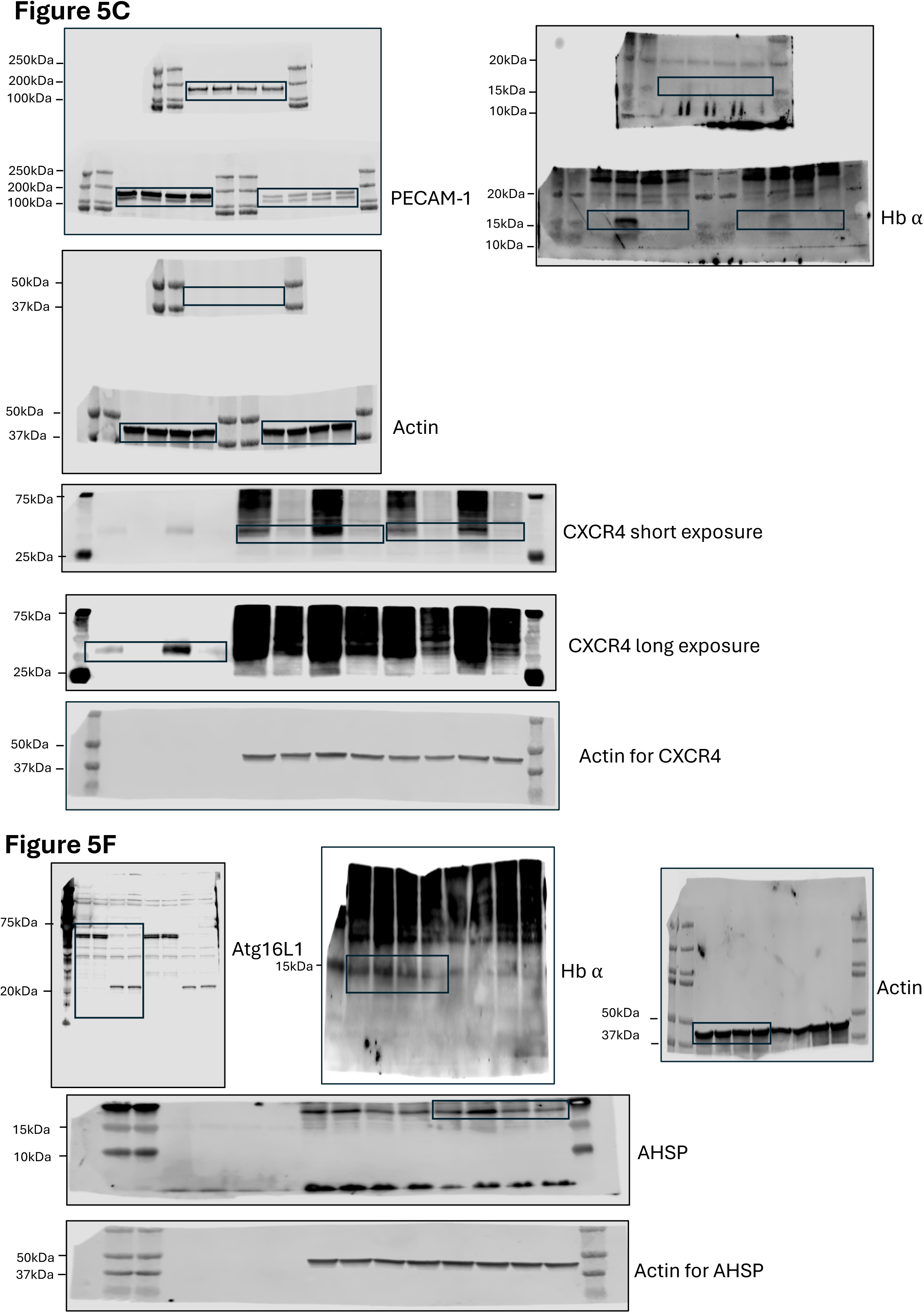
Unprocessed western blots for. **Figure 5** Uncropped blots from which figure 5 in this paper were derived from. Black boxes mark where western blots have been cropped. Molecular weights are indicated for each western blot.

**Figure S17:**
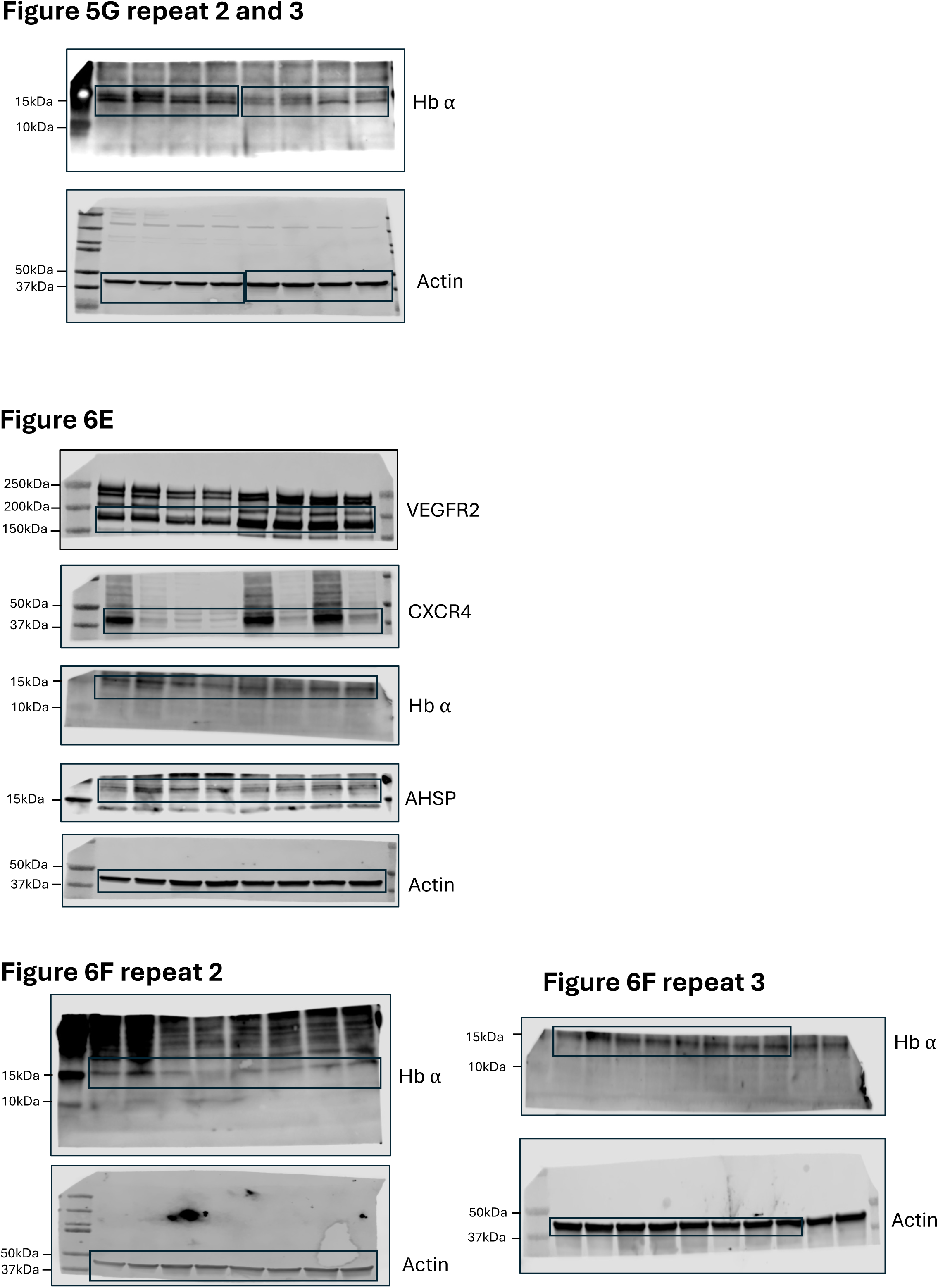
Unprocessed western blots for. **Figure 6** Uncropped blots from which figure 6 in this paper were derived from. Black boxes mark where western blots have been cropped. Molecular weights are indicated for each western blot.

**Figure S18:**
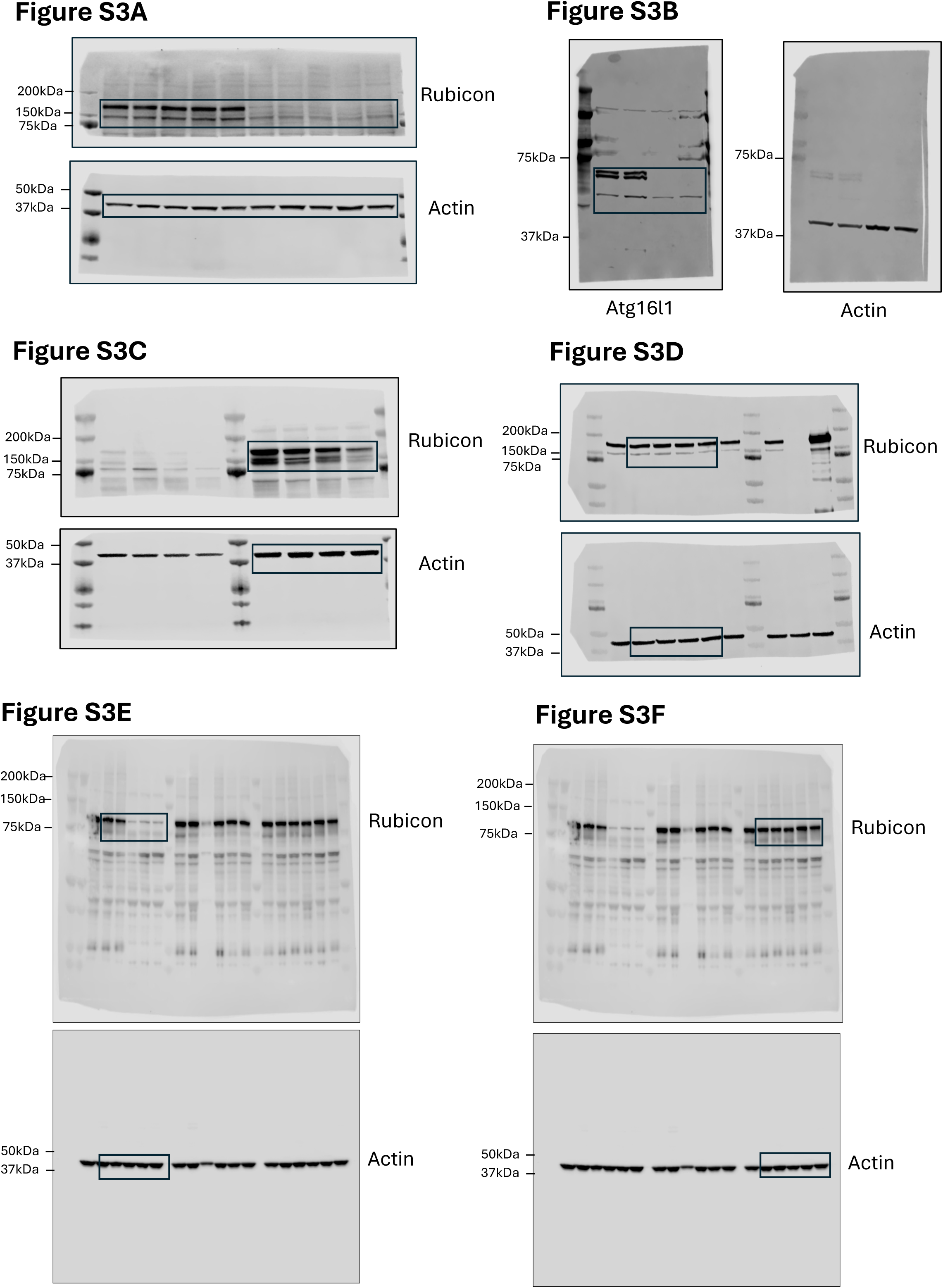
Unprocessed western blots for Figure S3. Uncropped blots from which figure S3 in this paper were derived from. Black boxes mark where western blots have been cropped. Molecular weights are indicated for each western blot.

**Figure S19:**
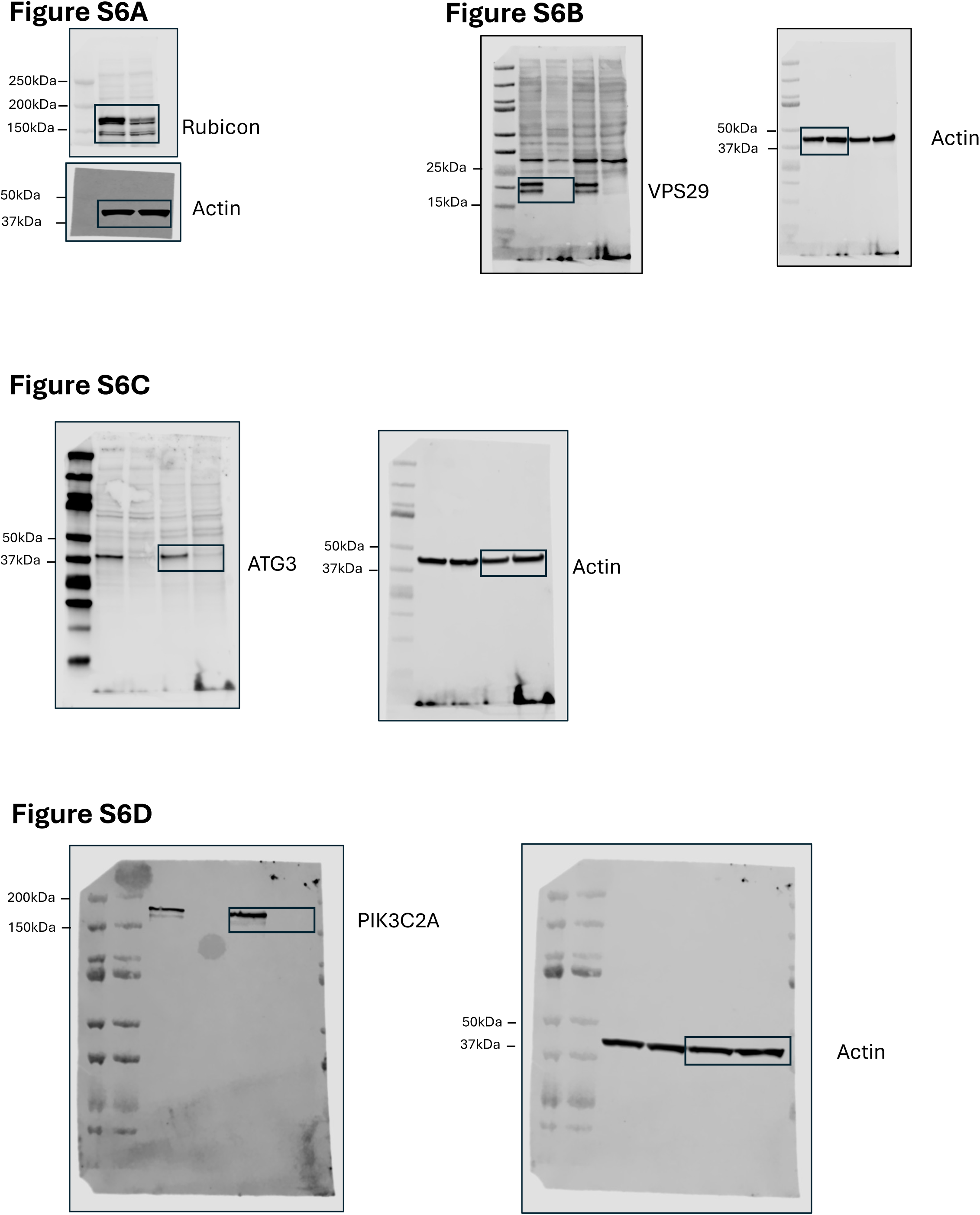
Unprocessed western blots for Figure S6. Uncropped blots from which figure S6 in this paper were derived from. Black boxes mark where western blots have been cropped. Molecular weights are indicated for each western blot.

**Table S1:** CV parameters for Atg16l1*^ΔWD^ ^ki^* phenotype in males.

|  | 2-month-old |  | 6-month-old |  |
| --- | --- | --- | --- | --- |
|  | *Atg16l1*<sup>wt</sup><br>n=8 | *Atg16l1*<sup>ΔWD *ki*</sup><br>n=6 | *Atg16l1*<sup>wt</sup><br>n=8 | *Atg16l1*<sup>ΔWD *ki*</sup><br>n=6 |
| Heart Rate (BPM) | 384.1±4.6 | 384.1±3.3 | 386.3±3.3 | 368.3±9.7* |
| LV diameter,s (mm) | 1.8±0.1 | 2.4±0.2* | 2.5±0.2<sup>††</sup> | 2.3±0.2 |
| LV diameter,d (mm) | 3.2±0.1 | 3.5±0.1* | 4.0±0.1<sup>††††</sup> | 3.9±0.1 |
| LV volume,s (μl) | 10.7±1.6 | 21.8±3.1* | 23.1±3.5<sup>††</sup> | 19.5±3.7 |
| LV volume,d (μl) | 40.3±3.5 | 52.5±3.3* | 69.2±4.4<sup>††††</sup> | 67.3±3.6<sup>‡</sup> |
| Stroke volume (μl) | 29.6±2.1 | 30.6±1.3 | 46.1±2.2<sup>††††</sup> | 47.8±1.4<sup>††††</sup> |
| LVEF (%) | 74.2±2.2 | 59.4±3.9** | 67.6±3.2 | 72.1±4.3<sup>‡</sup> |
| LVFS (%) | 42.1±1.8 | 31.2±2.8** | 37.6±2.5 | 41.6±3.9<sup>‡</sup> |
| Cardiac Output (mL/min) | 11.4±0.9 | 11.8±0.5 | 17.8±0.8<sup>††††</sup> | 17.6±0.6<sup>††††</sup> |
| LV Mass (mg) | 94.8±4.4 | 115.4±7.8* | 151.0±5.3<sup>††††</sup> | 148.6±3.9<sup>‡</sup> |
| LV Mass Corrected (mg) | 75.9±3.5 | 92.3±6.2* | 120.8±4.2<sup>†††</sup> | 118.9±3.1<sup>††</sup> |
| LVAW, s (mm) | 1.2±0.0 | 1.2±0.0 | 1.6±0.1 | 1.7±0.1 |
| LVAW,d (mm) | 1.0±0.1 | 1.0±0.0 | 1.1±0.0 | 1.1±0.0 |
| LVPW, s (mm) | 1.3±0.0 | 1.1±0.1 | 1.3±0.0 | 1.2±0.0 |
| LVPW,d (mm) | 0.8±0.0 | 0.8±0.0 | 0.8±0.0 | 0.8±0.0 |
| Body weight (g) | 26.0±0.8 | 25.3±0.8 | 29.8±1.1<sup>††††</sup> | 30.5±0.9<sup>††††</sup> |

Results are presented as mean ± SEM. Significance was calculated for the CV parameters with a 2-way ANOVA followed by Tukey's multiple comparison test. \*\* $P < 0.01$ , \* $P < 0.05$ for comparison between *Atg16l1* <sup>$\Delta$ WD *ki*</sup> vs *Atg16l1*<sup>wt</sup> at the same time-period. <sup>††††</sup> $P < 0.0001$ , <sup>†††</sup> $P < 0.001$ , <sup>††</sup> $P < 0.01$ for comparison between a parameter assessed at 2-month-old versus 6-month-old for *Atg16l1*<sup>wt</sup>. <sup>††††</sup> $P < 0.0001$ , <sup>†††</sup> $P < 0.001$ , <sup>††</sup> $P < 0.01$ , <sup>‡</sup> $P < 0.05$ for comparison between cardiac parameter assessed at 2-month-old versus 6-month-old for *Atg16l1* <sup>$\Delta$ WD *ki*</sup>. LV, left ventricle; s, systole; d, diastole; LVEF, left ventricle ejection fraction; LVFS, left ventricle fractional shortening; LVAWs, left ventricular end-systolic anterior wall diameter; LVAWd, left ventricular end-diastolic anterior wall diameter; LVPWs, left ventricular end-systolic posterior wall diameter; LVAWd, left ventricular end-diastolic posterior wall diameter.

**Table S2:** CV parameters for *Rubcn ^-/-^* phenotype in males.

|  | 2-month-old |  | 6-month-old |  |
| --- | --- | --- | --- | --- |
|  | *Rubcn*<sup>+/+</sup><br>n=8 | *Rubcn*<sup>-/-</sup><br>n=9 | *Rubcn*<sup>+/+</sup><br>n=8 | *Rubcn*<sup>-/-</sup><br>n=9 |
| Heart Rate (BPM) | 374.6±8.1 | 404.0±6.4<sup>**</sup> | 371.0±5.1 | 372.3±9.4<sup>‡</sup> |
| LV diameter,s (mm) | 2.3±0.1 | 2.9±0.1<sup>***</sup> | 2.2±0.1 | 2.8±0.1<sup>***</sup> |
| LV diameter,d (mm) | 3.6±0.1 | 3.9±0.1 | 3.6±0.1 | 4.2±0.1<sup>**‡</sup> |
| LV volume,s (μl) | 18.1±2.3 | 33.3±2.5<sup>***</sup> | 16.9±1.8 | 30.9±3.4<sup>***</sup> |
| LV volume,d (μl) | 56.2±4.3 | 68.1±3.7 | 56.2±4.5 | 78.5±5.7<sup>**‡</sup> |
| Stroke volume (μl) | 38.1±2.6 | 34.8±2.1 | 39.4±3.3 | 47.6±2.8<sup>*††</sup> |
| LVEF (%) | 68.4±2.4 | 51.3±1.9<sup>****</sup> | 70.4±2.1 | 61.5±2.4<sup>**‡</sup> |
| LVFS (%) | 37.7±1.9 | 25.8±1.1<sup>****</sup> | 39.3±1.7 | 33.0±1.8<sup>*‡</sup> |
| Cardiac Output (mL/min) | 14.2±0.9 | 14.0±0.8 | 14.7±1.3 | 17.7±1.1<sup>*‡</sup> |
| LV Mass (mg) | 123.1±5.4 | 129.2±7.7 | 172.6±9.1<sup>††</sup> | 160.0±7.9<sup>‡</sup> |
| LV Mass Corrected (mg) | 98.5±4.3 | 103.3±6.2 | 138.1±7.3<sup>††</sup> | 128.0±6.4<sup>‡</sup> |
| LVAW, s (mm) | 1.7±0.1 | 1.3±0.1<sup>*</sup> | 2.1±0.1<sup>††</sup> | 1.7±0.1<sup>**‡</sup> |
| LVAW,d (mm) | 1.1±0.0 | 0.9±0.1 | 1.5±0.1<sup>†††</sup> | 1.1±0.1<sup>***</sup> |
| LVPW, s (mm) | 1.2±0.0 | 1.0±0.1 | 1.3±0.0 | 1.1±0.1 |
| LVPW,d (mm) | 0.8±0.0 | 0.8±0.1 | 0.9±0.1 | 0.8±0.0 |
| Body weight (g) | 24.4±0.6 | 24.6±0.7 | 33.7±0.6<sup>†††</sup> | 31.0±0.9<sup>†††</sup> |

Results are presented as mean ± SEM. Significance was calculated for the CV parameters with a 2-way ANOVA followed by Tukey's multiple comparison test. \*\*\*\* $P < 0.0001$ , \*\*\* $P < 0.001$ , \*\* $P < 0.01$ , \* $P < 0.05$ for comparison between *Rubcn*<sup>-/-</sup> vs *Rubcn*<sup>+/+</sup> at the same time-period. ††† $P < 0.0001$ , †† $P < 0.001$ , † $P < 0.01$ for comparison between a parameter assessed at 2-month-old versus 6-month-old for *Rubcn*<sup>+/+</sup>. ††† $P < 0.0001$ , †† $P < 0.001$ , † $P < 0.01$ , ‡ $P < 0.05$ for comparison between cardiac parameter assessed at 2-month-old versus 6-month-old for *Rubcn*<sup>-/-</sup>. LV, left ventricle; s, systole; d, diastole; LVEF, left ventricle ejection fraction; LVFS, left ventricle fractional shortening; LVAWs, left ventricular end-systolic anterior wall diameter; LVAWd, left ventricular end-diastolic anterior wall diameter; LVPWs, left ventricular end-systolic posterior wall diameter; LVAWd, left ventricular end-diastolic posterior wall diameter.

**Table S3:** CV parameters for *Rubcn ^-/-^* phenotype in females.

|  | 2-month-old |  | 6-month-old |  |
| --- | --- | --- | --- | --- |
|  | *Rubcn*<sup>+/+</sup><br>n=6 | *Rubcn*<sup>-/-</sup><br>n=8 | *Rubcn*<sup>+/+</sup><br>n=6 | *Rubcn*<sup>-/-</sup><br>n=8 |
| Heart Rate (BPM) | 370.7±21.2 | 337.2±17.9 | 378.5±3.8 | 356.2±13.8 |
| LV diameter,s (mm) | 2.3±0.1 | 2.5±0.2 | 2.3±0.1 | 2.3±0.1 |
| LV diameter,d (mm) | 3.5±0.1 | 3.7±0.1 | 3.8±0.1 | 3.7±0.1 |
| LV volume,s (μl) | 19.2±1.7 | 22.5±3.6 | 19.4±1.9 | 19.2±2.4 |
| LV volume,d (μl) | 51.4±3.9 | 57.9±5.6 | 63.0±3.7 | 57.4±2.4 |
| Stroke volume (μl) | 32.2±2.5 | 35.4±2.4 | 43.5±2.2<sup>††</sup> | 38.2±1.6 |
| LVEF (%) | 62.6±1.9 | 62.5±2.5 | 69.5±1.9 | 67.2±3.2 |
| LVFS (%) | 33.1±1.4 | 33.3±1.7 | 38.6±1.6 | 37.0±2.5 |
| Cardiac Output (mL/min) | 12.1±1.3 | 11.7±0.5 | 16.5±0.8<sup>†††</sup> | 13.5±0.4<sup>‡</sup> |
| LV Mass (mg) | 140.7±8.6 | 126.1±8.9 | 178.9±19.1 | 157.7±22.2 |
| LV Mass Corrected (mg) | 112.5±6.9 | 100.8±7.1 | 143.1±15.3 | 126.2±17.8 |
| LVAW, s (mm) | 1.8±0.1 | 1.7±0.1 | 2.0±0.1 | 1.9±0.1 |
| LVAW,d (mm) | 1.3±0.0 | 1.1±0.1 | 1.4±0.1 | 1.3±0.1 |
| LVPW, s (mm) | 1.1±0.0 | 1.0±0.0 | 1.3±0.1 | 1.3±0.2 |
| LVPW,d (mm) | 0.8±0.1 | 0.7±0.0 | 0.8±0.1 | 0.9±0.1 |
| Body weight (g) | 20.4±0.4 | 18.6±0.4<sup>**</sup> | 26.8±0.3<sup>†††</sup> | 24.0±0.4<sup>****</sup> |

Results are presented as mean ± SEM. Significance was calculated for the CV parameters with a 2-way ANOVA followed by Tukey's multiple comparison test. \*\*\**P*<0.001, \*\**P*<0.01, \**P*<0.05 for comparison between *Rubcn*<sup>-/-</sup> vs *Rubcn*<sup>+/+</sup> at the same time-period. †††*P*<0.0001, †††*P*<0.001, ††*P*<0.01 for comparison between a parameter assessed at 2-month-old versus 6-month-old for *Rubcn*<sup>+/+</sup>. ††††*P*<0.0001, †*P*<0.05 for comparison between cardiac parameter assessed at 2-month-old versus 6-month-old for *Rubcn*<sup>-/-</sup>. LV, left ventricle; s, systole; d, diastole; LVEF, left ventricle ejection fraction; LVFS, left ventricle fractional shortening; LVAWs, left ventricular end-systolic anterior wall diameter; LVAWd, left ventricular end-diastolic anterior wall diameter; LVPWs, left ventricular end-systolic posterior wall diameter; LVAWd, left ventricular end-diastolic posterior wall diameter.

**Table S4:** Serum biochemistry for 2-month-old *Rubcn ^-/-^* phenotype in males.

|  | 2-month-old |  |
| --- | --- | --- |
|  | *Rubcn*<sup>+/+</sup><br>n=7 | *Rubcn*<sup>-/-</sup><br>n=4 |
| Albumin (g/dl) | 2.7±0.0 | 2.6±0.1 |
| Alkaline Phosphatase (U/L) | 44.3±7.2 | 35.9±4.0 |
| ALT (U/L) | 33.5±5.1 | 35.1±7.1 |
| Amylase (U/L) | 2,618.1±135.3 | 2,847.8±49.8 |
| Total Bilirubin (mg/dl) | 0.2±0.1 | 0.1±0.0 |
| Blood Urea Nitrogen(mg/dl) | 21.5±1.8 | 15.0±3.0 |
| Cholesterol (mg/dl) | 149.1±13.7 | 177.8±9.6 |
| HDL (mg/dl) | 101.9±9.7 | 117.2±5.3 |
| LDL (mg/dl) | 19.2±2.2 | 23.3±1.2 |
| Calcium (mg/dl) | 6.3±0.1 | 6.0±0.0 |
| CK-MB (U/L) | 289.7±29.0 | 458.3±118.4 |
| Creatinine (mg/dl) | 0.6±0.0 | 0.6±0.1 |
| Glucose (mg/dl) | 159.5±16.2 | 193.5±32.8 |
| Potassium (mmol/L) | 5.4±0.1 | 5.5±0.3 |
| Sodium (mmol/L) | 151.3±0.6 | 148.5±0.9 |
| Phosphorous (mg/dl) | 7.1±0.5 | 6.5±0.1 |
| Total Protein (g/dl) | 4.7±0.1 | 4.5±0.1 |
| Triglyceride (mg/dl) | 24.1±4.8 | 24.9±4.7 |
| Uric Acid (mg/dl) | 1.7±0.1 | 2.4±0.1 |
| Globulin (g/dl) | 2.0±0.1 | 2.0±0.1 |

Results are presented as mean ± SEM. Significance was calculated for the biochemistry parameters with a multiple unpaired t test corrected with Benjamini method for multiple testing.

**Table S5:** Complete blood count for 2-month-old *Rubcn ^-/-^* phenotype in males.

|  | 2-month-old |  |
| --- | --- | --- |
|  | *Rubcn*<sup>+/+</sup><br>n=7 | *Rubcn*<sup>-/-</sup><br>n=4 |
| White blood cells (x 10<sup>3</sup>/μL) | 5.5±0.3 | 5.9±0.5 |
| Neutrophils (count) | 0.9±0.1 | 0.8±0.1 |
| Lymphocytes (count) | 4.4±0.3 | 4.9±0.4 |
| Monocytes (count) | 0.2±0.0 | 0.2±0.0 |
| Eosinophils (count) | 0.0±0.0 | 0.0±0.0 |
| Basophils (count) | 0.0±0.0 | 0.0±0.0 |
| Nucleated red blood cells (count) | 0.0±0.0 | 0.0±0.0 |
| Neutrophils (%) | 16.3±1.3 | 13.1±1.1 |
| Lymphocytes (%) | 79.8±1.2 | 82.8±0.9 |
| Monocytes (%) | 3.7±0.6 | 3.8±0.4 |
| Eosinophils (%) | 0.1±0.0 | 0.2±0.1 |
| Basophils (%) | 0.0±0.0 | 0.1±0.1 |
| Nucleated red blood cells (%) | 0.0±0.0 | 0.0±0.0 |
| Hematocrit (%) | 34.9±0.5 | 34.6±0.4 |
| Red blood cells (x 10<sup>6</sup>/μL) | 8.6±0.1 | 8.5±0.1 |
| Hemoglobin (g/dL) | 12.8±0.2 | 12.7±0.2 |
| Mean corpuscular volume (fL) | 40.8±0.1 | 40.8±0.3 |
| Mean corpuscular hemoglobin (pg) | 14.9±0.2 | 15.0±0.1 |
| Mean corpuscular hemoglobin concentration (g/dL) | 36.5±0.5 | 36.8±0.4 |
| Red cell distribution width (%) | 13.3±0.2 | 13.0±0.1 |
| Red cell distribution width standard deviation (%) | 5.4±0.1 | 5.3±0.1 |
| Reticulocytes (count) | 9.1±5.3 | 6.4±6.4 |
| Reticulocytes (%) | 0.1±0.1 | 0.1±0.1 |
| Platelets (x 10<sup>3</sup>/μL) | 483.3±19.6 | 496.3±28.6 |
| Mean platelets volume (fL) | 4.8±0.0 | 4.8±0.0 |
| Platelets distribution width | 49.9±0.4 | 50.2±0.4 |
| Plateletcrit | 0.2±0.0 | 0.2±0.0 |

Results are presented as mean ± SEM. Significance was calculated for the biochemistry parameters with a multiple unpaired t test corrected with Benjamini method for multiple testing

**Table S6:** CV parameters for *Rubcn^fl/fl^, Cdh5-Cre^Tg+^* phenotype.

|  | 2-month-old |  | 6-month-old |  |
| --- | --- | --- | --- | --- |
|  | *Rubcn<sup>fl/fl</sup>*,<br>*Cdh5-Cre<sup>Tg-</sup>*<br>n=8 | *Rubcn<sup>fl/fl</sup>*,<br>*Cdh5-Cre<sup>Tg+</sup>*<br>n=7 | *Rubcn<sup>fl/fl</sup>*,<br>*Cdh5-Cre<sup>Tg-</sup>*<br>n=8 | *Rubcn<sup>fl/fl</sup>*,<br>*Cdh5-Cre<sup>Tg+</sup>*<br>n=7 |
| Heart Rate (BPM) | 385.3±6.7 | 392.2±3.0 | 393.3±9.8 | 380.1±4.9 |
| LV diameter,s (mm) | 2.4±0.1 | 2.8±0.1 | 2.8±0.1 | 3.1±0.2 |
| LV diameter,d (mm) | 3.6±0.1 | 3.8±0.1 | 4.0±0.1 | 4.3±0.2 |
| LV volume,s (μl) | 21.1±2.8 | 29.9±3.3 | 30.8±3.2 | 40.9±5.4 |
| LV volume,d (μl) | 56.5±5.2 | 63.1±4.1 | 71.2±4.7 | 84.2±6.9<sup>‡</sup> |
| Stroke volume (μl) | 35.5±2.8 | 33.2±1.1 | 40.4±2.6 | 43.3±2.7<sup>‡</sup> |
| LVEF (%) | 63.5±2.5 | 53.5±2.5<sup>*</sup> | 57.4±2.5 | 52.9±3.8 |
| LVFS (%) | 34.0±1.7 | 27.2±1.6<sup>*</sup> | 29.9±1.7 | 27.3±2.5 |
| Cardiac Output (mL/min) | 13.6±1.0 | 13.0±0.5 | 16.0±1.3 | 16.4±0.9<sup>‡</sup> |
| LV Mass (mg) | 119.4±6.6 | 124.8±5.4 | 158.9±10.0<sup>†</sup> | 187.7±27.2<sup>††</sup> |
| LV Mass Corrected (mg) | 95.6±5.2 | 99.8±4.3 | 127.1±8.0<sup>†</sup> | 150.1±21.7<sup>††</sup> |
| LVAW, s (mm) | 1.3±0.1 | 1.1±0.1 | 1.6±0.1<sup>††</sup> | 1.6±0.1<sup>††</sup> |
| LVAW,d (mm) | 1.0±0.0 | 0.9±0.0 | 1.2±0.1<sup>††</sup> | 1.2±0.1<sup>‡</sup> |
| LVPW, s (mm) | 1.2±0.1 | 1.1±0.0 | 1.2±0.1 | 1.2±0.1 |
| LVPW,d (mm) | 0.9±0.1 | 0.8±0.0 | 0.7±0.0 | 0.8±0.0 |
| Body weight (g) | 28.0±0.6 | 29.6±0.3 | 31.8±0.7<sup>†††</sup> | 34.3±0.7<sup>**†††</sup> |

Results are presented as mean ± SEM. Significance was calculated for the CV parameters with a 2-way ANOVA followed by Tukey's multiple comparison test. <sup>\*\*</sup>*P*<0.01, <sup>\*</sup>*P*<0.05 for comparison between *Rubcn<sup>fl/fl</sup>*, *Cdh5-Cre<sup>Tg+</sup>* vs *Rubcn<sup>fl/fl</sup>*, *Cdh5-Cre<sup>Tg-</sup>* at the same time-period. <sup>†††</sup>*P*<0.0001, <sup>††</sup>*P*<0.01, <sup>†</sup>*P*<0.05 for comparison between a parameter assessed at 2-month-old versus 6-month-old for *Rubcn<sup>fl/fl</sup>*, *Cdh5-Cre<sup>Tg-</sup>*. <sup>††††</sup>*P*<0.0001, <sup>††</sup>*P*<0.01, <sup>†</sup>*P*<0.05 for comparison between cardiac parameter assessed at 2-month-old versus 6-month-old for *Rubcn<sup>fl/fl</sup>*, *Cdh5-Cre<sup>Tg+</sup>*. LV, left ventricle; s, systole; d, diastole; LVEF, left ventricle ejection fraction; LVFS, left ventricle fractional shortening; LVAWs, left ventricular end-systolic anterior wall diameter; LVAWd, left ventricular end-diastolic anterior wall diameter; LVPWs, left ventricular end-systolic posterior wall diameter; LVAWd, left ventricular end-diastolic posterior wall diameter.

**Table S7:**
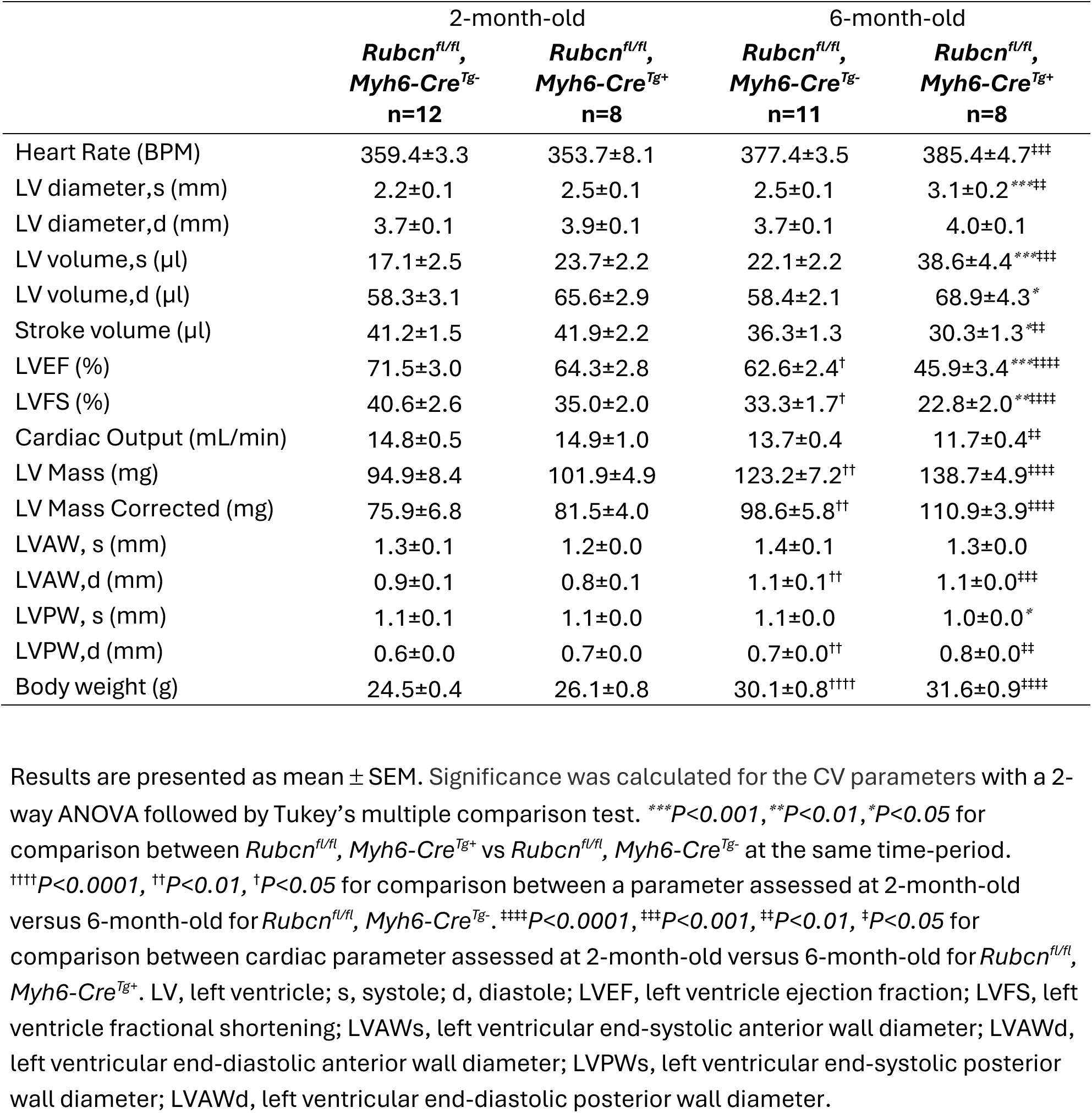
CV parameters for *Rubcn^fl/fl^, Myh6-Cre^Tg+^* phenotype.

|  | 2-month-old |  | 6-month-old |  |
| --- | --- | --- | --- | --- |
|  | *Rubcn<sup>fl/fl</sup>*,<br>*Myh6-Cre<sup>Tg-</sup>*<br>n=12 | *Rubcn<sup>fl/fl</sup>*,<br>*Myh6-Cre<sup>Tg+</sup>*<br>n=8 | *Rubcn<sup>fl/fl</sup>*,<br>*Myh6-Cre<sup>Tg-</sup>*<br>n=11 | *Rubcn<sup>fl/fl</sup>*,<br>*Myh6-Cre<sup>Tg+</sup>*<br>n=8 |
| Heart Rate (BPM) | 359.4±3.3 | 353.7±8.1 | 377.4±3.5 | 385.4±4.7<sup>††</sup> |
| LV diameter,s (mm) | 2.2±0.1 | 2.5±0.1 | 2.5±0.1 | 3.1±0.2<sup>***††</sup> |
| LV diameter,d (mm) | 3.7±0.1 | 3.9±0.1 | 3.7±0.1 | 4.0±0.1 |
| LV volume,s (μl) | 17.1±2.5 | 23.7±2.2 | 22.1±2.2 | 38.6±4.4<sup>***†††</sup> |
| LV volume,d (μl) | 58.3±3.1 | 65.6±2.9 | 58.4±2.1 | 68.9±4.3<sup>*</sup> |
| Stroke volume (μl) | 41.2±1.5 | 41.9±2.2 | 36.3±1.3 | 30.3±1.3<sup>††</sup> |
| LVEF (%) | 71.5±3.0 | 64.3±2.8 | 62.6±2.4<sup>†</sup> | 45.9±3.4<sup>***†††</sup> |
| LVFS (%) | 40.6±2.6 | 35.0±2.0 | 33.3±1.7<sup>†</sup> | 22.8±2.0<sup>***†††</sup> |
| Cardiac Output (mL/min) | 14.8±0.5 | 14.9±1.0 | 13.7±0.4 | 11.7±0.4<sup>††</sup> |
| LV Mass (mg) | 94.9±8.4 | 101.9±4.9 | 123.2±7.2<sup>††</sup> | 138.7±4.9<sup>†††</sup> |
| LV Mass Corrected (mg) | 75.9±6.8 | 81.5±4.0 | 98.6±5.8<sup>††</sup> | 110.9±3.9<sup>†††</sup> |
| LVAW, s (mm) | 1.3±0.1 | 1.2±0.0 | 1.4±0.1 | 1.3±0.0 |
| LVAW,d (mm) | 0.9±0.1 | 0.8±0.1 | 1.1±0.1<sup>††</sup> | 1.1±0.0<sup>††</sup> |
| LVPW, s (mm) | 1.1±0.1 | 1.1±0.0 | 1.1±0.0 | 1.0±0.0<sup>*</sup> |
| LVPW,d (mm) | 0.6±0.0 | 0.7±0.0 | 0.7±0.0<sup>††</sup> | 0.8±0.0<sup>††</sup> |
| Body weight (g) | 24.5±0.4 | 26.1±0.8 | 30.1±0.8<sup>†††</sup> | 31.6±0.9<sup>†††</sup> |

Results are presented as mean ± SEM. Significance was calculated for the CV parameters with a 2-way ANOVA followed by Tukey's multiple comparison test. <sup>\*\*\*</sup> $P < 0.001$ , <sup>\*\*</sup> $P < 0.01$ , <sup>\*</sup> $P < 0.05$ for comparison between *Rubcn<sup>fl/fl</sup>*, *Myh6-Cre<sup>Tg+</sup>* vs *Rubcn<sup>fl/fl</sup>*, *Myh6-Cre<sup>Tg-</sup>* at the same time-period. <sup>††††</sup> $P < 0.0001$ , <sup>††</sup> $P < 0.01$ , <sup>†</sup> $P < 0.05$ for comparison between a parameter assessed at 2-month-old versus 6-month-old for *Rubcn<sup>fl/fl</sup>*, *Myh6-Cre<sup>Tg-</sup>*. <sup>††††</sup> $P < 0.0001$ , <sup>†††</sup> $P < 0.001$ , <sup>††</sup> $P < 0.01$ , <sup>†</sup> $P < 0.05$ for comparison between cardiac parameter assessed at 2-month-old versus 6-month-old for *Rubcn<sup>fl/fl</sup>*, *Myh6-Cre<sup>Tg+</sup>*. LV, left ventricle; s, systole; d, diastole; LVEF, left ventricle ejection fraction; LVFS, left ventricle fractional shortening; LVAWs, left ventricular end-systolic anterior wall diameter; LVAWd, left ventricular end-diastolic anterior wall diameter; LVPWs, left ventricular end-systolic posterior wall diameter; LVAWd, left ventricular end-diastolic posterior wall diameter.

**Table S8:** CV parameters for *Rubcn^fl/fl^, LysM-Cre^Tg+^* phenotype.

|  | 2-month-old |  | 6-month-old |  |
| --- | --- | --- | --- | --- |
|  | *Rubcn<sup>fl/fl</sup>*,<br>*LysM-Cre<sup>Tg-</sup>*<br>n=11 | *Rubcn<sup>fl/fl</sup>*,<br>*LysM-Cre<sup>Tg+</sup>*<br>n=9 | *Rubcn<sup>fl/fl</sup>*,<br>*LysM-Cre<sup>Tg-</sup>*<br>n=11 | *Rubcn<sup>fl/fl</sup>*,<br>*LysM-Cre<sup>Tg+</sup>*<br>n=9 |
| Heart Rate (BPM) | 408.0±9.0 | 381.5±13.4* | 414.3±2.9 | 414.2±2.8<sup>††</sup> |
| LV diameter,s (mm) | 2.0±0.2 | 1.9±0.2 | 2.3±0.2 | 2.1±0.1 |
| LV diameter,d (mm) | 3.2±0.1 | 3.2±0.1 | 3.7±0.1<sup>††</sup> | 3.5±0.1<sup>†</sup> |
| LV volume,s (μl) | 14.8±3.1 | 13.1±2.5 | 19.5±3.1 | 15.3±2.7 |
| LV volume,d (μl) | 42.5±3.8 | 40.6±3.7 | 59.7±3.6<sup>††</sup> | 51.4±4.2<sup>†</sup> |
| Stroke volume (μl) | 27.7±1.5 | 27.5±2.0 | 40.2±2.2<sup>†††</sup> | 36.1±2.4<sup>††</sup> |
| LVEF (%) | 68.1±3.9 | 69.4±4.4 | 68.7±4.0 | 71.6±3.1 |
| LVFS (%) | 37.8±2.9 | 39.3±3.9 | 38.8±3.0 | 40.5±2.5 |
| Cardiac Output (mL/min) | 11.3±0.7 | 10.6±0.9 | 16.7±0.9<sup>†††</sup> | 15.0±1.0<sup>††</sup> |
| LV Mass (mg) | 84.3±5.2 | 81.8±5.6 | 92.3±4.3 | 87.0±5.8 |
| LV Mass Corrected (mg) | 67.5±4.1 | 65.4±4.5 | 73.8±3.5 | 69.6±4.6 |
| LVAW, s (mm) | 1.0±0.1 | 1.2±0.1 | 0.9±0.0 | 0.8±0.0<sup>†††</sup> |
| LVAW,d (mm) | 0.7±0.1 | 0.8±0.0 | 0.6±0.0<sup>†</sup> | 0.6±0.0<sup>†††</sup> |
| LVPW, s (mm) | 1.3±0.1 | 1.2±0.1 | 1.3±0.0 | 1.5±0.1 |
| LVPW,d (mm) | 0.9±0.0 | 0.8±0.0 | 0.8±0.0 | 0.9±0.0 |
| Body weight (g) | 20.8±0.7 | 20.8±1.5 | 29.7±0.6<sup>††††</sup> | 29.4±0.7<sup>†††</sup> |

Results are presented as mean ± SEM. Significance was calculated for the CV parameters with a 2-way ANOVA followed by Tukey's multiple comparison test. \* $P<0.05$ for comparison between *Rubcn<sup>fl/fl</sup>*, *LysM-Cre<sup>Tg+</sup>* vs *Rubcn<sup>fl/fl</sup>*, *LysM-Cre<sup>Tg-</sup>* at the same time-period. <sup>††††</sup> $P<0.0001$ , <sup>†††</sup> $P<0.001$ , <sup>††</sup> $P<0.01$ , <sup>†</sup> $P<0.05$ for comparison between a parameter assessed at 2-month-old versus 6-month-old for *Rubcn<sup>fl/fl</sup>*, *LysM-Cre<sup>Tg-</sup>*. <sup>††††</sup> $P<0.0001$ , <sup>†††</sup> $P<0.001$ , <sup>††</sup> $P<0.01$ , <sup>†</sup> $P<0.05$ for comparison between cardiac parameter assessed at 2-month-old versus 6-month-old for *Rubcn<sup>fl/fl</sup>*, *LysM-Cre<sup>Tg+</sup>*. LV, left ventricle; s, systole; d, diastole; LVEF, left ventricle ejection fraction; LVFS, left ventricle fractional shortening; LVAWs, left ventricular end-systolic anterior wall diameter; LVAWd, left ventricular end-diastolic anterior wall diameter; LVPWs, left ventricular end-systolic posterior wall diameter; LVAWd, left ventricular end-diastolic posterior wall diameter.

**Table S9:** Three main sources of QTL used for genomic analysis.

| QTL | TISSUES | N SAMPLES | PUBLICATION | PMID | MINOR ALLELE FREQUENCY FILTER | TOTAL GENES OR PROTEINS | GENOTYPING | p-VALUE FILTER |
| --- | --- | --- | --- | --- | --- | --- | --- | --- |
| eQTL | whole-blood | 31 684 blood samples | Vosa et al. NatGenetics 2021 | 34475573 | MAF>0.01 | 19960 genes | different platforms (23 different studies) | $P < 2.02 \times 10^{-5}$ |
| eQTL | 10 selected tissues | 838 samples for GTEX V8 | GTEx consortium Science 2020 | 32913098 | MAF>0.01 | 23268 genes | whole-genome sequencing | FDR<0.05 |
| pQTL | Bran (Cortex) | 330 postmortem dorsolateral prefrontal cortex | Robins et al. AJHG 2021 | 33571421 | MAF>0.05 | 7376 proteins | whole-genome sequencing on PBMC or frozen cortex | FDR<0.05 |

**Table S10:**
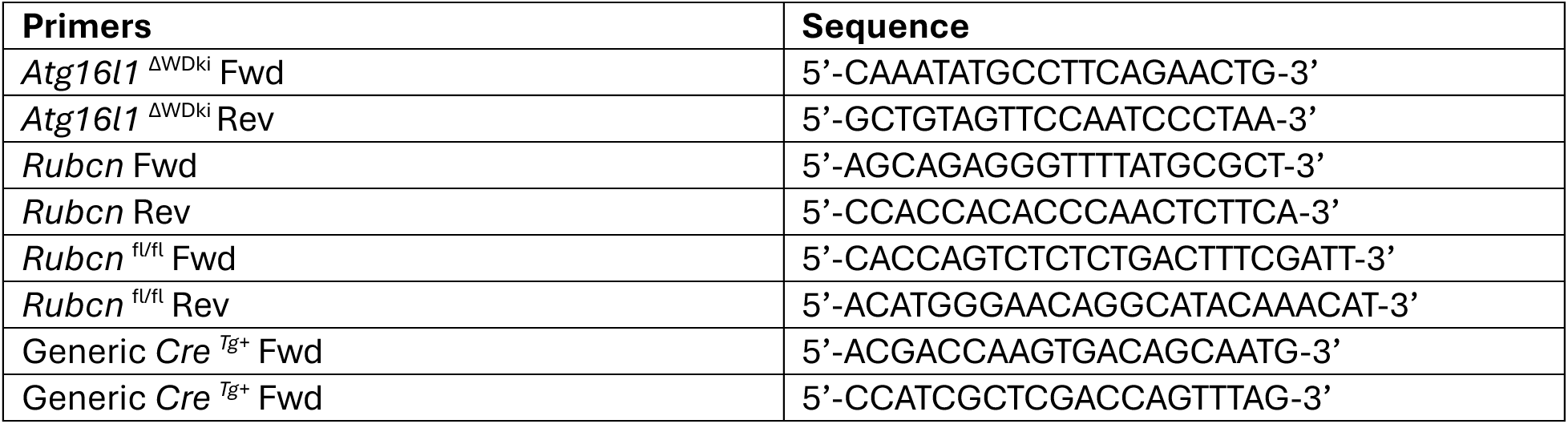
Oligonucleotide primers sequences for genotyping.

**Table S11:** List of antibodies used.

| **Antibody** | **Dilution** | **Company** | **Catalog number** |
| --- | --- | --- | --- |
| ***Western blot*** |  |  |  |
| Anti-Rubicon | 1:500 | Cell signaling Technology | 8465 |
| Anti-ATG16L1 | 1:500 | MBL | PM040 |
| Anti-total VEGFR2 | 1:1000 | Cell signaling Technology | 2479 |
| Anti-Y1175 phospho VEGFR2 | 1:400 | Cell signaling Technology | 2478 |
| Anti-total eNOS | 1:500 | Cell signaling Technology | 9571 |
| Anti-human Ser1177 phospho eNOS | 1:400 | Cell signaling Technology | 9572 |
| Anti-mouse Ser1177 phospho eNOS | 1:500 | BD | 612392 |
| Anti-CXCR4 | 1:1000 | R and D | MAB21651-100 |
| Anti-PECAM1 | 1:1000 | Cell signaling Technology | 77699 |
| Anti-Hemoglobin alpha | 1:300 | Abcam | Ab174536 |
| Anti-AHSP | 1:500 | Abcam | Ab174536 |
| Anti-VPS29 | 1:1000 | Cell signaling Technology | 73540 |
| Anti-ATG3 | 1:1000 | MBL | M133-3 |
| Anti-PIK3C2A | 1:1000 | Proteintech | 22028-I-AP |
| Anti-Actin | 1:20000 | Santa-Cruz | SC-47778-HRP |
| ***Recycling assay*** |  |  |  |
| Anti-VEGFR2 | 1:200 | R and D | MAB3573 |
| Anti-VE cadherin | 1:200 | Sigma | MABT134 |
| Anti-CD31 | 1:1000 | ThermoScientific | MA5-13188 |
| Anti-Piezo-1 | 1:100 | Proteintech | 82625-4-RR |
| Goat Anti-mouse-AF Plus 647 | 1:500 | ThermoScientific | A48289 |
| Goat Anti-Rabbit-AF Plus 647 | 1:500 | ThermoScientific | A48285 |
| IgG1 isotype control | 1:10000 | BioXCell | BE0083 |
| IgG2 control | 1:10000 | BioXCell | BE0301 |
| Hoechst 33342 | 1:20000 | Thermo | H1399 |
| ***Immunofluorescence*** |  |  |  |
| Anti-Hemoglobin alpha | 1:100 | Abcam | Ab174536 |
| Hoechst 33342 | 1:20000 | Biolegend | 422801 |
| Goat Anti-Rabbit-AF Plus 647 | 1:500 | ThermoScientific | A32733 |
| CellMask™ Green Actin Stain | 1:5000 | ThermoScientific | A57243 |

